# The distribution, phenology, host range and pathogen prevalence of *Ixodes ricinus* in France: a systematic map and narrative review

**DOI:** 10.1101/2023.04.18.537315

**Authors:** Grégoire Perez, Laure Bournez, Nathalie Boulanger, Johanna Fite, Barbara Livoreil, Karen D. McCoy, Elsa Quillery, Magalie René-Martellet, Sarah I. Bonnet

## Abstract

The tick *Ixodes ricinus* is the most important vector species of infectious diseases in European France. Understanding its distribution, phenology, and host species use, along with the distribution and prevalence of associated pathogens at national scales is essential for developing prevention strategies. The aim of this paper is to provide a systematic map and narrative review of the existing knowledge on the eco-epidemiology of *I*. *ricinus* in France. Using literature published up to 2020, the present paper provides a distribution map for the species and a summary of environmental factors explaining observed geographical differences in phenology and temporal differences in abundance. The diversity of vertebrate host species used by this tick, along with their degree of infestation when available, are presented and discussed with respect to their potential contribution to the population dynamics of *I*. *ricinus* and the circulation of tick-borne zoonotic pathogens. Prevalence data of detected pathogens are summarised in different maps. Results from 187 identified references show that the species is present in most departments, but scarce under Mediterranean climate and in coastal habitats. Its phenology is generally bimodal with variations depending on climate. Abundance seems positively influenced by forest cover and host abundance. Rodents and ruminants are the most studied species groups, but the diversity of sampling protocols (e.g., location, season, exhaustivity of inspection) precluded direct comparisons between species groups. Data on pathogens are patchy, with most studies conducted near research units. Among pathogens, *Borrelia burgdorferi* sensu lato is the most searched for in ticks and seems more prevalent in north-eastern and central France. The review carried out here has made it possible to highlight the gaps in our knowledge of tick-host-pathogen interactions, their ecology and their distribution, and the need to address these gaps in order to optimize tick and tick-borne diseases prevention and control strategies.

## 1 Introduction

Among the forty one tick species known to be present in European (metropolitan) France (i.e., excluding the ultra-marine French territories, France hereafter), the hard tick *Ixodes ricinus* is the most frequently involved in human tick bites (Gilot & Marjolet 1982; Pérez-Eid 2007). This tick transmits several pathogens responsible for human and animal diseases (e.g. Lyme borreliosis, tick-borne encephalitis, piroplasmosis, anaplasmosis) (Heyman et al. 2010). Both the biology and ecology of this European tick species have been recently reviewed by Gray et al. (2021), and are known to vary greatly among regions. Data collected at national levels can provide insight into the factors responsible for at least some of this variation, but are rarely available to the scientific community because of a mix of published and unpublished work and its production in different languages. Here, we synthesize the available data on *I. ricinus* and its associated pathogens in France up to 2020.

Some aspects concerning the life cycle of *I. ricinus* are well-known. For example, like most hard ticks (*Ixodidae*), each life stage of *I*. *ricinus* (larva, nymph and adult) takes only a single blood meal, except adult males which do not need to feed, but may take a limited amount of blood. After each blood meal, *I*. *ricinus* drops off to the ground to moult into the next stage or, in the case of adult females, to lay eggs and die. Feeding on a host can last three to six days for larvae, four to seven days for nymphs and seven to ten days for adult females. Other aspects of the life cycle are more geographically variable. Indeed, the *I*. *ricinus* life cycle can take anywhere from two to four years to complete, depending on environmental conditions and the length of diapause periods (Gray et al. 2021). The questing activity of this tick is regulated by photoperiod and is constrained to temperatures between 5 and 30°C and to a minimum relative humidity of 80% (Gray et al. 2021). Nevertheless, despite these generalities, local adaptation of *I*. *ricinus* populations is known to occur (Gray et al. 2021; Tomkins et al. 2014). The life cycle of *I. ricinus* is triphasic, that is, each stage feeds on a different individual host. It is a generalist (or telotrope) tick in terms of its host range. The composition and abundance of the local vertebrate host population directly influences tick infestation dynamics, relative abundance and dispersal capacities. In turn, this local vertebrate community will affect the transmission dynamics of tick-borne pathogens and their relative prevalence in questing ticks (Hofmeester et al. 2016). Host choice varies according to tick life stage and can be at least partly explained by the height at which *I*. *ricinus* ticks quest on the vegetation, which itself depends on their relative resistance to desiccation (lowest in larvae and highest in adults), the type of vegetation, and the microclimatic conditions (Mejlon & Jaensen 1997; Randolph & Storey 1999). For example, due to the presence of larval ticks within the leaf litter (where egg laying takes place), small mammals (wood mice, voles and shrews) are often considered as the primary hosts of larvae, followed by passerine birds living in closed habitats (e.g., forest or hedgerows) and feeding on the ground (e.g., blackbirds, thrushes, tits, robin) (Hofmeester et al. 2016). Nymphs are frequently found on birds, lizards, and medium to large mammals, while adult females, which often quest higher on the vegetation, are generally observed only on medium to large mammals (Hofmeester et al. 2016; Mendoza-Roldana & Colella 2019). Indeed, the distribution and abundance of wild ungulates are considered key factors determining the probability of tick exposure (Takumi et al. 2019). Birds can contribute to the dissemination of ticks in the landscape at short, medium and large scales (Vollmer et al. 2011; Marsot et al. 2012; Hofmeester et al. 2016). The high variation in the population density of small mammals, itself influenced by tree seed production (masting), is also thought to be an important driver of tick population dynamics at local scales (Perez et al. 2016; Bregnard, Rais & Voordouw 2020). Because of their differential susceptibility to environmental conditions and to variation in ontological processes, the different tick life stages also differ in their seasonal activity periods, with larvae generally appearing later in the season than nymphs and adults (Wongnak et al. 2022). Identifying the main hosts and the timing of tick activity according to the life stage are essential elements for understanding the transmission dynamics of tick-borne pathogens and thus for implementing local prevention measures (Randoph et al. 1999).

France covers 552,000 km² in Europe, extending through about 9° of latitude and including a large panel of climatic zones (semi-continental, Mediterranean, oceanic and mountainous zones) and a rich diversity of habitats therein. Climates and environments suitable for the development of *I*. *ricinus* cover a large part of the territory, thus explaining the broad distribution and major importance of this tick species for both human and veterinary health (Wongnak et al. 2022). Consequently, several studies have been conducted on *I. ricinus* and its associated pathogens in France, but in a piecemeal way, focusing for example on a particular tick-borne disease (e.g. Lyme borreliosis, babesiosis), a particular host group (e.g. cattle, small mammals), or a particular geographic area. The present work represents the first attempt to synthesize available data on *I. ricinus* and the pathogens it transmits in France. On the basis of a systematic literature search, we synthesize knowledge on:

1. Host infestation by *I*. *ricinus*, to detect potential differences in host use within France and among areas in Europe.
2. The spatio-temporal distribution of *I*. *ricinus* in France, to identify possible geographic trends in tick abundance, variation in activity patterns and the influence of environmental factors.
3. Tick-borne pathogens detected in *I*. *ricinus*, to better assess their spatial distribution and variation in exposure risk.

Finally, by highlighting knowledge gaps in the ecology of *I. ricinus* and acarological risk in France, we aim to orient future research and preventive measures in this country, and to provide a comparable basis for studies in other regions of *I. ricinus’* distribution.

## 2 Materials and methods

### 2.1 Research approach

To retrieve a comprehensive set of available literature on the ecology of *I*. *ricinus* in France and the pathogens it transmits, we chose a “**Population**, **Context**, **Comparators** and **Outcome”** (PCCO) approach (Livoreil et al. 2017). This approach aims (1) to determine the words to use in the literature search and (2) to structure the criteria for including or excluding a reference. Studied populations, contexts, comparators and outcome variables are presented in **Table 1**.

**Table 1:** Components elements of the literature search.

| Component | Elements |
| --- | --- |
| Population | Any *I. ricinus* populations at larval, nymphal or adult stage |
| Context | European France, i.e. excluding overseas French territories |
| Comparators | No comparator were excluded. Comparators include, for instance: temporal comparisons, i.e. intra-an-<br>nual, weather-related and daily; spatial comparisons according to climate (biome), land cover and land<br>use (landscape context and management, habitats, vegetation types, anthropic activities, management);<br>host density and diversity (wildlife management, ex-closure or enclosure, population fluctuation); poten-<br>tial treatments (e.g. acaricide treatments, vaccinations) |
| Outcome variables | Presence/absence; abundance; density; hosts parasitic burden (presence, prevalence or mean infesta-<br>tion); distribution and dispersal (displacements at different scales, gene flows); behaviour (e.g. questing<br>activity, host preference); search for tick-borne pathogens (presence and/or prevalence) |

The target “**populations”** were *I*. *ricinus* at all life stages: larvae, nymph and adult. The “**context”** corresponded to European France as the target area (French territories outside the European continent were excluded because they are outside the distributional range of *I*. *ricinus*) and included the environmental factors of interest identified from both short-term punctual studies and long-term monitoring and from micro-habitat to bio-regional scales. The “**comparator”** corresponds to the kinds of comparisons that could be made with the references. All types of comparators were eligible for the review *a priori*. Expected comparators were spatio-temporal comparisons, including inter-annual (i.e., dynamics), intra-annual (i.e., seasonal variation in questing tick density), and daily comparisons (i.e., abundance according to the hour of the day). Spatial comparisons occurred at multiple scales: micro-habitats, habitats, landscapes and the biogeographic scale. It also included comparisons between vegetation management types, levels of anthropic pressure and/or types of anthropic activities, vertebrate host densities and physical, chemical or biological treatments of the environment or the host. The “**outcome”** variables of interest were those providing information on the ecology of *I. ricinus*: relative abundance in space and time, survival, activity, host use and infestation intensity by life stage, distribution and dispersal. The assessment of tick-borne pathogens in ticks was also considered as an outcome variable.

### 2.2 Search strategy

#### 2.2.1 Keywords and search lines

The literature search was conducted on titles, abstracts and keywords in English, but references in both English and French were considered. The search line included the name of the species of interest, “*Ixodes ricinus*” and all the outcome variables identified in **Table 1**. The keyword “France” was not included in the search line because the sampling country is not always mentioned in the title or abstract. The keywords “pathogen” and “infectious agents” were not included to avoid references focused only on cellular or molecular aspects of tick-borne pathogens in *I*. *ricinus*, assuming that the detection of pathogens would have been expressed as presence or prevalence. The final search line was:

> **“Ixodes ricinus”** AND **(abundance** OR **activity** OR **[behaviour** OR **behavior]** OR **burden** OR **density** OR **dispersal** OR **distribution** OR **dynamics** OR **infestation** OR **presence** OR **prevalence** OR **questing)**.

This search line was adapted to correspond to the different database constraints (see exact search lines in **SI 1**).

#### 2.2.2 Databases

The search was conducted between November 14^th^ and December 16^th^ 2019 in the following databases: CAB Direct, JSTOR, Pascal & Francis, PubMed, ScienceDirect, Scopus, WorldCat (see **SI 1** for further details). To access French publications and grey literature, a complementary search was conducted over the same period using “*Ixodes ricinus*” as the unique keyword in the following databases: Agricola, Open Access Thesis and Dissertations, Thèses vétérinaires – i.e. Doctor in Veterinary Medicine (DVM) dissertation theses – (French only), and theses.fr – i.e. PhD theses – (French only). The efficiency of the search strategy was tested using a previously established test-list of 110 references known to be relevant for at least one of the established criterion references (see **SI 2**). To reach a more exhaustive corpus, additional references identified opportunistically and published until 2020 were also considered and submitted to the same eligibility screening (see below).

### 2.3 Eligibility screening

#### 2.3.1 Eligibility criteria

We retained peer-reviewed research articles and reviews (not including conference proceedings), book sections, veterinary or university theses published at any time prior to December 16^th^, 2019. Only references in French or English were considered. Screening was conducted by one of the authors (GP), first on titles, then on abstracts, and finally on the full text when available. Finally, studies were included in this systematic review when they met all of the following criteria (see detailed criteria in table **SI 3**):

-at the title and abstract stage:

**Population**: The reference title must mention *I*. *ricinus* or ticks or any word suggesting that *I*. *ricinus* was studied in the paper (e.g., ectoparasites, tick, vector-borne disease). In the latter case, references were retained for subsequent examination (at full text stage).

**Context**: Studies must be performed at least in part in France. If the country of study was not mentioned, the reference was retained for subsequent examination (at full text stage).

**Comparator**: no type of comparator was excluded.

**Outcome**: Studies must be focused directly on the ecology of *I*. *ricinus*. For example, those that focused on cellular or molecular aspects of *I. ricinus* biology or on tick-borne pathogens alone, were excluded. If there was a doubt, the reference was retained for subsequent examination of the full text.

-at full text stage:

**Population:** only articles concerning *I. ricinus* were retained, i.e., with at least one mention of the species.

**Context**: all references including at least one sample from France, Corsica island or other French islands of the Mediterranean Sea, Bay of Biscay, Celtic Sea, English Channel, and North Sea were retained.

**Comparator**: no type of comparator was excluded.

**Outcomes**: the same criteria used for the title and abstract stage were applied.

### 2.4 Systematic map and narrative synthesis

#### 2.4.1 Database for the systematic map

A dataset was created, composed of 17 bibliographic fields and 41 descriptive fields. The 41 descriptive fields were filled mainly on the basis of “Materials and methods” and “Results” sections, annexes and supplementary materials of the retained references. These fields described the study type (2 fields), the sampling strategy in time (4 fields) and space (10 fields), the sampling methods (17 fields), the pathogens studied (4 fields) and the explanatory variables (2 fields). A field was added to justify the potential exclusion of the reference, and another field for comments. The dataset, with a detailed fields description, is available in **SI 4**.

#### 2.4.2 Narrative synthesis

The narrative synthesis presented the type of collected data, the spatial and temporal scales studied, as well as the abiotic and biotic variables associated with the presence (/absence), abundance and activity of *I*. *ricinus.* The vertebrate hosts studied and their mean infestation rates by different *I*. *ricinus* life stages were retrieved from the literature. These data were discussed with respect to existing knowledge on the host species use in Europe. Tick-borne pathogen prevalence per tick life stage was summarised, along with associated detection methods, and distribution maps were established. The collected data on pathogen detection were then discussed taking into account the vector competence of *I*. *ricinus* and the potential reservoir role of the different vertebrate hosts.

## 3 Results and discussion

### 3.1 Reference search results and bibliographic description

#### 3.1.1 Reference search and screening

The systematic search retrieved 19,654 references, as presented in the PRISMA flow chart (**Figure 1**). This included 3,091 references from CAB Direct, 858 from JSTOR, 662 from Pascal & Francis, 2,102 from PubMed, 2,731 from ScienceDirect, 2,473 from Scopus, and 6,174 from WorldCat databases. The complementary search for grey literature in other databases allowed us to identify 1,316 references from Agricola, 150 references from Open Access Thesis and Dissertations, 40 French DVM dissertation theses, and 57 French PhD theses. Only five of the 110 established references from the test-list were not found during our systematic literature search (i.e., 95.4% of references were found). Additionally, 15 references found opportunistically, including five between 2019 and 2020, and fulfilled the inclusion criteria.

After having discarded 11,298 duplicates (57.4%), 8,370 references underwent title and abstract screening and 198 underwent full-text screening for eligibility (**Figure 1**). The most common reasons for exclusion at the title and abstract stage were that sampling was not conducted in France or studies focused on cellular or molecular aspects of *I*. *ricinus* biology. The most common reasons for exclusion at the full text stage were that studies focused on tick-borne pathogen without reference to the vector (laboratory studies or interactions with the vertebrate host only) (7 cases), or were laboratory-based studies of *I*. *ricinus* vector competence. In the end, 187 references fulfilled the inclusion criteria (**SI 5**).

**Figure 1:**
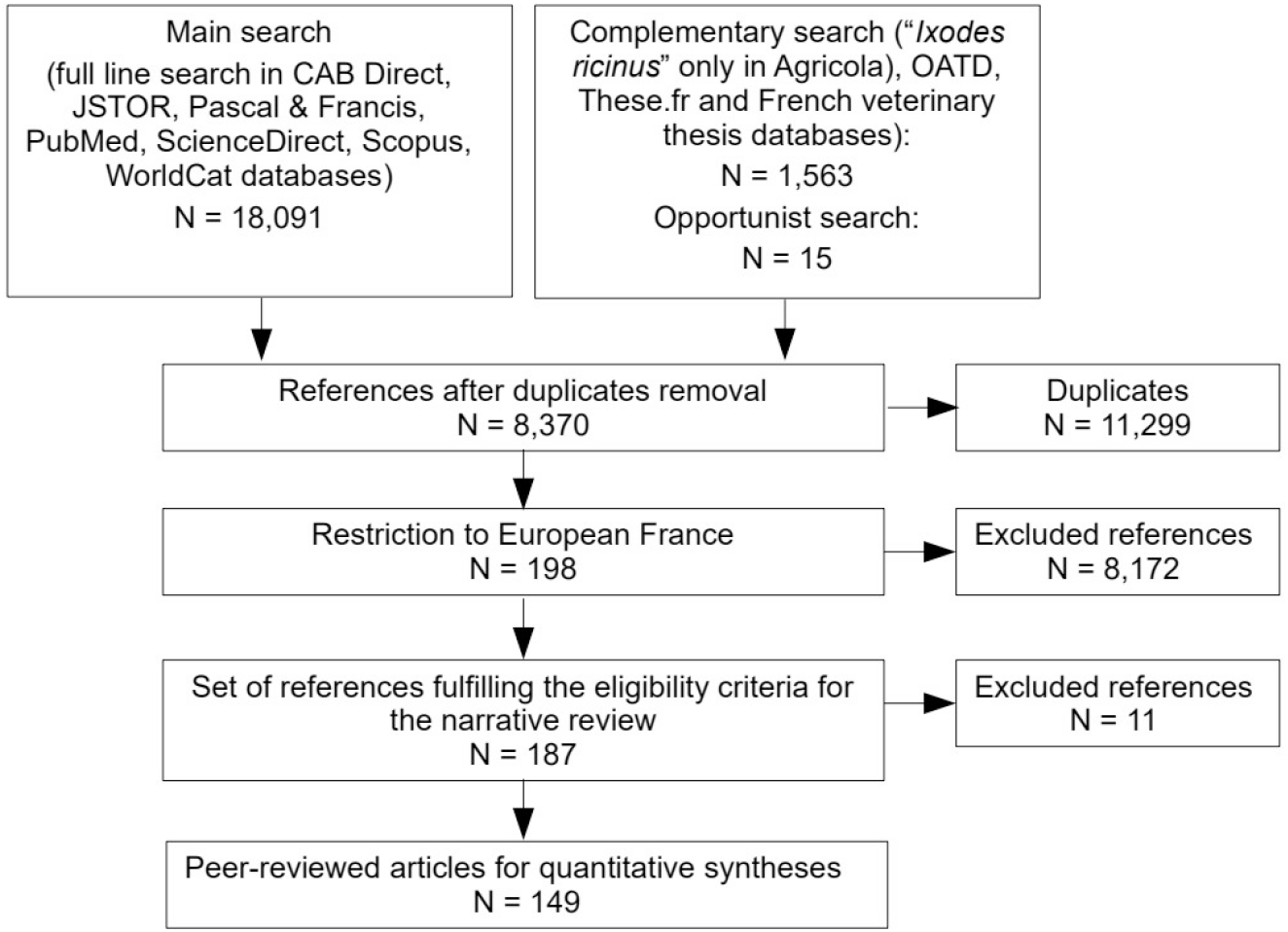
PRISMA-style scheme reporting the literature search and selection strategy arriving to the final 1 87 references including the 149 peer-reviewed articles used for data compilation.

#### 3.1.2 Description of eligible references

Most of the 187 references were peer-reviewed journal articles and included primary research articles (n = 137, 73.3%) and reviews (n = 12, 6.4%), whereas other documents were DVM and Doctor of Pharmacy (PharmD) dissertations (n = 22, 11.8%), PhD theses (n = 15, 8.0%), or book sections (n = 1, 0.5%). Two-thirds of the journal articles were in English (n = 97, 65.1%), while others were in French (n = 52, 34.9%). The oldest reference recorded was published in 1965. References were scarce until the 1990’s, slowed-down again in the 2000’s and then increased again from 2010 onwards (**SI 6**). The 12 selected reviews provided only a broad overview of ticks and tick-borne diseases or, conversely, a synthesis on some precise aspect of *I*. *ricinus* ecology and tick-borne diseases in France; none contained a systematic map providing a summary of existing data for the species.

Of the peer-reviewed articles, most were based on field data followed by laboratory analyses (e.g., searching for pathogens) (n = 130, 87.2%), but some used field-collected ticks to conduct further laboratory experiments (e.g., assessing the effect of a variable on tick activity) (n = 3, 2.2%) or laboratory methodological evaluation (n = 4, 8.1%). Field studies were mainly exploratory (i.e., considering at least one explanatory variable; n = 68, 45.6%) and case/occurrence reports (i.e., only descriptive; n = 60, 40.3%), but two of them (1.5%) were methodological evaluations of sampling techniques. Among studies followed by experiments, one (0.7%) was conducted to assess the reservoir competence of Siberian chipmunks, *Tamia sibiricus barberi*, for *Borrelia* spp., and two (1.3%) combined field data and laboratory experiments to study tick adaptation to climate. Among laboratory methodological evaluations, three (2.2%) were associated with the development of laboratory methods for pathogen detection, and one (0.7%) was involved the development of a method to study tick population genetics based on field-sampled ticks.

### 3.2 Descriptive synthesis of the ecology of *Ixodes ricinus* in France

#### 3.2.1 Distribution of *Ixodes ricinus*

Over the past fifty years, several studies were conducted to assess the distribution of *I*. *ricinus* in different regions of France. The presence of *I*. *ricinus* was reported when ticks were questing on vegetation using the dragging or flagging method (n = 98), attached on a host (n = 59), or both (n = 25). The sampling locations at the department level could be identified in 127 articles. Thus, we used this spatial scale (NUTS-3) to depict the known distribution of the tick species. A distribution map by department was produced from the retrieved data (**Figure 2** and **SI 7**). Sampling effort varied greatly across the territory, with most studies concentrated on the departments of Ille-et-Vilaine (n = 23, Brittany, western), Bas-Rhin (n = 18, Alsace, north-eastern), Essonne (n = 16, Paris region), Loire-Atlantique (n = 15, Pays-de-la-Loire, western), Haut-Rhin (n = 15, Alsace, north-eastern), Puy-de-Dôme (n = 13, Auvergne, central), and Sarthe (n = 12, Pays-de-la-Loire, western). Studies covered 90 out of 96 European French departments. Among these 90 departments, the presence of *I*. *ricinus* was confirmed in 87 (97.7%).

An inventory in the Rhône Valley (south-eastern France) of a 330×80 km area, from Mediterranean sea and northward, recorded the presence of *I. ricinus* in the northern part of the study area, but not the southernmost part (Gilot et al. 1989). In south-western France, in a study that aimed at detecting the presence of *I*. *ricinus* on 100 sites over 15 different vegetation types covering 23 departments, the species was never detected in sites with Mediterranean vegetation (Doche et al. 1993; Gilot et al. 1995). However, in Corsica, Grech-Angelini et al. (2016) reported the species feeding on cattle in areas above 600 m above sea level (a.s.l.) and recently Sevestre et al. (2021) reported the species on hosts and questing in south-eastern France (Alpes-Maritimes) between 450 m and up to 1300 m a.s.l.. Thus, *I. ricinus* ticks seem to be present in some Mediterranean departments, but only under specific environmental conditions and above a certain altitude (Stachurski & Vial 2018; Sevestre et al. 2021).

**Figure 2:**
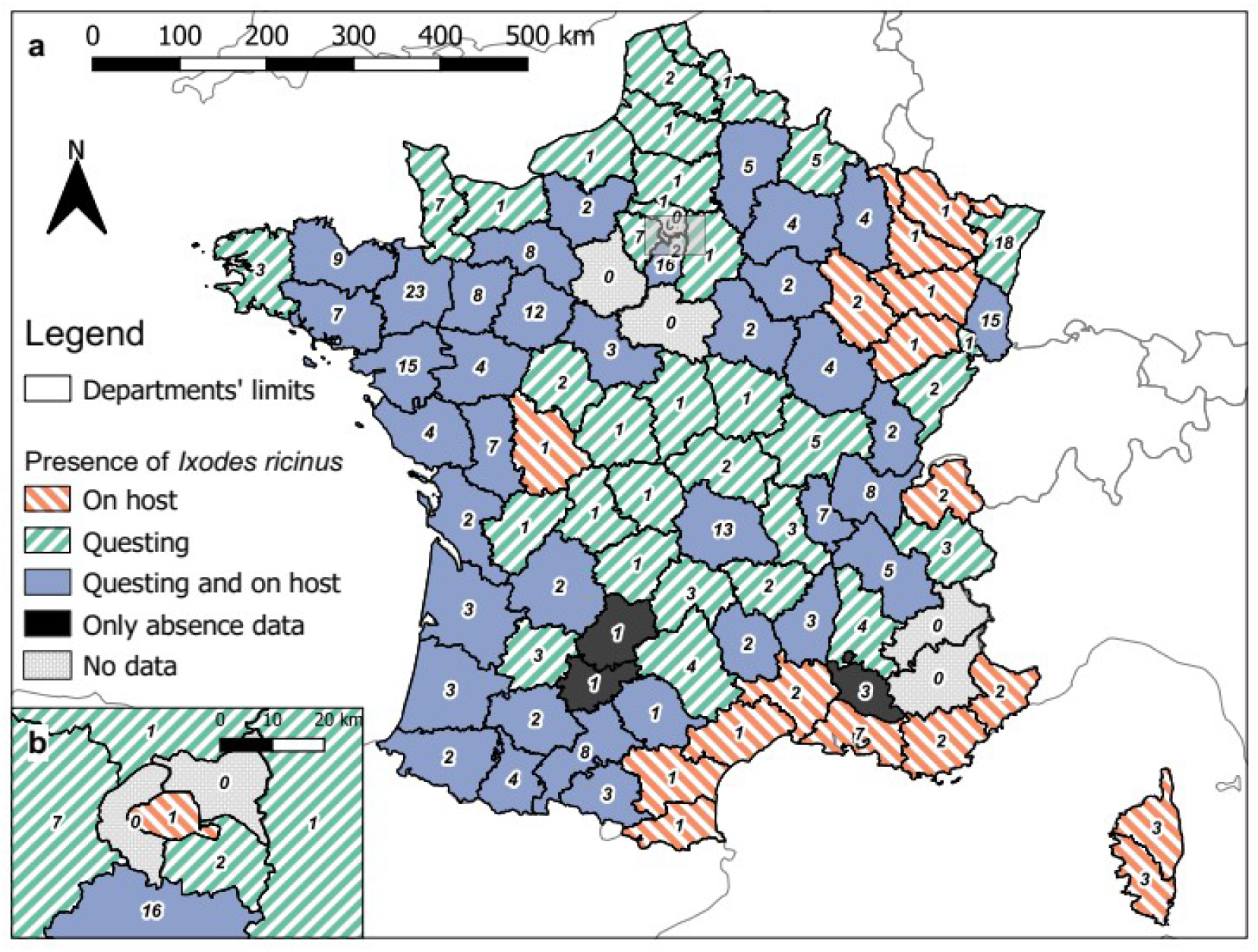
Distribution map of *Ixodes ricinus* in European France by departments according to the final 187 references, and whether ticks were questing or attached on host. Refer to **SI 7** for the name of the departments.

Climate conditions above a certain altitude also limit *I*. *ricinus*. For instance, Gilot et al. (1989) in the Rhône Valley recorded the species from 200 to 1,150 m a.s.l., but the species was more recently collected at the top of the Pic de Bazès, in the eastern part of the French Pyrenees, at 1,800 m a.s.l. (Akl et al. 2019). These data support other observations in Europe that have shown that its presence is increasingly detected at higher altitudes, beyond the previously established 1,500 m (Gern, Morán Cadenas & Burri 2008; Danielova et al. 2006; Gilbert 2010; Martello et al. 2014; Garcia-Vozmediano et al. 2020).

The species was also considered to be scarce on the Atlantic coast because of coastal environmental conditions, like wind and spray which can be highly desiccative (Degeilh et al. 1994). This tick is nonetheless present on Belle-Île-en-Mer island (Bonnet et al. 2007), indicating that the species can be present on islands, but at low abundance. Indeed, ten years after the first study, Michelet et al. (2016) did not collect any *I*. *ricinus* ticks on this same island.

At the landscape level, the presence of the species is associated with wooded areas, either deciduous or deciduous/coniferous mixed forests, small woods, copses and hedgerows in agricultural matrix, and can be found in adjacent grassland and wet meadows. The species is rare in other open habitats, like crops and grasslands, but can be found in public parks depending on their level of forestation (Gilot, Pautou & Moncada 1975; Doche et al. 1993; Degeilh et al. 1994; Gilot et al. 1994; Doby & Degeilh 1996; Pichot et al. 1997; Mémeteau et al. 1998; Boyard et al. 2007; Boyard et al. 2011; Agoulon et al. 2012; Perez et al. 2016; Vourc’h et al. 2016; Ehrmann et al. 2017; Goldstein et al. 2018; Mathews-Martin et al. 2020). Landscapes with a moderately fragmented forest, which provides forest-grassland or forest-agriculture ecotones and harbouring high host densities can favour high nymph abundance (Perez et al. 2016; Wongnak et al. 2022).

#### 3.2.2 Temporal variation in the density of questing *Ixodes ricinus*

Of the 98 articles that sampled questing ticks, 42 analysed temporal variation in tick density, either per sampling area (e.g., nymphs/10m²; n = 31) or per sampling time (e.g., nymphs per hour; n = 11). Both the number of collections and sampling intervals varied strongly among these studies (from a single sample to monthly sampling over several years and from one sample per year to several samples on the same day). Studies have mostly focused on nymphs, with few reports on annual or seasonal activity patterns of larvae. As *I*. *ricinus* is susceptible to desiccation, weather conditions, and particularly temperature and humidity, were most frequently examined. Here, we report variables that have been linked to variation in *I*. *ricinus* activity, without assessing potential bias.

Temperature is an important factor affecting tick activity. In France, *I. ricinus* activity seems optimal at about 16°C with a very sharp drop below 5°C (Aubert 1975; Tomkins et al. 2014; Cat et al. 2017). Nonetheless, it should be noticed that despite negative temperatures the days before collection (as low as -7°C), attached feeding-ticks are reported on large mammals, possibly activated by the animal’s warmth while resting (Doby et al. 1994).

Conversely, temperatures above 30°C inhibit tick activity, even if residual activity can persist (Cat et al 2017). More generally, it is the cumulative temperature over several days that affects the activity of *I. ricinus* (Perret et al. 2000; Tagliapietra et al. 2011). For instance, in north-eastern France, adult tick feeding activity – estimated by individuals found feeding on foxes – was triggered by a 10-day mean maximum temperature above 9°C prior to the fox hunt, while a mean minimum 10-day temperature below 1°C inhibited it (Aubert 1975). It is nonetheless important to note that threshold temperatures for tick activity could vary among climates if tick populations undergo local adaptation, with ticks remaining active at lower temperatures in colder climatic regions for example (Tomkins et al. 2014; Gilbert et al. 2014).

Combined with temperature, humidity is also an important limiting factor for the survival and activity of *I. ricinus* ticks. It is generally considered that ticks need a relative humidity greater than 70-80% to maintain water balance (Gray et al. 2016). Therefore, ticks regularly interrupt host-seeking activity and move down to moist litter to rehydrate. Saturation deficit, which integrates both relative humidity and temperature, is considered as a better predictor of tick activity than temperature or humidity alone (Perret et al. 2000; Hauser et al. 2018). In France, it was reported that the activity of *I*. *ricinus* nymphs could be partially predicted by the mean 7-day minimal relative humidity, and marginally by precipitation in the four weeks prior to sampling, two determinants of the water available in the environment (Paul et al 2016; Cat et al. 2017).

Photoperiod can also drive questing activity of *I*. *ricinus* as it influences the beginning of developmental diapause and the end of behavioural diapause (Gray et al. 2016). Briefly, decreasing daylight induces a developmental diapause in eggs and fed larvae and nymphs, delaying development for at least three months, and results in a behavioural diapause of tick activity (Perret et al. 2003). Conversely, increasing daylight triggers tick activity for nymphs and adults. Darkness is favourable for the horizontal displacement of ticks, likely because it is generally associated with a reduction in desiccation risk (Perret et al. 2003).

As daylight, temperature and humidity change daily, tick activity is subject to daily variation. Recent results from eastern France (near the city of Lyon, Rhône) suggest that *I. ricinus* nymphal activity in spring decreases during the day (between 7 am and 4 pm) with increasing temperatures and decreasing humidity (Kraemer 2018). Another study, carried out in south-western France (Haute-Garonne), suggested that ticks were more active during the afternoon (3-6 pm) in winter, at the end of the day in spring and summer, and remained active at night in summer, as long as temperature and humidity are favourable (Coiffait 2019). These results are concordant with other studies in Europe showing increasing activity after sunrise in cold months, decreasing activity and shift in activity toward the night in hot months (Mejlon 1997; Zöldi et al. 2013; Edwards & Campbell 2021) . Nocturnal activity in summer could explain an increase in tick infestation of nocturnal and crepuscular host species like red foxes (Aubert 1975).

It was hypothesized that daily tick activity is adapted to host activity to increase encounter rate. For instance, Matuschka et al. (1990) observed in laboratory conditions that larvae feeding on different host species detached at different periods of the day, but whether this is an effect of host activity (e.g., grooming, exploratory movements) or a behavioural adaptation remains unknown. Recently, in south-western France, Coiffait (2019) compared the daily activity of adult ticks with the daily activity of roe deer, *Capreolus capreolus*, which have a crepuscular activity, but found no relationship. However, these preliminary results need to be confirmed insofar as roe deer behaviour itself can be modified in anthropic landscapes (Bonnot et al. 2013). More data are thus needed on the influence of local host community on tick activity of all stages.

Given these dependencies, tick activity may vary strongly over a year and under different climates, resulting in seasonal and geographical variation in exposure risk for humans and their domestic animals. In France, the general phenology of *I*. *ricinus* seems to be the same as in other parts of Europe under the same climates (Kurtenbach et al. 2006). In temperate oceanic and temperate semi-continental climates, the activity peak of larvae is generally reached in late spring or summer, depending on meteorological conditions (L’Hostis et al. 1995; Agoulon et al. 2019; Bournez et al. 2020). However, data on larval activity are scarce and differences in phenology should be confirmed. Moreover, vegetation height can bias larval sampling, which typically quest low in the vegetation (Dobson et al. 2011). Larvae are scarce or absent in winter, suggesting that even if a female tick feeds and lays in late autumn or early winter, hatching occurs only in spring (L’Hostis et al. 1995). Under climates with mild temperatures throughout the year, as in the temperate oceanic climate of western France, nymphs can be found throughout the year since the required minimum favourable abiotic conditions are met (Agoulon et al. 2019). Under these climates, the peak in nymphal activity occurs between March and June (Cat et al. 2017; Degeilh 1996; Agoulon et al. 2019). This peak can be delayed in years with mild springs and shortened by high local host densities that rapidly remove ticks from the questing population (Randolph and Steele 1985; Vassallo, Paul & Pérez-Eid 2000). Under semi-continental climates, the peak of nymphal activity is observed in May-June (Pérez-Eid 1989; Ferquel et al. 2006; Beytout et al. 2007; Goldstein et al. 2018; Bournez et al. 2020a) with no (or very few) questing nymphs found during the cold season. A weaker peak of activity was sometimes observed in September-October (L’Hostis et al. 1995; Degeilh et al.1996; Cat et al. 2017; Lejal et al. 2019b); this second peak could result from the emergence of larvae fed in spring rather than to the revived activity of nymphs which survived the summer (Randolph and Steele 1985; Bregnard et al. 2021). Under mountainous climates, a unique peak of activity is observed in summer (Jouda, Perret & Gern 2004). As adults are less susceptible to desiccation than immature stages (Perret, Rais & Gern 2004), their activity tends to be more widespread through the year (Pérez-Eid 1989), but still follows the same general pattern as nymphs (Aubert 1975; Degeilh et al. 1996; L’Hostis et al.1996a; L’Hostis et al.1996b; Marchant et al. 2017; Goldstein et al. 2018; Agoulon et al. 2019). Under Mediterranean climates, like in Corsica, the peak in adult activity is in autumn, while data on larval and nymphal activity are lacking (Grech-Angelini et al. 2016).

To study the trends in population dynamics of *I*. *ricinus*, long-term data are needed to dampen year-to-year variation. Paul et al. (2016) sampled questing ticks every spring between 2008 and 2014 in a suburban forest in the Paris region (Essonne). This study reported that main climatic variables explaining inter-annual variation in the density of questing ticks were the number of days below zero degrees within a year and minimal temperatures 8-9 months prior to tick collection. Thus, low winter temperatures, but also large temperature variation in winter and extreme climatic events could have a negative effect on tick survival, explaining part of the annual variation in tick density (Herrmann & Gern 2013; Paul et al. 2016; Hauser et al. 2018). However, in Paul et al. (2016), inter-annual fluctuations in questing nymph density were considered to be more closely linked to fluctuations in vertebrate host densities and vegetation cover than to meteorological data per se. Similar inferences were made in longitudinal studies performed in Switzerland and Germany (Brugger et al. 2018; Hauser et al. 2018; Bregnard, Rais & Voordouw 2020). Perez et al. (2016) also reported that nymphal density was associated with both the presence of larvae and the abundance of small mammals the previous year. Larval density was itself associated with the proportion of woodlands in the surrounding landscape, a privileged habitat for roe deer, which are the main hosts of adult females (Perez et al. 2016; Morellet et al. 2011).

Despite some studies documenting a northward and altitudinal expansion of *I*. *ricinus* in Europe, probably linked to climate changes, few sampling campaigns have been conducted over a sufficiently long time period to test an effect of climate change on tick abundance at more local scales (Lindgren et al.; 2000; Léger et al. 2013; Medlock et al. 2013). To our knowledge, the longest *I*. *ricinus* yearly population survey, conducted in the eastern part of the species range (Tula region, western Russia) between 1977 and 2011, showed a general increase in adult abundance since 1990 in relation to an increase in degree days (cumulated daily temperature) during the August-October period. This increase was also associated with changes in forest coverage and fragmentation observed in the region following socio-economic changes (Korotkov, Kozlova & Kozlovskaya 2015). Thus, while weather conditions have an influence on annual tick population densities, hosts and landscape changes seem to have an influence as well. These factors are nonetheless difficult to disentangle since they generally interact.

#### 3.2.3 Host use

Out of 59 articles reporting tick sampled on hosts, only 33 (55.9%) presented sufficient details to be informative on host-associated infestation rates. The potential vertebrate host species of *I*. *ricinus* studied in France and the degree of parasitism by each life stage is presented in (**SI 8**). Overall, 55 species were examined for the presence of *I*. *ricinus* ticks. These species included 29 mammalian species (4 Carnivora, 8 Cetartiodactyla, 4 Eulipotyphla, 1 Lagomorpha, 1 Perissodactyla and 11 Rodentia) reported in 34 articles, and 26 avian species (24 Passiformes and 2 Piciformes) reported in 4 articles. No data on Squamata (i.e., lizards and snakes) were found. More detailed data on parasitism (i.e., infesting life stages and/or prevalence/intensity of infestation) were available in 26 articles for mammalian hosts, but in only two studies for avian hosts. In the first avian study, 21 different avian species were included (Marsot et al. 2012), while in the second one, only the Eurasian blackbird, *Turdus merula,* was considered (Grégoire et al. 2002).

Data from different studies preclude direct comparisons of host infestation levels because biogeographic zones and sampling season usually differ from one study to another. For instance, wild ungulates are typically studied in winter, during the hunting season, when ticks are less active (Doby et al. 1994), whereas birds are captured in spring, when ticks are highly active (Marsot et al. 2012), and small mammals are trapped from spring to autumn (L’Hostis et al. 1996b; Perez et al. 2016). In addition, it is not possible to compare levels of host infestation when comparing areas where *I*. *ricinus* is very frequent, like in the northern part of France (Pérez-Eid 1990; Doby et al. 1994; L’Hostis et al. 1996b) to areas where this species is very rare, like in Corsica (Grech-Angelini et al. 2016). Finally, the modalities (e.g., time of inspection, body part inspected) of tick collection on hosts also varies from one study to another. For instance, wild boars, *Sus scrofa,* were inspected only five minutes in the field (Doby et al. 1994), while red foxes *Vulpes vulpes* were meticulously inspected in laboratory (Aubert et al. 1975). These limits highlight the need to standardize studies, controlling for spatial and temporal factors, and employing comparable on-host sampling strategies. For instance, two countrywide standardised studies based on tick collections on red squirrels, *Sciurus vulgaris*, confirmed that *I. ricinus* is rare under Mediterranean climates compared to Atlantic, semi-continental and mountainous climates (Romeo et al. 2013; Pisanu et al. 2014).

The proportion of each *I*. *ricinus* life stage present on a given host species at a given time, can nonetheless provide some indication on their role as blood source and can enable us to formulate testable hypotheses for future work. On small rodents (except for Eurasian red squirrels and Siberian chipmunks), larvae were predominant whereas nymphs were rare (Anderson et al. 1986; Doby et al. 1992; L’Hostis et al. 1996b; Richter et al. 2004; Vourc’h et al. 2007; Boyard, Vourc’h & Barnoin 2008; Marsot et al. 2013; Romeo et al. 2013; Pisanu et al. 2014; Le Coeur et al. 2015; Perez et al. 2017; Pérez-eid 1990; Bournez et al. 2020a). Among rodents, adult ticks were observed only on black rats, *Rattus rattus*, from Corsica (Cicculli et al. 2019). We can note that this observation questions the role of black rats in transporting ticks in peri-urban and urban areas.

Larvae and nymphs were frequent on passerine birds, with sometimes more nymphs than larvae (Grégoire et al. 2002; Marsot et al. 2012). In eastern France, Grégoire et al. (2002) found that blackbirds were significantly more infested in rural habitats (74%) compared to urban habitats (2%). This suggests a scarcity of ticks in urban areas compared to rural woodlands, but also a possible role of these hosts in disseminating ticks in urban and peri-urban green spaces. A similar pattern was observed when comparing tick densities in a reference forest plot and peri-urban and urban parks in the City of Lyon (eastern France); ticks were more abundant in the forest plot than in the peri-urban and in urban parks (Mathews-Martin et al. 2020).

Females, nymphs and larvae were all observed on wild ungulates, sometimes with more adults than nymphs (e.g., roe deer, red deer, *Cervus elaphus*, and Pyrenean chamois, *Rupicapra pyrenaica*) (Davoust et al. 2012; Doby et al. 1994; Gilot et al. 1994a). Domestic ungulates can also host numerous female ticks with up to 73 per individual on cattle (L’Hostis, Bureaud & Gorenflot 1996). However, there is no strong evidence that domestic ungulates contribute to an increase of *I*. *ricinus* populations. For instance, Ruiz-Fons et al. (2012) found a positive relationship of horse abundance with questing larvae, but not nymph, abundance, and Sprong et al. (2019) found a weak negative relationship of cattle abundance with questing adult abundance, while Steigedal et al. (2013) found a negative relationship of sheep presence with nymph and adult abundance. However, in neither case the authors quantified ticks on cattle, horse or sheep, although they may had removed a significant part of the questing ticks (Randolph & Steele 1985). Furthermore, Ruiz-Fons et al. (2012) and Sprong et al. (2019) found a positive relationship of cattle abundance with *Anaplasma phagocytophilum* prevalence, a pathogen for which cattle are reservoir hosts, suggesting that they had fed a significant part of the tick population.

Few studies were performed and report *I*. *ricinus* ticks from wild carnivores in France and all were on foxes (Aubert et al. 1975; Doby et al. 1991). Given their low density compared to other host species, their contribution in feeding ticks is likely low (Hofmeester et al. 2016). The presence of wild Carnivores might even have a negative effect on tick populations due to the predation pressure and behavioural modifications they exert on larvae hosts such as small mammals (Hofmeester et al. 2017). More studies are needed to evaluate the importance of wild Carnivores in the population dynamics and maintenance of *I*. *ricinus* populations.

Studies on domestic Carnivores have focused mainly on infested dogs and cats and frequently fail to report sampling effort, precluding an estimate of infestation level (Panas et al. 1976; Pichot et al. 1997; Gilot, Pichot & Doche 1989; Ulmer et al. 1999; Geurden et al. 2018; Grech-Angelini et al. 2016). However, cats and dogs can be highly infested, with up to 26 and 64 *I*. *ricinus* ticks, respectively (Geurden et al. 2018). Infestation levels of domestic Carnivores are also influenced by environmental conditions. For instance, Martinod, Brossard & Moreau (1985) showed that dog infestation by *I*. *ricinus* varied greatly between two areas in a 40 km² zone in eastern France (southern Jura mountains) in relation to land use (5.3 and 0.032 *I*. *ricinus* adult ticks per dogs on 249 dogs from areas with pastures and residential neighbourhoods and 190 dogs from mountainous and plain areas). Like domestic ungulates, the contribution of domestic Carnivores to the population dynamics of *I*. *ricinus* is unclear. Indeed, domestic vertebrates may act as ecological traps for ticks because of their breeding conditions (housing, antiparasitic treatments).

Despite limited data, observations on host use in France are in general agreement with what is known for the host range of *I. ricinus* across its range (Hofmeester et al. 2016). However, more data is required to better understand the relative contribution of these different host types and to evaluate the contribution of understudied groups (Eulipotyphla, Lagomorpha, Perissodactyla and Squamata) in France and elsewhere.

### 3.3 Tick-borne pathogens in *Ixodes ricinus* ticks of France

Different tick-borne pathogens, that have been examined in *I*. *ricinus* ticks in France, are briefly presented below (see McCoy & Boulanger for more details). We summarised existing data from across studies and present these data in detail in the Supplementary Information (**SI 9-17**). Not enough data exist to establish prevalence maps for most pathogens (with the exception of *Borrelia burgdorferi* s.l.), but presence/absence information is illustrated (**Figures 3-10**).

#### 3.3.1 The genus *Anaplasma*

Bacteria of the genus *Anaplasma* are obligate Gram-negative alphaproteobacteria. They are intracellular pathogens transmitted by ticks and responsible for anaplasmoses in vertebrates. Among this genus, *A*. *phagocytophilum* is the most prevalent species in northern Europe and infects monocytes and granulocytes of ruminants, small mammals, horses, canids, birds and humans (Keesing et al. 2012). Several genetic variants circulate in Europe with some host and vector specificities. Jahfari et al. (2014) identified four genetic variants called “ecotypes” associated with different host groups in Europe. According to these authors, ecotype I is associated with ungulates, dogs, hedgehogs and humans; ecotype II is associated mainly with roe deer; ecotype III is associated with rodents; and ecotype IV is associated with birds, results confirmed by other studies (Jahfari et al. 2014; Stigum et al. 2019; Grassi et al. 2021). These ecotypes also differ by their vectors. Ecotypes I and II are mainly transmitted by *I*. *ricinus;* ecotype III seems mainly transmitted by *I*. *trianguliceps*; vectors of ecotype IV are still uncertain, but are suspected to be among the hard ticks that specifically exploit birds; and an ecotype IV-like was associated with *I*. v*entalloi,* a tick species mostly feeding on European rabbits, *Oryctolagus cuniculus*, and present in France (Gilot, Rogers & Lachet 1985; Santos et al. 2018; Jaarsma et al. 2019). Transmission of some genetic variants by ticks of the genus *Dermacentor* is suspected, but has not been demonstrated (Baldridge et al. 2009). Other tick species might be vector of other specific genetic variants. Anaplasmoses can be of significant medical and economic importance when infecting humans and livestock. It may cause severe disease marked by anemia and leukopenia, fever, headache, and myalgia (Rymaszewska & Grenda 2008; Stuen et al. 2013; Dugat et al. 2015). Few human cases are diagnosed in France, and occur mainly in the eastern part of the country (Koebel et al. 2012).

DNA of *A*. *phagocytophilum* bacterium has been detected by PCR methods in questing *I. ricinus* from almost all regions of France and prevalence in ticks shows some geographic disparities (**Figure 3; SI 9** and references associated therein), which may be explained, in part, by wild and domestic ungulate densities (Chastagner et al. 2017). Prevalence varied from 0.4% to 5.3% in questing nymphs and from 0.5% to 7% in questing adults.

**Figure 3:**
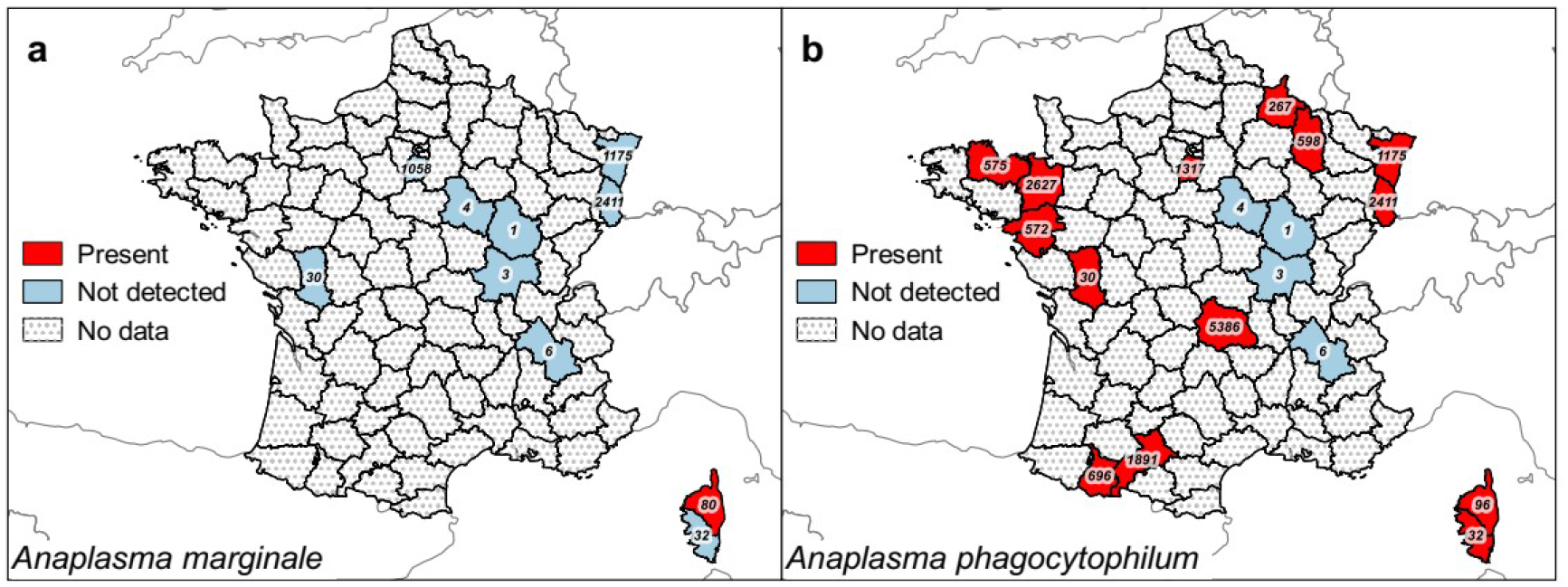
Presence maps of *Anaplasma marginale* and *A*. *phagocytophilum* in questing and attached nymph and adult *Ixodes ricinus* ticks in European France by department. Detected presence by PCR methods and respective sampling effort per department in number of tested ticks (nymphs and adults) for *A*. *marginale* (**a**) and *A*. *phagocytophilum* (**b**).

Only five studies have attempted to detect DNA of other *Anaplasma* species in questing *I*. *ricinus* ticks in France (Bonnet et al. 2013; Michelet et al. 2014; Lejal et al. 2019a; Lejal et al. 2019b; Grech-Angelini et al. 2020a). These other *Anaplasma* species infecting animals are *A*. *centrale*, *A*. *marginale* and *A*. *ovis*, infecting erythrocytes of ruminants, *A*. (*Ehrlichia*) *bovis*, infecting monocytes and granulocytes of ruminants, and *A*. (*E*.) *platys* infecting mainly platelets of dogs (Rymaszewska & Grenda 2008). However, *I*. *ricinus* is not considered to be the vector of these pathogens. Among these other species, *A*. *marginale* was detected in only one study in *I. ricinus* ticks feeding on cattle and red deer in Corsica with a prevalence of 1.7% (Grech-Angelini et al. 2020a). The presence of these other *Anaplasma* species in *I*. *ricinus* ticks in France is thus currently anecdotal (*A*. *marginale*) (see **SI 9** for further details).

#### 3.3.2 The genus *Bartonella*

The *Bartonella* genus is composed of facultative intracellular bacteria transmitted by arthropods such as fleas, lice, haematophagous Diptera, and Acari including ticks, and occurs worldwide (Billeter et al. 2008). Known reservoirs are cats, canids and rodents (Chomel, Boulouis & Breitschwerdt 2004). The main pathogenic species for humans are *B*. *bacilliformis*, *B*. *henselae* and *B*. *quintana* (Angelakis & Raoult 2014), but only *B*. *henselae* is of known concern in France. It is responsible for cat-scratch disease which is usually characterized by mild symptoms (aches, malaise), but can sometimes lead to more severe symptoms like endocarditis or meningoencephalitis. Although *B*. *henselae* is mainly transmitted among the cat reservoirs via fleas (*Ctenocephalides felis felis*) (Chomel et al. 1996), *I*. *ricinus* is an experimentally confirmed vector and other tick species and haematophagous Diptera are suspected vectors (Billeter et al. 2008; Cotté et al. 2008; Grech-Angelini et al. 2020a; Wechtaisong 2020). The species *B*. *birtlessi* that infect rodents is also transmitted by *I*. *ricinus*, but is likely not pathogenic for humans (Reis et al. 2011b).

In France, *Bartonella* spp. have been found from different departments, but with limited sampling effort (**Figure 4a**; **SI 10**). The bacterium *B*. *henselae* was the most frequently detected *Bartonella* species in *I*. *ricinus*, varying from 0% to 38% (**Figure 4b**; **SI 10**). DNA of this bacterium was found in questing nymphs and adults sampled in north-eastern France (Ardennes and Bas-Rhin) (Dietrich et al. 2010; Michelet et al. 2014; Moutailler et al. 2016b), and in feeding adult ticks in Corsica (Grech-Angelini 2020a). In the Paris region (Essonne), DNA of *B*. *birtlesii* was detected in questing *I*. r*icinus* nymphs (Reis et al. 2011b). Other *Bartonella* spp. have been detected in other parts of France, but without an identification to species (Halos et al. 2005; Cotté et al. 2010; Davoust et al. 2012; Bonnet et al. 2013; Paul et al. 2016; Bonnet et al. 2017; Nebbak et al. 2019).

**Figure 4:**
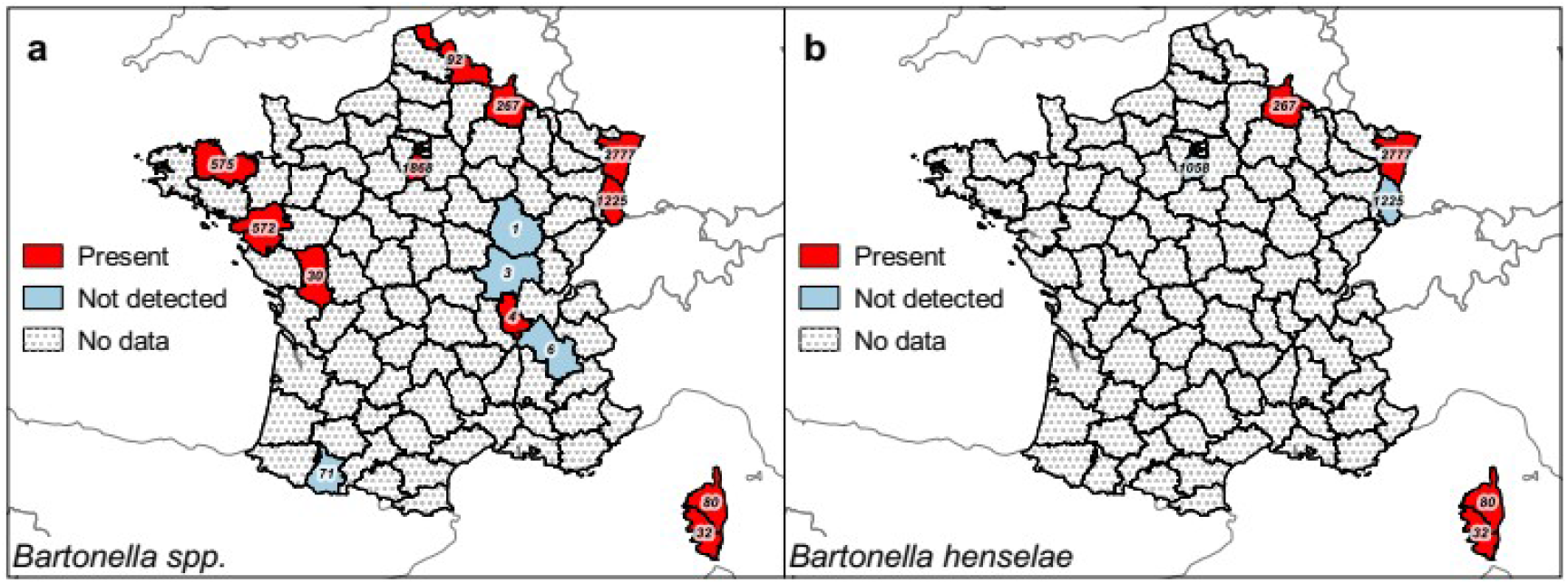
Presence maps of *Bartonella* spp. and *B*. *henselae* alone in questing and attached nymph and adult *Ixodes ricinus* ticks in European France by department. Detected presence by PCR methods and respective sampling effort per department in number of tested ticks (nymphs and adults) for *Bartonella* spp. (**a**) and *B*. *henselae* alone (**b**).

#### 3.3.3 *Borrelia burgdorferi* sensu lato (s.l.) complex

Some bacterial species of the *B. burgdorferi* s.l. complex are responsible for Lyme borreliosis, the most important vector-borne disease for humans in the northern hemisphere (Borchers et al. 2015) and that most frequent in France. In 2020, the French national surveillance programme estimated a mean incidence of 90 cases / 100,000 inhabitants, varying between 11 and 667 cases/ 100,000 inhabitants depending on the region (https://www.sentiweb.fr). In Europe, several genospecies (genetically-discriminated species) of the complex circulate in different reservoir hosts and can be associated with different pathogenicity (Steinbrink et al. 2022). Small mammal-associated genospecies include *B*. *afzelii*, *B*. *garinii* subsp. *bavariensis*, *B*. *bissettii*, and *B*. *spielmanii*, whereas *B*. *garinii garinii*, *B*. *turdi*, and *B*. *valaisiana* are considered to be bird-associated. *B*. *burgdorferi* sensu stricto (s.s.) is harboured by both small mammals and birds, whereas *B*. *lusitaniae* is mostly associated with lizards. There is no known reservoir host for *B*. *finlandensis* (Wolcott et al. 2021). The species most frequently involved in human cases of Lyme borreliosis in Europe are *B*. *afzelii* and *B*. *garinii* (Stanek & Reiter 2011), causing cutaneous and neurological forms of the disease respectively (Strle et al. 2006). Both *B*. *burgdorferi* s.s. and *B. garinii* subsp. *bavariensis* (formerly *B*. *garinii* OspA serotype 4, but not differentiated from *B*. *garinii garinii* in all studies; Hördt et al. 2020) are less frequent in Europe, but are also disease-causing; *B. burgdorferi* s.s. is associated with neurological and arthritic forms of Lyme borreliosis, whereas *B*. *garinii* subsp. *bavariensis* only manifests neurological forms (Wilske et al. 1996; Margos et al. 2013). Other circulating genospecies like *B*. *spielmanii*, *B*. *bissettii*, *B*. *lusitaniae* and *B*. *valaisiana* rarely cause human borreliosis (Steinbrink et al. 2022). The genospecies *B*. *finlandensis* and *B*. *turdi* are only rarely detected in *I*. *ricinus* ticks and their pathogenicity for humans is unknown (Steinbrink et al. 2022). The role of *I*. *ricinus* as a vector has been experimentally demonstrated for *B*. *afzelii*, *B*. *garinii* and *B*. *burgdorferi* s.s. (Eisen 2020).

DNA from different *B*. *burgdorferi* s.l. genospecies, except *B*. *garinii* subsp. *bavariensis*, *B*. *bissettii* and *B*. *finlandensis*, has been detected in questing *I*. *ricinus* ticks collected in France (**SI 11**). Although studies on *B*. *burgdorferi* s.l. have focused on a few French departments, the highest prevalence to date is found in the north-eastern, eastern and central parts of France, where prevalence ranges from 2.0 to 26.2% in nymphs (**Figure 5a** and **5b**; **SI 11** and references therein). The prevalence of the different genospecies in questing nymphs can be compared between regions from which a sufficient number of ticks were tested. In the north-eastern quarter of France, *B*. *afzelii* was the most frequently detected genospecies (35-50% of positive ticks) and the only one in Corsica (**Figure 5c**). The prevalence of *B*. *burgdorferi* s.s. was generally found to be low across France. However, in one study in northern France, it was the predominant species (32% of positive ticks); the authors suspect that the high prevalence of this genospecies might result from the introduction of the Siberian chipmunk in the sampled forest, which is a competent reservoir with a higher tick burden than local rodent species (**Figure 5d**) (Pisanu et al. 2010; Marsot et al. 2011; Jacquot et al. 2014; Bonnet et al. 2015; Jacquot et al. 2016). This genospecies was also the most predominant in southern France (32%) where the overall *B*. *burgdorferi* s.l. prevalence was low. In north-western France, *B*. *garinii* was the genospecies most frequently detected (26%), but at low prevalence compared to those observed in the north-eastern quarter of France, again because of an overall low *B*. *burgdorferi* s.l. prevalence (**Figure 5e**). The prevalence of *B*. *valaisiana* varied between 0 and 21% of positive ticks, but did not follow any clear pattern in prevalence (**Figure 5f**).

The higher *B*. *burgdorferi* s.l. prevalence observed in north-eastern, eastern and central regions can be explained by their high proportion of forested landscapes that support higher wild vertebrate densities, in particular wild ungulates (that are good amplifiers of ticks), and wild rodents and birds (reservoirs of the bacteria), compared to northern and western regions with more open landscapes and less forested areas. Given the strong host specificity of genospecies, regional disparities in prevalence in questing nymphs suggests that tick host use differs between regions with different climates, landscapes and host communities. For instance, the proportion of *Borrelia* species harboured by avian species (e.g., *B*. *garinii*) seems higher in western regions and those harboured by small mammal species (e.g., *B*. *afzelii*) seem higher in north-eastern regions. Nonetheless, when considering genospecies prevalence, these results are mostly in accordance with the E-W geographical gradients in *B*. *burgdorferi* s.l. prevalence observed across Europe (Strnad et al. 2017).

**Figure 5:**
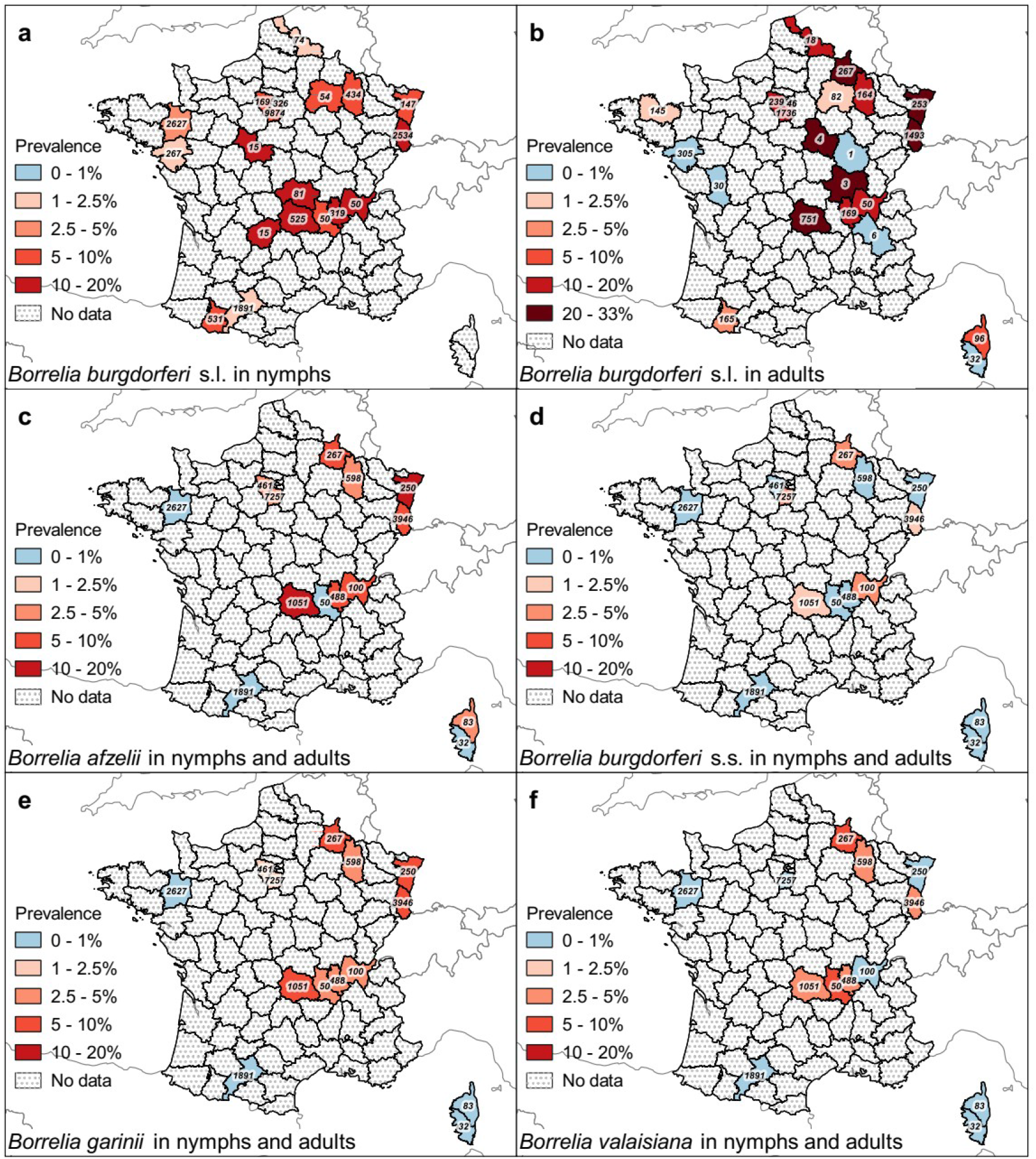
Prevalence maps of *Borrelia burgdorferi* sensu lato and *B*. *afzelii*, *B*. *burgdorferi* sensu stricto, *B*. *garinii*, and *B*. *valaisiana* in questing and attached nymph and adult *Ixodes ricinus* ticks in European France by department. Prevalence, corresponding to the minimum detected prevalence by PCR methods, and sampling effort by department, in number of tested ticks, for *B*. *burghdorferi* s.l. in nymphs (**a)** and in adults (**b**), and in nymph and adult ticks together for the genospecies *B*. *afzelii* (**c**), *B*. *burgdorferi* s.s. (**d**), *B*. *garinii* (**e**) and *B*. *valaisiana* (**f**).

The *B*. *burgdorferi* s.l. prevalence was also found to vary from site to site and from year to year within regions (Boulanger et al. 2018). For instance, in a four-year follow-up on four sites in Alsace region, Boulanger et al. (2018) reported prevalence varying from 0.7% to 13.6% between sites in the same year, and from 7.7% to 26.7% between years for the same site. Studies at landscape and local scales suggest that pathogen prevalence is positively associated with wooded habitat ecotones (Halos et al. 2010; Perez et al. 2020), but negatively to habitat diversity (Ehrman et al. 2018). This may be explained by differences in the proportion of each host group used by larvae, which might also vary temporally because of high fluctuations in small mammal population sizes (Perez et al. 2017).

#### 3.3.4 Borrelia miyamotoi bacterium

The bacterium *B*. *miyamotoi* is currently the only known species among *Borrelia* relapsing fever (RF) bacteria to be transmitted by a hard tick in Europe; soft ticks and lice are the vectors of the other species of this group (Platonov et al. 2011; Wagemakers et al. 2015). Suspected reservoir hosts are rodents, but these bacteria can also infect some bird species (e.g., wild turkeys, *Meleagris gallopavo,* common blackbirds) and large mammals (roe deer, wild boars) (Wagemakers et al. 2015). In humans, *B*. *miyamotoi* is responsible for acute febrile illness and meningoencephalitis (Wagemakers et al. 2015) and the number of human cases reported is continuously increasing in Europe (Henningsson et al. 2019; Tobudic et al. 2020). It has been detected in *Ixodes* ticks, including *I*. *ricinus*, from different regions of the Northern Hemisphere, and, in contrast with other *Borrelia* spp., trans-ovarial transmission in ticks is thought to occur (Wagemakers et al. 2015; van Duijvendijk et al. 2016).

DNA of *B*. *miyamotoi* was formally detected by PCR methods for the first time in north-eastern France (Ardennes), both in samples from bank voles (four positive rodents out of 72 tested – 5.5%) and from questing adult female *I*. *ricinus* ticks (eight positive out of 267 tested – 3.0%) (Cosson et al. 2014; Moutailler et al. 2016b). The presence of this bacterium was subsequently detected by PCR methods in the Paris region, in north-eastern France (Alsace, Champagne-Ardenne), and in Corsica island, with observed prevalence varying between 0.75 and 4.2% in questing nymphs and between 0 and 21.7% in adults (**SI 12**) (Vayssier-Taussat et al. 2013; Paul et al. 2016; Boulanger et al. 2018; Lejal et al. 2019a; Lejal et al. 2019b; Nebbak et al. 2019; Boyer et al. 2020). Other RF *Borrelia* sp. have been detected in north-eastern France (Alsace region) that could be *B*. *miyamotoi*, but the species was not identified (Richter, Schlee & Matuschka 2003).

#### 3.3.5 Coxiella burnetii bacterium

The bacterium *Coxiella burnetii* is a worldwide pathogen responsible for Q fever (Baca & Paretsky 1983). Potential reservoir host species include ticks, mammals (more than a hundred species recorded) and birds (at least twenty species recorded) (González-Barrio & Ruiz-Fons 2018; Ioannou et al. 2009). However, domestic ruminants are the most frequent reservoir host leading to human transmission (Eldin et al. 2017). The disease causes fever, myalgia, nausea, diarrhea, endocarditis and hepatitis or meningoencephalitis in humans. In ruminants, it is responsible for reproductive disorders and abortions (Agerholm 2013; González-Barrio & Ruiz-Fons 2018; Eldin et al. 2017). DNA of this bacterium has been found in more than 40 hard tick species and in 14 soft tick species (Eldin et al. 2017), but vector competence has only been experimentally confirmed for few species (*D*. *andersoni*, *Haemaphysalis humerosa*, H. *aegyptium*, *H. asiaticum*, *I*. *holocyclus, Ornithodoros hermsi*, and *O*. *moubata*) (Duron et al. 2015). The competence of *I*. *ricinus* has not been tested, and tick bites are likely not the main transmission route of the bacteria (see in Duron et al. 2015). Indeed, infections can be acquired directly by aerosols from infected vertebrate feces and urine (Eldin et al. 2017) or via the milk of infected female mammals (Pexara, Solomakos & Govaris 2018; Eldin et al. 2016). The bacterium is also excreted in tick feces, which probably contribute to its dissemination in the environment and to infection via aerosol transmission (Körner et al. 2020).

**Figure 6:**
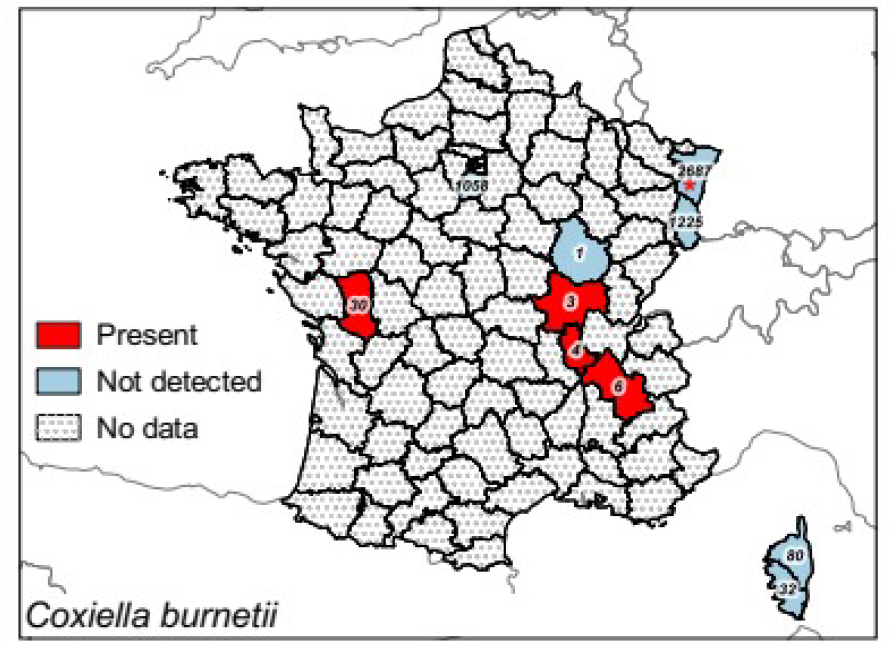
Presence map of *Coxiella burnetii* in questing and attached nymph and adult *Ixodes ricinus* ticks in European France by department. Detected presence by PCR methods and respective sampling effort per department in number of tested ticks (nymphs and adults) for *C*. *burnetii*. * DNA of an unidentified *Coxiella* sp, was detected.

In France, few studies have searched for *C*. *burnetii* in questing *I*. *ricinus* (**Figure 6**; **SI 13**). For this bacteria, sequencing is essential to avoid confusion with *Coxiella*-like symbionts (Jourdain et al. 2015). Bonnet et al. (2013) detected DNA of this bacterium in adult ticks from four out of five departments sampled (8 of 44 tested ticks, 18.2%). However, none of 998 questing nymphs, and 60 questing adult ticks from the Paris region (Essonne) were found positive (Lejal et al. 2019a; Lejal et al. 2019b), nor the 62 adults and 3,850 questing nymphs collected in Alsace (Vayssier-Taussat et al. 2013). The 115 *I*. *ricinus* ticks collected directly on diverse wild and domestic animals in Corsica were all also negative for *C*. *burnetii* DNA (Grech-Angelini et al. 2020a).

#### 3.3.6 The genus *Ehrlichia*

Species of the *Ehrlichia* genus are obligate intracellular bacteria that infect mammalian blood cells. Several *Ehrlichia* species are transmitted by ticks and can cause infections in humans and domestic animals (Rar & Golovljova 2011). The known pathogenic species for humans *E*. *chaffeensis*, *E*. *ewingii, E*. *minasensis* and *E*. *ruminantium*, and their known reservoir hosts are cervids, canids and cervids, cattle and cervids, and ruminants respectively. Only *E*. *canis*, *E*. *chaffeensis*, *E*. *minasensis* are known to be present in Europe (Rar & Golovljova 2011: Cicculli et al. 2019). Symptoms in humans are fever, headache, myalgia, and nausea, with only the more severe disease leading to death.

The role of *I*. *ricinus* as vector of these bacteria is still uncertain and requires explicit evaluation. Studies were conducted to search for *Ehrlichia* DNA in questing *I*. *ricinus* ticks sampled in France, but with mixed results. No positive ticks were found in intensive studies of questing ticks carried out in north-eastern France (Ardennes) (Moutailler et al. 2016b), in the Paris region (Essonne) (Lejal et al. 2019a; Lejal et al. 2019b), and in two sites in north-eastern France (Alsace) (Michelet et al. 2014). Similarly, no positive *I*. *ricinus* adults were collected from hosts in Corsica (Grech-Angelini et al. 2020a). DNA of the *Ehrlichia* genus was however detected in north-eastern France (Alsace region) in questing *I*. *ricinus* nymphs and adults, but the species was not identified (Ferquel et al. 2006). Another study in Alsace also found *E*. *canis* DNA in questing *I*. *ricinus* nymphs, but the prevalence could not be estimated (Vayssier-Taussat et al. 2013). Strains of *Ehrlichia* bacteria were also detected in western France (Brittany region); one was closely related to a strain from northern Italy, one was very similar to the *Ehrlichia* sp. HI-2000 strain of the *E*. *canis* group, and the last to a strain identified in Japan (*Ehrlichia* sp. Yamaguchi) and closely related to *E*. *muris*, which infects rodents (Marumoto et al. 2007). These results suggest a low prevalence of these bacteria in questing *I*. *ricinus* ticks in France.

#### 3.3.5 Neoehrlichia mikurensis

*Neoehrlichia mikurensis* is an obligatory intracellular bacterium. It was first detected in *I*. *ricinus* in the Netherlands and *I*. *ovatus* in Japan, which are – with *I*. *persulcatus –* likely the main vectors, and in rodents (brown rats, *R*. *norvegicus)* in China, the reservoir hosts (Schouls et al. 1999; Pan et al. 2003; Kawahara et al. 2004). The bacterium was also found in dog blood (Rar & Golovljova 2011). The first human cases of infection by *N. mikurensis* were described in Germany and Sweden in 2007 (von Loewenich et al. 2010; Welinder-Olsson et al. 2010). More recently, the first human cases in France were described (Boyer et al. 2021). This bacterium causes a systemic inflammatory syndrome in persons with hematological or immunological deficiencies, accompanied by fever, diarrhea and other potential complications (Silaghi et al. 2015; Portillo et al. 2018; Wass et al. 2019).

**Figure 7:**
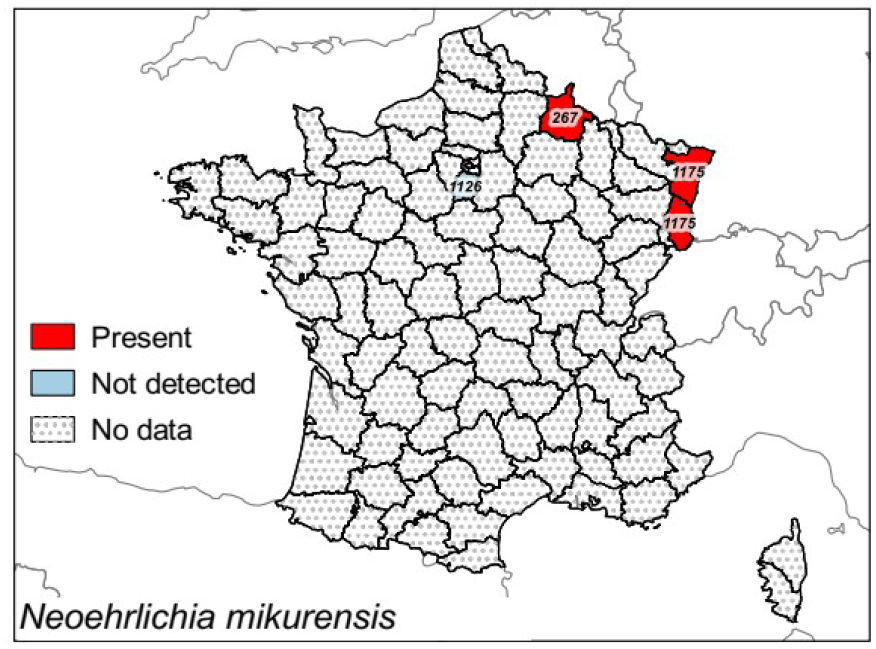
Presence map of *Neoehrlichia mikurensis* in questing and attached nymph and adult *Ixodes ricinus* ticks in European France by department. Detected presence by PCR methods and respective sampling effort per department in number of tested ticks (nymphs and adults) for *N*. *mikurensis*.

In France, DNA of the bacterium was detected in five out of 276 bank voles from north-eastern France (Ardennes) (Vayssier-Taussat et al. 2012). Questing ticks from the same areas were subsequently tested for infection and 4 out of 267 *I*. *ricinus* adult females were positive (Moutailler et al. 2016b). Other studies in north-eastern France (Alsace) have reported the presence of the bacterium: 1.7% of questing adult *I*. *ricinus* (Richter & Matuschka 2012), 1.6% (0.9%-2.5%) of questing nymphs and between 3.2%-6.4% of questing adults of the region (Vayssier-Taussat et al. 2013), and 0.2% (0.02%-0.6%) and 1.3% (0.7%-2.2%) of questing nymphs in Murbach and Wasselone respectively (Michelet et al. 2014). In contrast, no DNA of *N*. *mikurensis* was found in questing nymphs and adults from the Paris region (Essonne) (Lejal et al. 2019a; Lejal et al. 2019b), nor in Corsica from nine tick species collected from hosts, including 115 *I. ricinus* (Grech-Angelini et al. 2020a). It seems that in France *N*. *mikurensis* is well present in the north-eastern part (**Figure 7**). Given the limited number of studies to date, additional work will now be required to complete our knowledge of its overall geographical distribution in the country.

#### 3.3.6 Francisella tularensis and F. philomiragia bacteria

*Francisella tularensis* is a facultative intracellular bacterium, present in North-America, Asia and Europe, and detected in several mammal species, mainly lagomorphs and rodents, which are highly susceptible to the bacterium (Larson et al. 2020). It is responsible for tularaemia, a disease characterised in humans by endocarditis, fever, ulcer, pneumonia, septicaemia, myalgia and headache depending on subspecies, site and mode of infection (Hestvik et al. 2014; Gaci et al. 2016). It is suspected to be transmitted by biting insects such as bedbugs, deer flies and ticks and via contact with infected mammals or contaminated water or soil (Genchi et al. 2015; Telford & Goethert 2020).

The bacterium *F*. *philomiragia* causes pneumonia, fever and septicaemia in humans (Mailman & Schmidt 2005). Infections with this bacterium are however rare and involve mainly immunodeficient persons or those exposed to contaminated water in their lungs such as after a near drowning experience (Mailman & Schmidt 2005; Kreitmann et al. 2015). Not much is known about the ecology of this bacterium, but it was found in other tick species and possible ecological similarities with the related *F*. *tularensis* are suspected (Bonnet et al. 2013)

In France, studies on the presence of *F. tularensis* and *F*. *philomiragia* in questing *I*. *ricinus* ticks are rare (**Figure 8**; **SI 14**). For this bacteria too, sequencing or specific PCR are necessary to discriminate between *Francisella*-like symbionts and *F*. *tularensis* (Michelet et al. 2013). DNA of *F*. *tularensis* was detected in questing ticks sampled in Paris region (Essonne) in two studies, while that of *F*. *philomiragia* was not (Reis et al. 2011a; Paul et al. 2016). The low prevalence (0-4.3%) in questing adult *I*. *ricinus* ticks in France is similar to results found in other European countries, and suggests that *I*. *ricinus* might be only a vector of *F. tularensis* (Reye et al. 2013; Sormunen et al. 2021; Egyed et al. 2012; Kirczuk, Piotrowski & Rymaszewska 2021).

**Figure 8:**
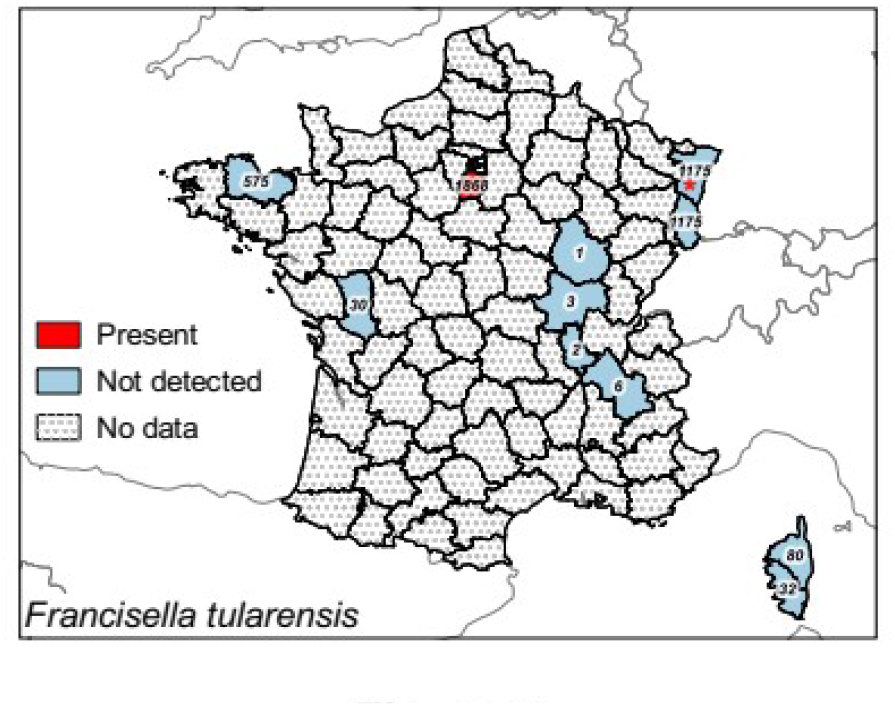
Presence map *Francisella tularensis* in questing and attached nymph and adult *Ixodes ricinus* ticks in European France by department. Detected presence by PCR methods and respective sampling effort per department in number of tested ticks (nymphs and adults) for *F*. *tularensis*. * Detection of unidentified *Fransicella* sp..

#### 3.3.7 The genus *Rickettsia*

The genus *Rickettsia* includes more than thirty intracellular bacterial species. The pathogenic species are separated into two groups: the “typhus” group (TG), with two insect-transmitted species (*R*. *typhi* transmitted by fleas and *R*. *prowazekii* transmitted by lice) and the “spotted fever” group (SFG), with most species transmitted by hard ticks (except for *R. felis* transmitted by fleas and *R*. *akari* by mites). Many other *Rickettsiae* are endosymbionts involved in various metabolic functions in their hosts (Abdad et al. 2018; Bonnet & Pollet 2021). For the pathogenic *Rickettsiae,* several vertebrate species could play the role of reservoir. Species of the SFG are more or less pathogenic in humans and responsible for rickettsioses, characterised by an eschar at the bite site and/or fever and headache, and sometimes with complications like cutaneous necrosis, pneumonia, endothelial damages and organ failure (Abdad et al. 2018).

In France, the *Rickettsia* species detected in questing *I*. *ricinus* ticks, when identified at the species level, are *R*. *felis*, *R*. *helvetica* and *R*. *monacensis* (**SI 15**). Data are too scarce to allow valid comparison of the prevalence between geographical regions (**Figure 9d**). DNA of *R*. *helvetica* was found in most regions examined, suggesting a distribution throughout France (**Figure 9b**) (Parola et al. 1998b; Davoust et al. 2012; Michelet et al. 2014; Moutailler et al. 2016b; Bonnet et al. 2017). When detected, observed prevalence was estimated between 1.1 and 14.3% in nymphs and between 1.6 and 25.0% in adults (Cotté et al. 2010; Michelet et al. 2014; Nebbak et al. 2019; Lejal et al. 2019a).. DNA of *R*. *felis* was detected in questing ticks from the Paris region (Essonne) and from north-eastern (Alsace region) France (**Figure 8a**) (Vayssier-Taussat et al. 2013; Lejal et al. 2019a; Lejal et al. 2019), whereas *R*. *monacensis* DNA was found in southern France (Hautes-Pyrénées; **Figure 8c**) (Akl et al. 2019). Tests to detect DNA of *Rickettsia* SFG were carried out on a large number of questing adult and nymphal ticks in northern-eastern France, but none were found positive (Michelet et al. 2014).

**Figure 9:**
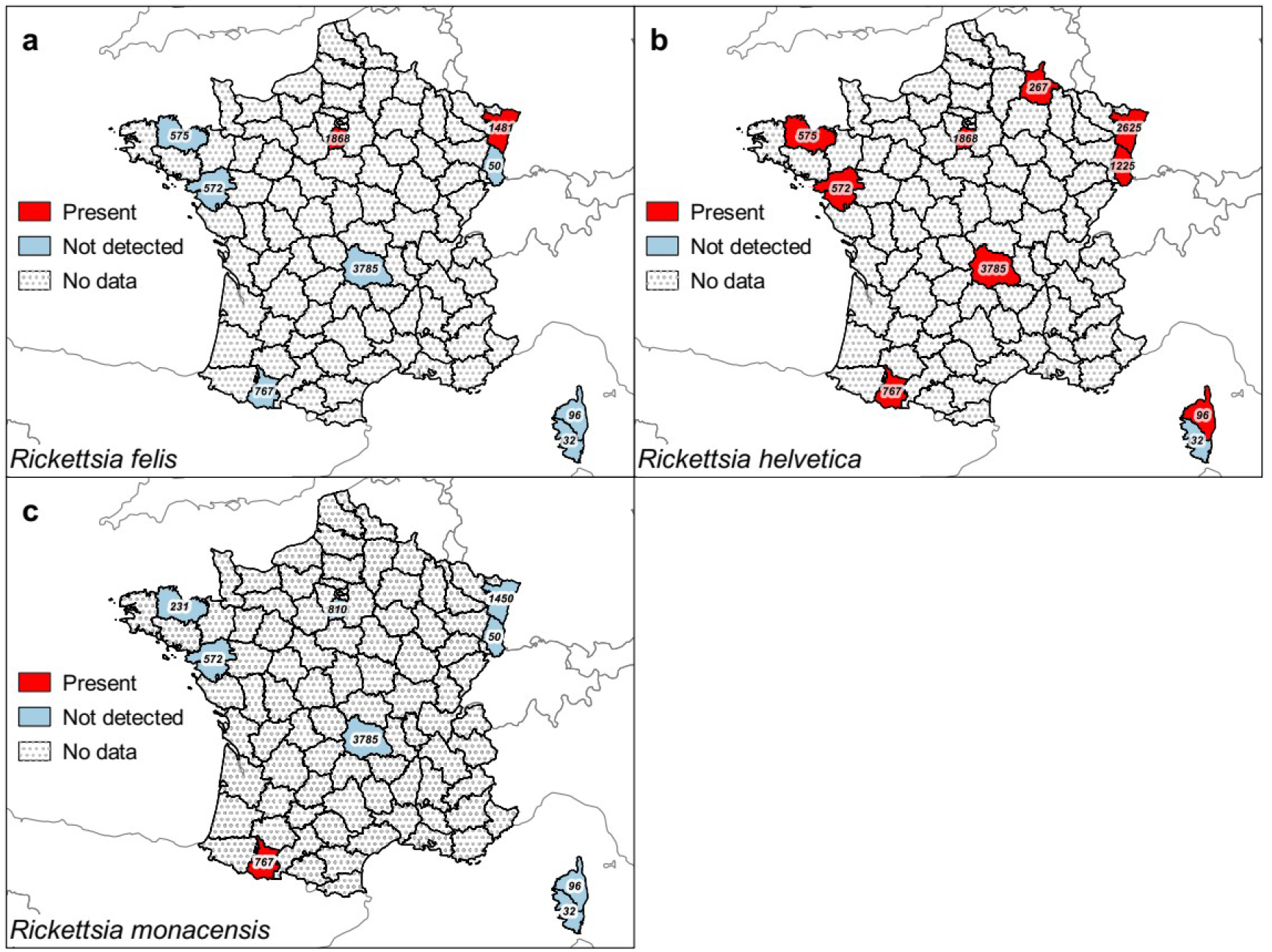
Presence maps of *Rickettsia felis*, and *R*. *helvetica*, and *R*. *monacensis* in questing and attached nymph and adult *Ixodes ricinus* ticks in European France by department. Detected presence by PCR methods and respective sampling effort per department in number of tested ticks (nymphs and adults) for *R*. *felis* (**a**), *R*. *helvetica* (**b**), and *R*. *monacensis* (**c**).

#### 3.3.10 The genus *Babesia*

*Babesia* sensu stricto belongs to a species-rich genus of unicellular eukaryotes of the Apicomplexa phylum. They are intra-erythrocyte parasites of vertebrates transmitted principally by ticks and generally characterised by transovarian transmission (Walter & Weber 1981; Schnittger et al. 2012; Vannier & Krause 2012; Jalovecka et al. 2019). *Babesia* species are responsible for babesiosis, worldwide tick-borne diseases that affect many mammalian species (mainly cattle, sheep, goat, equidae and canids) including humans as accidental hosts (Chauvin et al. 2009). The disease is characterized by hyperthermia, anaemia and haemoglobinuria and can led to death, particularly in immunosuppressed and splenectomised patients (Hildebrandt et al. 2021). Although tick bites are the main mode of transmission, infections can also occur by blood transfusion (Siński, Welc-Falęciak & Pogłód 2011; Fang & McCullough 2016). Transmission between dogs has been reported by dog bites for *B*. *gibsoni* and vertical transmission in dogs is suspected for *B*. *canis* and *B*. *vulpes* (formerly *Theileria annae* or “*B*. *microti*-like”; Baneth et al. 2019) (Solano-Gallego et al. 2016). *Babesia* species responsible for human infection in Europe are mainly *B*. *divergens* and *B*. *venatorum*, and exceptionally *B*. *microti* (Hildebrandt et al. 2021). Both *B*. *divergens*, for which the main reservoir host is cattle, and *B*. *venatorum* (formerly known as sp. EU1), for which the main reservoir hosts are cervids, are transmitted by *I*. *ricinus* (see review by Bonnet & Nadal 2021). *B*. *microti*, which infects rodents and shrews, is transmitted by other *Ixodes* species, specific of small mammals such as *I*. *trianguliceps*. The role of *I. ricinus* in its transmission remains unclear (Walter & Weber 1981; Gray et al. 2002; Bown et al. 2008).

Several studies have tested for the presence of zoonotic *Babesia* species in questing *I*. *ricinus* ticks from France (**Figure 9a**; **SI 16)**. *B*. *divergens* DNA was detected in questing *I*. *ricinus* ticks from north-eastern (Ardennes), the Paris region (Essonne), central (Saône-et-Loire) and western (Ille-et-Vilaine) France (**Figure 9c**) (Bonnet et al. 2013; Moutailler et al. 2016b; Paul et al. 2016; Jouglin et al. 2017b). *B*. *venatorum* DNA was detected in ticks from the Paris region (Essonne), and the north-eastern (Alsace region), southern (Haute-Garonne), and south-western (Haute-Garonne and Hautes-Pyrénées) regions of the country (**Figure 9d**) (Reis et al. 2011a; Michelet et al. 2014; Paul et al. (2016); Akl et al. 2019; Lejal et al. 2019a; Lejal et al. 2019b; Lebert et al. 2020). Differences in the distribution of the *Babesia* species could be explained by differences in the abundance and prevalence of infection of reservoir hosts in each studied area: *B*. *venatorum* was more often found in forested landscapes, whereas *B*. *divergens* occurs in more fragmented landscape with a high proportion of land cover dedicated to cattle breeding.

DNA of other *Babesia* species of medical and veterinary interest has also been examined in questing *I*. *ricinus*: *B. capreoli* that infects cervids (Young et al. 2019); *B. bovis*, *B. bigemina*, *B. major*, *B. occultans* and *B. ovata* which infect mainly cattle; *B. crassa*, *B. motasi* and *B. ovis which* infect mainly goats and sheep; *B*. *caballi* which infects equines; *B*. *canis*, *B*. *gibsoni*, *B*. *vogeli* and *B*. *vulpes* which mainly infect canids (Schnittger et al. 2012; Baneth et al. 2015; Solano-Gallego et al. 2016); and *B*. *microti* (which actually belongs to a separate phylogenetic clade; Schnittger et al. 2012). Among these, only *B. capreoli* was confirmed to be transmitted by *I. ricinus* ticks and its DNA was detected in questing ticks of this species sampled in western France (Ille-et-Vilaine) and in the Paris region (Essonne) (**Figure 9b**) (Jouglin et al. 2017b; Lejal et al. 2019b).

**Figure 9:**
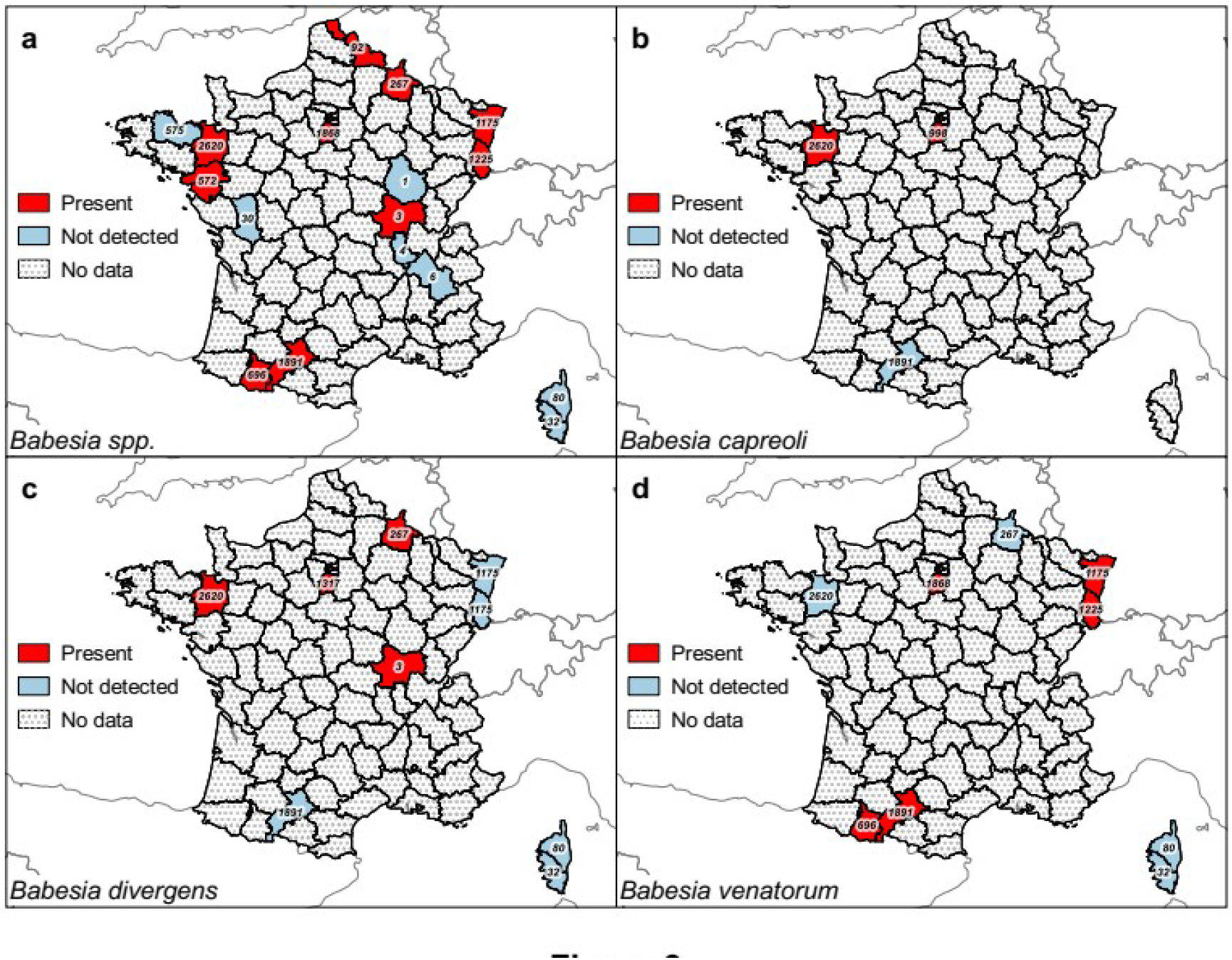
Presence maps of all *Babesia* spp., *B*. *capreoli*, *B*. *divergens* and *B*. *venatorum* in questing and attached nymph and adult *Ixodes ricinus* ticks in European France by department. Detected presence maps and respective sampling effort per department in number of tested ticks (nymphs and adults) for *Babesia* spp. (**a**), *B*. *capreoli* (**b**), *B*. *divergens* (**c**) and *B*. *venatorum* (**d)**.

#### 3.3.11 The genus *Theileria*

Although closely related, parasites from the *Theileria* genus differ from those of *Babesia* by a pre-erythrocyte stage in lymphocytes in the vertebrate host, and by the absence of transovarian transmission in the tick vector (Mans, Pienaar & Latif 2015). These parasites infect mainly ruminants in countries from the southern Hemisphere, where they are responsible for important economic loss (Mans, Pienaar & Latif 2015), and horses (*Theileria equi*) worldwide (Nadal, Bonnet & Marsot 2021). Parasites belonging to this genus are not known to infect humans.

In several studies, *I*. *ricinus* ticks sampled in France were examined for the presence of DNA from several *Theileria* species that are known to occur in Europe and around the Mediterranean basin (**SI 16**); none was detected. However, strains closely related to *T*. *parva* and *T*. *taurotragi* were found by high-throughput sequencing methods in questing *I*. *ricinus* nymphs sampled in north-eastern France (Alsace region), but no prevalence estimates were calculated (Bonnet et al. 2014). Interestingly, *T. equi* DNA was detected in adult *I. ricinus* that were questing or attached to piroplasm-free vertebrate hosts in Spain, Italy and the Netherlands (Butler et al. 2016; Garcia-Sanmartin et al 2008; Iori et al. 2010). These results highlight the need to study the vector competence of *I. ricinus* for *T. equi* and other *Theileria* species present in Europe to evaluate its potential contribution in the transmission of the pathogen to horses.

#### 3.3.12 Tick-borne encephalitis virus

Tick-borne encephalitis (TBE) was described for the first time in 1931 in Austria and the virus (TBEV) was isolated in 1937 in Russia (see review by Kaiser 2008 and Zlobin, Pogodina & Kahl 2017). This virus belongs to the *Flavivirus* genus of the Flaviviridae family (simple stranded RNA virus). Three main virus sub-types have been identified: the “European” sub-type, the “Far-Eastern” sub-type, and the “Siberian” sub-type. However, two other subtypes have been recently described: the Balkanian and the Himalayan sub-types (Ruzek et al. 2019). The main reservoir hosts are small mammals, among which *Apodemus* spp. mice seem to be particularly infectious for ticks (Labuda et al. 1993c; Labuda et al. 1996; Labuda et al. 1997; Knap et al. 2012). The virus causes meningitis and meningoencephalitis in humans, which can lead to neurological sequelae, such as paralysis, and death. The Far-Eastern subtype results in the most severe forms of the disease, with death in up to 2% of patients (Ruzek et al. 2019). TBEV is mainly transmitted by *I*. *ricinus* and *I*. *persulcatus* ticks in Europe (Mansfield et al. 2009). Human infections can occur not only by tick bites, but also by consumption of unpasteurised milk or dairy products from infected animals (Süss 2011). Transovarian transmission in ticks is weak (Danielova et al. 2002; Labuda et al. 1993a), but co-feeding transmission, where an infected tick infects non-infected ticks feeding nearby on the host, is of greater epidemiological importance (Labuda et al. 1993c; Labuda et al. 1997). Indeed, it was established that co-feeding transmission is of paramount importance in maintaining the virus in tick populations and thus its circulation in vertebrate hosts (Randolph, Gern & Nuttal 1996).

**Figure 10:**
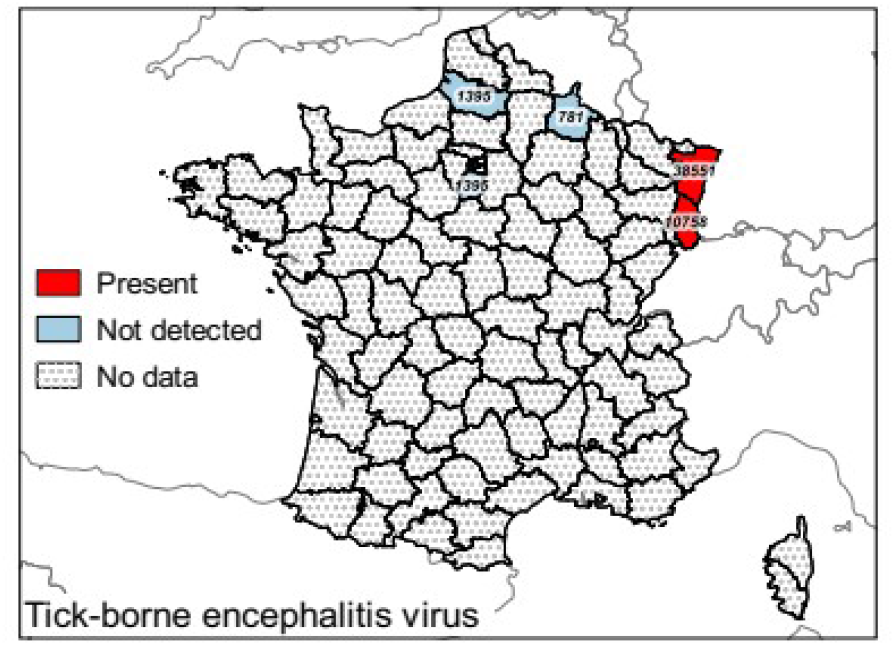
Presence map of tick-borne encephalitis virus (TBEV) in questing and attached nymph and adult *Ixodes ricinus* ticks in European France by department. Detected presence of TBEV by RT-PCR methods and sampling effort per department in number of tested ticks (nymphs and adults).

Only the “European” sub-type, considered as the least virulent, is present in western Europe (Bestehorn et al. 2018). In France, TBEV prevalence in questing *I*. *ricinus* ticks seems low (<0.1% in larvae and <1% in nymphs and adults) and is probably localised in micro-foci (Chatelain & Ardouin 1978; Pérez-Eid, Hannoun & Rodhain 1992; Bournez et al. 2020a). Too few studies have been conducted to conclude on the distribution of TBEV and its spatio-temporal variation in questing ticks in France, but it is likely present throughout the north-eastern part of the country, with well-identified endemic foci in the Alsace region (**Figure 10**; **SI 17**) (Pérez-Eid, Hannoun & Rodhain 1992; Hansmann et al. 2006; Rigaud et al. 2016; Bournez et al. 2020a). Serological results from wild ungulates suggest that the virus is also likely circulating in northern, north-eastern and eastern France, and that a closely related virus is present in the Pyrenees (Bournez et al. 2020b). Human TBE cases have been reported in western and central France (Rhône-Alpes and in Auvergne regions) during the last decade (Velay et al. 2018; Botelho-Nevers et al. 2019). In 2020, 43 TBE cases occurred in eastern France (Ain) following the consumption of goat cheese made with contaminated raw milk and thus identified a new focus (Beaufils et al. 2021; Gonzalez et al. 2022). The distribution of the virus seems to be changing in Europe, probably because of global changes that affect larvae and nymph co-occurrence on hosts and thus the probability of co-feeding transmission. However, this trend might be biased by a better detection of human cases and other socio-economical changes (Randoph et al. 2000; Randolph & Rogers 2000). Since 2021, TBE has been listed as a reportable disease in France, a measure that should improve our knowledge on human exposure.

#### 3.3.13 Other viruses

The Eyach virus is a *Coltivirus* that could be responsible for neurological damage, as seropositivity was associated with neuropathies in areas of the former Czechoslovakia (Malkova et al. 1980; Charrel et al. 2004). The virus was first isolated in Germany in 1972 from *I*. *ricinus* ticks and then again in western France in 1981 (Mayenne) from *I*. *ricinus* and *I*. *ventalloi* ticks collected from a European rabbit (Rehse-Küpper et al. 1976; Chastel et al. 1984). However, the vector competence of *I*. *ricinus* for this virus has not been formally demonstrated and the role of the European rabbit as reservoir remains unclear (Chastel 1998; Charrel et al. 2004). More recently, viral RNA was detected in questing *I*. *ricinus* ticks in north-eastern France; in the Alsace region in 2010 and in the Ardennes department in 2012 (Moutailler et al. 2016a).

## 4 Conclusions

This systematic map and narrative review assembled results of studies published in European France between 1965 and 2020 at various spatial scales (country-wide to site-specific) and exploring the influence of different variables (meteorological, biogeographic zones, landscape features, host density, local vegetation, soil characteristics) on the density of questing *I*. *ricinus.* It also provided summary information on the prevalence and distribution of several tick-borne pathogens detected in *I. ricinus* ticks in the country.

The first studies on the distribution of ixodid tick species in France started in the 60’s (Lamontellerie 1965; Panas et al. 1976). Gilot and colleagues then started to examine the association of *I*. *ricinus* with different vegetation types and its related phenology (Gilot, Pautou & Mancada 1975; Gilot et al. 1975). After that, more standardised studies on the distribution and phenology of the species were performed (e.g., Gilot et al. 1989; Gilot et al. 1994). More recent work has focused on identifying landscape, vegetation and microclimatic factors associated with the abundance and phenology of the species using standardised methods and new technologies (e.g., via Geographic Information Systems). The advent of PCR technologies also enabled researchers to examine the present of associated pathogens. These different studies have thus made it possible to accumulate a certain amount of knowledge about *I*. *ricinus* in France, but no synthesis of these data was available. This study aimed to fill this void and to highlight remaining knowledge gaps on the distribution, phenology, and host range of *I. ricinus* and on the prevalence of its associated pathogens in certain regions of France.

For example, larval phenology is still poorly described overall, and larval-nymphal phenology and host use under Mediterranean climates remain to be investigated. More longitudinal studies on *I. ricinus* phenology that include multiple environmental parameters are also necessary to better define factors essential for the development of the species and to predict acarological risk, particularly in a context of climate change. Very few such studies have been conducted in France so far (Paul et al. 2016; Wongnak et al. 2022).

Studies on tick host use has focused mainly on Rodentia (mice and voles) and wild Cetartiodactyla (e.g., roe deer), whereas the role of domestic Cetartiodactyla (cattle, sheep, and goats), Perissodactyla (horses and donkey), Carnivora (e.g. cats, dogs, foxes), Aves (birds), Eulipotyphla (shrews and hedghogs), Lagomorpha (e.g., rabbits and hares), and Squamata (e.g. lizards and snakes) remain poorly quantified. It should also be noted that the few studies that exist are rarely comparable (different location, season and/or protocol), preventing direct comparisons of tick burdens between species or between different climates or environments for a same species. A standardisation of protocols is thus called for.

Studies of tick-borne pathogens have focused mainly on *B*. *burgdorferi* s.l. In addition, due to logistical constraints, these studies have often concentrated around locations of the investigating research units and, more distant or isolated regions have therefore received less attention, leading to a lack of data to inform and prevent tick-borne disease in some local populations (Septfons et al. 2018). This highlights the need to scale studies according to expected outcomes, to ensure for instance that the targeted area lies within the expected domain of the desired model. That said, the data available on *B*. *burgdorferi* s.l. were sufficient to compare prevalence and the occurrence of different genospecies in questing ticks among regions. The observed gradients mirror observations at the European scale: an increasing prevalence along a SW-NE gradient with an increasing proportion of prevalence of *B*. *afzelii* (Strnad et al. 2017). These prevalence gradients are likely driven by biogeographic gradients in Europe and the evolutionary history of *Borrelia* genospecies (Margos et al. 2011). The distribution and the spatial variation in prevalence of other tick-borne pathogens are still poorly known and deserve further study. Due to the veterinary importance of babesiosis, the fact that several species can infect humans, the numerous side effects of babesicidal drugs, and the absence of a vaccine against most of the species that occur in France (only a vaccine against *B*. *canis* exists, to our knowledge: Pirodog, Boehringer Ingelheim Animal Health, France), it is particularly important to intensify research and monitoring of these parasites. Likewise, recent distributional changes in TBEV exposure should be more closely tracked to anticipate prevention. For example, understanding the potential role of birds as dispersers of infected ticks for the geographic expansion of the virus would be extremely informative (Wilhelmsson et al. 2020).

Detection methods of infectious agents in ticks have rapidly improved over the last decades, particularly with the development of high throughput sequencing methods and more sensitive PCR techniques (i.e., quantitative PCR, microfluidic PCR, digital PCR). That said, it is important to remember that the detection of pathogen DNA in an arthropod does not imply that the arthropod is a vector of this organism, it only indicates that a blood meal was taken on an infected host. The formal demonstration of vector competence requires experimental study (Khal et al. 2002; Bonnet & Nadal 2021). Although the vector competence of *I*. *ricinus* has been experimentally demonstrated for many important infectious agents, its role in the transmission of other agents such as *C. burnetii*, *F*. *tularensis*, *F. philomiragia*, and *Rickettsia* spp. still requires experimental validation. Reciprocally, other species can be the vectors of some pathogens transmitted by *I*. *ricinus* and their role in the zoonotic cycle requires explicit consideration. For instance, few tick species have been experimentally assessed for vector competence for *B*. *burgdorferi* s.l. (Eisen 2020), although these bacteria have been frequently detected in *I*. *acuminatus* (Doby et al. 1990; Szekeres et al. 2015), *I*. *arboricola* (Spitalska et al. 2011; Heylen et al. 2013), *I*. *frontalis* (Heylen et al. 2013; Heylen et al. 2017a; Heylen et al. 2017b; Palomar et al. 2017), *I*. *hexagonus* (Gern et al. 1991; Geurden et al. 2018) and *I*. *trianguliceps* (Doby et al. 1990; Nefedova et al. 2005). Knowledge on vector diversity and the relative roles of different vector species is essential for understanding the distribution and observed variation in pathogen prevalence and to establish pertinent prevention measures. The role of other co-infecting tick microorganisms, symbiotic or commensal, in the ecology, physiology and vector competence of *I*. *ricinus* also needs to be investigated (Bonnet & Pollet 2021; Lejal et al. 2021). Such studies could uncover new ways of controlling ticks and the pathogens they transmit.

To conclude, more studies are needed to better assess the distribution of *I*. *ricinus* and its associated pathogens in France, particularly in the context of current global changes that could see a shift in the distribution of native species, the emergence of tropical species in Europe and changes in tick phenology and transmission dynamics. Efforts focused on a better understanding of tick-borne pathogen ecology is also necessary to better evaluate acarological risk. Indeed, human cases of Lyme borreliosis have increased over the last two decades for diverse reasons (e.g., awareness of clinicians, increasing outdoor recreational activities, modifications of the tick ecosystems, etc.) and new tick-borne diseases have emerged like relapsing fever associated with *B. miyamotoi* or *N. mikurensis* (Figoni et al. 2019). Current participative science projects like CITIQUE (https://www.citique.fr/), based on voluntary declarations of tick-bites and tick collection, may fulfil some of these gaps in the future. For example, data collected under these programs should help complete the distribution map of *I. ricinus* and its associated pathogens and should provide information on high human exposure periods (Eisen & Eisen 2021). More generally, an integrated research programme that links data and knowledge from diverse sources would help to develop more efficient strategies to control tick populations and reduce tick-borne disease risk.

## Supporting information

version 1

## Acknowledgments

The author thanks Gilles Bourgoin, Sébastien Grech-Angelini, Sara Moutailler, Frank Boué, Mathilde Gondard, Guy Joncour, Lorraine Michelet, Gérald Umhang, Muriel Vayssier-Taussat, Olivier Plantard, Alber Agoulon, Thomas Pollet, Karine Chalvet-Montfray, Émilie Lejal, Maud Marsot, Frédéric Stachurski and Laurence Vial for sharing their raw data on the presence of *I*. *ricinus* (Figure 2).

## Data and code availability

All encoded and compiled data are available as supplementary information at https://doi.org/10.1101/2023.04.18.537315.

## Authors’ contributions

GP and SB designed the study. GP conducted the literature search, screened the references, developed the bibliographic database and compiled all the data, wrote the first draft of the manuscript and produced all figures and tables. BL gave valuable methodological recommendations. LB, NB, KDM, JF, BL, EQ, MRM and SB discussed data interpretation and presentation and edited the manuscript, figures and tables.

