## Supplementary material for "The distribution, phenology, host range and pathogen prevalence of *Ixodes ricinus* in France: a systematic map and narrative review": SI 1

**Supplementary information 1: Exact search lines**

For the literature searches in CAB Direct, JSTOR, ScienceDirect, Scopus, Pascal & Francis, PubMed and WorldCat database the searches were conducted in three runs.

Search run A was conducted with:

"**Ixodes ricinus**" AND (**abundance** OR **activity** OR **density** OR **distribution** OR **dynamics**)

Search run B was conducted with:

"**Ixodes ricinus**" AND (**presence** OR **questing** OR **behaviour** OR **behavior** OR **dispersal**)

Search run C was conducted with:

"**Ixodes ricinus**" AND (**prevalence** OR **burden** OR **infestation**)

**Dates (dd/mm/yyyy) of searches per run**

|  | Date of run A | Date of run B | Date of run C |
| --- | --- | --- | --- |
| CAB Direct | 12/12/2019 | 12/12/2019 | 12/12/2019 |
| JSTOR | 14/11/2019 | 11/12/2019 | 12/12/2019 |
| ScienceDirect | 12/12/2019 | 12/12/2019 | 12/12/2019 |
| Scopus | 14/11/2019 | 11/12/2019 | 12/12/2019 |
| Pascal & Francis | 27/11/2019 | 12/12/2019 | 12/12/2019 |
| PubMed | 12/12/2019 | 12/12/2019 | 12/12/2019 |
| Worldcat | 18/11/2019 | 12/12/2019 | 16/12/2019 |

Searches in Agricola (<https://agricola.nal.usda.gov/>), Open Access Thesis and Dissertations (<https://oatd.org/>), Thèses vétérinaires (available at <http://scbev.vetagro-sup.fr/rechcat/rechcat.htm>) and theses.fr (<http://www.theses.fr/>) were conducted with only “**Ixodes ricinus**” in the search line during the same period.

This is a supplementary material for “Perez, G., Bournez, L., Boulanger, N., Fite, J., Livoreil, B., McCoy, K. D., Quillery, E., René-Martellet, M., and Bonnet, S. I. The distribution, phenology, host range and pathogen prevalence of *Ixodes ricinus* in France: a systematic map and a narrative review”, a preprint recommended in PCI Infections <https://doi.org/10.24072/pci.infections.100076>.
