## Supplementary material for "The distribution, phenology, host range and pathogen prevalence of *Ixodes ricinus* in France: a systematic map and narrative review": SI 2

**Supplementary information 2: Test list of references used to evaluate the literature search efficiency**

* References not found in the literature search

This is a supplementary material for “Perez, G., Bournez, L., Boulanger, N., Fite, J., Livoreil, B., McCoy, K. D., Quillery, E., René-Martellet, M., and Bonnet, S. I. The distribution, phenology, host range and pathogen prevalence of *Ixodes ricinus* in France: a systematic map and a narrative review”, a preprint recommended in PCI Infections <https://doi.org/10.24072/pci.infections.100076>.
