## Supplementary material for "The distribution, phenology, host range and pathogen prevalence of *Ixodes ricinus* in France: a systematic map and narrative review": SI 3

**Supplementary information 3: Table of the inclusion and exclusion criteria.**

|  | **Inclusion criteria** | **Exclusion criteria** |
| --- | --- | --- |
| **Date / Peer-review** | Peer-reviewed articles, book sections, veterinary or university theses published at any time prior to December 16th, 2019. | Other kind of documents (e.g., conference proceedings), non-peer-reviewed articles and articles published after December 31th, 2020. |
| **Population** | *I. ricinus* ticks at all stages | Other tick species, search of tick-borne pathogens in vertebrate hosts only. |
| **Context** | All references including at least one sample from European France (mainland France, Corsica island and other French islands of Mediterranean Sea, Bay of Biscay, Celtic Sea, English Channel, and North Sea were retained). | No specific sampling location or samples from outside European France including overseas French territories. |
| **Comparator** | Temporal comparisons, including inter-annual (i.e., dynamics), intra-annual (i.e., seasonal variation in questing tick density), and daily comparisons (i.e., abundance in function of day hour), and spatial comparisons at multiple scales: micro-habitats, habitats, landscapes and macro-scale. It also included comparisons between vegetation management types, levels and/or types of anthropic activities, vertebrate host densities and physical, chemical or biological treatments of the environment or the host. | No comparator was excluded. |
| **Outcomes** | References directly related to the ecology of *I*. *ricinus* and/or studies on tick-borne pathogens in *I*. *ricinus* ticks. | References not directly related to the ecology of *I. ricinus*. For example, those that focused on cellular or molecular aspects of *I. ricinus* biology or on tick-borne pathogens alone, or laboratory-based studies of *I. ricinus* – tick-borne pathogens interactions. Studies with tick-borne pathogens in vertebrate hosts only. |
| **Language** | Articles written in English or French. | Languages other than English or French. |

This is a supplementary material for “Perez, G., Bournez, L., Boulanger, N., Fite, J., Livoreil, B., McCoy, K. D., Quillery, E., René-Martellet, M., and Bonnet, S. I. The distribution, phenology, host range and pathogen prevalence of *Ixodes ricinus* in France: a systematic map and a narrative review”, a preprint recommended in PCI Infections <https://doi.org/10.24072/pci.infections.100076>.
