## Supplementary material for "The distribution, phenology, host range and pathogen prevalence of *Ixodes ricinus* in France: a systematic map and narrative review": SI 5

**Supplementary information 5: Final corpus of references**

Agoulon, A., Malandrin, L., Lepigeon, F., Vénisse, M., Bonnet, S., Becker, C.A.M.M., Hoch, T., Bastian, S., Plantard, O., Beaudeau, F., 2012. A vegetation index qualifying pasture edges is related to *Ixodes ricinus* density and to *Babesia divergens* seroprevalence in dairy cattle herds. Vet. Parasitol. 185, 101–109. https://doi.org/10.1016/j.vetpar.2011.10.022

Agoulon, A., Hoch, T., Heylen, D., Chalvet-Monfray, K., Plantard, O., 2019. Unravelling the phenology of *Ixodes frontalis*, a common but understudied tick species in Europe. Ticks Tick. Borne. Dis. 10, 505–512. https://doi.org/10.1016/j.ttbdis.2018.12.009

Akl, T., Bourgoin, G., Souq, M.-L., Appolinaire, J., Poirel, M.-T., Gibert, P., Abi Rizk, G., Garel, M., Zenner, L., 2019. Detection of tick-borne pathogens in questing *Ixodes ricinus* in the French Pyrenees and first identification of *Rickettsia monacensis* in France. Parasite 26, 9 p. https://doi.org/10.1051/parasite/2019019

Anderson, J.F., Doby, J.M., Coutarmanac’h, A., Hyde, F.W., Johnson, R.C., 1986. Antigenically different *Borrelia burgdorferi* strains from *Ixodes ricinus* in Brittany, France. Med. Mal. Infect. 16, 171–175. https://doi.org/10.1016/S0399-077X(86)80221-9

Aubert, M.F., 1983. Contribution à l’étude des ectoparasites du renard et d’autres carnivores sauvages de l’Est de la France et à l’étude du rôle des arthropodes dans l’épizootie de la rage vulpine.

Aubert, M.F.., 1975. Contribution a l’étude du parasitisme du renard (*Vulpes vulpes* L.) par les Ixodidae (Acarina) dans le Nord-Est de la France interprétation de la dynamique saisonnière des parasites en relation avec la biologie de l’hôte. Acarologia 53, 383–415. https://doi.org/10.1051/acarologia/20142144

Bestehorn, M., Weigold, S., Kern, W. V, Chitimia-dobler, L., Mackenstedt, U., Dobler, G., Id, J.P.B., 2018. Phylogenetics of Tick-Borne Encephalitis virus in endemic foci in the upper Rhine region in France and Germany. PLoS One 13, e0204790 (10 p). https://doi.org/https://doi.org/10.1371/journal. pone.0204790

Beytout, J., George, J.C., Malaval, J., Garnier, M., Beytout, M., Baranton, G., Ferquel, E., Postic, D., 2007. Lyme borreliosis incidence in two French departments: Correlation with infection of *Ixodes ricinus* ticks by *Borrelia burgdorferi* sensu lato. Vector-Borne Zoonotic Dis. 7, 507–517. https://doi.org/10.1089/vbz.2006.0633

Bineau, S., 2009. La tique dure *Ixodes ricinus* dans le marais Breton-Vendéen : étude de sa distribution spatio-temporelle et des facteurs influant sur sa présence. DVM thesis. Université de Nantes, France.

Bonnet, S., De La Fuente, J., Nicollet, P., Liu, X., Madani, N., Blanchard, B., Maingourd, C., Alongi, A., Torina, A., Fernández De Mera, I.G., Vicente, J., George, J.-C., Vayssier-Taussat, M., Joncour, G., 2013. Prevalence of tick-borne pathogens in adult *Dermacentor* spp. ticks from nine collection sites in France. Vector-Borne Zoonotic Dis. 13, 226–236. https://doi.org/10.1089/vbz.2011.0933

Bonnet, S.I., Paul, R.E.L., Bischoff, E., Cote, M., Le Naour, E., 2017. First identification of *Rickettsia helvetica* in questing ticks from a French Northern Brittany forest. PLoS Negl. Trop. Dis. 11, e0005416 (10 p). https://doi.org/10.1371/journal.pntd.0005416

Bonnet, S., Choumet, V., Masseglia, S., Cote, M., Ferquel, E., Lilin, T., Marsot, M., Chapuis, J.L., Vourc’h, G., 2015. Infection of Siberian chipmunks (*Tamias sibiricus barberi*) with *Borrelia* sp. reveals a low reservoir competence under experimental conditions. Ticks Tick. Borne. Dis. 6, 393–400. https://doi.org/10.1016/j.ttbdis.2015.03.008

Bonnet, S., Michelet, L., Moutailler, S., Cheval, J., Hébert, C., Vayssier-Taussat, M., Eloit, M., 2014. Identification of parasitic communities within European ticks using next-generation sequencing. PLoS Negl. Trop. Dis. 8, e2753 (6 p). https://doi.org/10.1371/journal.pntd.0002753

Bord, S., Druilhet, P., Gasqui, P., Abrial, D., Vourc’h, G., 2014. Bayesian estimation of abundance based on removal sampling under weak assumption of closed population with catchability depending on environmental conditions. Application to tick abundance. Ecol. Modell. 274, 72–79. https://doi.org/10.1016/j.ecolmodel.2013.12.004

Bord, S., 2014. Bayesian estimation of abundance based on removal sampling with variability of the sampling: application to host-questing *Ixodes ricinus* ticks. PhD thesis. Université Blaise-Pascale, Clermont-Ferrand II, France.

Boulanger, N., Boyer, P., Talagrand-Reboul, E., Hansmann, Y., 2019. Ticks and tick-borne diseases. Médecine Mal. Infect. 49, 87–97. https://doi.org/10.1016/j.medmal.2019.01.007

Boulanger, N., Zilliox, L., Goldstein, V., Boyer, P., Napolitano, D., Jaulhac, B., 2018. Surveillance du vecteur de la borréliose de Lyme, *Ixodes ricinus*, en Alsace de 2013 à 2016. Bull Epidemiol Hebdo 19–20, 400–405.

Bournez, L., Umhang, G., Moinet, M., Richomme, C., Demerson, J., Caillot, C., Devillers, E., Boucher, J., 2020. Tick-borne encephalitis virus: seasonal and annual variation of epidemiological parameters related to nymph-to-larva transmission and exposure of small mammals. Pathogens 9, e518. https://doi.org/10.3390/pathogens9070518

Boyard, C., Barnouin, J., Gasqui, P., Vourc’h, G., 2007. Local environmental factors characterizing *Ixodes ricinus* nymph abundance in grazed permanent pastures for cattle. Parasitology 134, 987–994. https://doi.org/10.1017/S0031182007002351

Boyard, C., 2007. Environmental factors influencing the variation of *Ixodes ricinus* tick abundance in two model zones in Auvergne (central France). PhD thesis. Université Blaise Pascal - Clermont-Ferrand II 2007-12-18, France.

Boyard, C., Barnouin, J., Bord, S., Gasqui, P., Vourc’h, G., 2011. Reproducibility of local environmental factors for the abundance of questing *Ixodes ricinus* nymphs on pastures. Ticks Tick. Borne. Dis. 2, 104–110. https://doi.org/10.1016/j.ttbdis.2011.02.001

Boyard, C., Vourc’h, G., Barnouin, J., 2008. The relationships between *Ixodes ricinus* and small mammal species at the woodland-pasture interface. Exp. Appl. Acarol. 44, 61–76. https://doi.org/10.1007/s10493-008-9132-3

Boyer, N., Réale, D., Marmet, J., Pisanu, B., Chapuis, J.-L., 2010. Personality, space use and tick load in an introduced population of Siberian chipmunks *Tamias sibiricus*. J. Anim. Ecol. 79, 538–547. https://doi.org/10.1111/j.1365-2656.2010.01659.x

Boyer, P.H., Boulanger, N., Nebbak, A., Collin, E., Jaulhac, B., Almeras, L., 2017. Assessment of MALDI-TOF MS biotyping for *Borrelia burgdorferi* sl detection in *Ixodes ricinus*. PLoS One 12, e0185430 (16 p). https://doi.org/10.1371/journal.pone.0185430

Boyer, P.H., De Martino, S.J., Hansmann, Y., Zilliox, L., Boulanger, N., Jaulhac, B., 2017. No evidence of *Borrelia mayonii* in an endemic area for Lyme borreliosis in France. Parasites and Vectors 10, 10:282. https://doi.org/10.1186/s13071-017-2212-7

Carrer, H., 2007. Etude de la prévalence de *Babesia divergens* chez la tique *Ixodes ricinus*. DVM thesis. Université de Nantes, France.

Cat, J., 2017. Integrating the effects of weather factors in the modeling of the survival and activity in *Ixodes ricinus* tick populations in the context of climate change. PhD thesis. Clermont Auvergne, France.

Cat, J., Beugnet, F., Hoch, T., Jongejan, F., Prangé, A., Chalvet-Monfray, K., 2017. Influence of the spatial heterogeneity in tick abundance in the modeling of the seasonal activity of *Ixodes ricinus* nymphs in Western Europe. Exp. Appl. Acarol. 71, 115–130. https://doi.org/10.1007/s10493-016-0099-1

Chastagner, A., 2014. Etude des cycles épidémiologiques d’*Anaplasma phagocytophilum* en France : apport des approches de caractérisation génétique. PhD thesis. Université Blaise Pascal - Clermont-Ferrand II, France.

Chastagner, A., Bailly, X., Leblond, A., Pradier, S., Vourc’H, G., 2013. Single genotype of *Anaplasma phagocytophilum* identified from ticks, Camargue, France. Emerg. Infect. Dis. 19, 825–827. https://doi.org/10.3201/eid1905.121003

Chastagner, A., Pion, A., Verheyden, H., Lourtet, B., Cargnelutti, B., Picot, D., Poux, V., Bard, É., Plantard, O., McCoy, K.D., Leblond, A., Vourc’h, G., Bailly, X., 2017. Host specificity, pathogen exposure, and superinfections impact the distribution of *Anaplasma phagocytophilum* genotypes in ticks, roe deer, and livestock in a fragmented agricultural landscape. Infect. Genet. Evol. 55, 31–44. https://doi.org/10.1016/j.meegid.2017.08.010

Chastel, C., 1998. Erve and Eyach: two viruses isolated in France, neuropathogenic for man and widely distributed in Western Europe. Bull. Acad. Natl. Med. 182, 801–810.

Chastel, C., Main, A.J., Couatarmanac’h, A., Le Lay, G., Knudson, D.L., Quillien, M.C., Beaucournu, J.C., 1984. Isolation of Eyach virus (Reoviridae, Colorado tick fever group) from *Ixodes ricinus* and *I. ventalloi* ticks in France. Arch. Virol. 82, 161–171. https://doi.org/10.1007/BF01311160

Chatelain, J., Ardoin, P., 1978. Some demographic data from *Ixodes ricinus* Linné: vector of a virus strain of central european encephalitis in Alsace. Rev. Epidemiol. Sante Publique 26, 211–223.

Chatelain, J., Hannoun, C., Rodhain, F., 1979. Ecology of indigenous arbovirus in Alsace: tick central European encephalitis. Rev. Epidemiol. Sante Publique 27, 277–299.

Cicculli, V., Capai, L., Quilichini, Y., Masse, S., Fernandez-Alvarez, A., Minodier, L., Bompard, P., Charrel, R., Falchi, A., 2019. Molecular investigation of tick-borne pathogens in ixodid ticks infesting domestic animals (cattle and sheep) and small rodents (black rats) of Corsica, France. Ticks Tick. Borne. Dis. 10, 606–613. https://doi.org/10.1016/j.ttbdis.2019.02.007

Coiffait, C., 2019. Rhythm of the activity of the tick *Ixodes ricinus*: a synchronisation with roe deer’s activity? DVM thesis. 2019., France.

Cosson, J.-F., 2018. Ecology of Lyme disease. Rev. For. française 70, 185–203. https://doi.org/10.3917/spub.190.0073

Cosson, J.F., Michelet, L., Chotte, J., Le Naour, E., Cote, M., Devillers, E., Poulle, M.L., Huet, D., Galan, M., Geller, J., Moutailler, S., Vayssier-Taussat, M., 2014. Genetic characterization of the human relapsing fever spirochete *Borrelia miyamotoi* in vectors and animal reservoirs of Lyme disease spirochetes in France. Parasites and Vectors 7, 233 (5 p). https://doi.org/10.1186/1756-3305-7-233

Cotté, V., Bonnet, S., Cote, M., Vayssier-Taussat, M., 2010. Prevalence of five pathogenic agents in questing *Ixodes ricinus* ticks from western France. Vector-Borne Zoonotic Dis. 10, 723–730. https://doi.org/10.1089/vbz.2009.0066

Davoust, B., Socolovschi, C., Revelli, P., Gibert, P., Marié, J. Lou, Raoult, D., Parola, P., 2012. Detection of *Rickettsia helvetica* in *Ixodes ricinus* ticks collected from Pyrenean chamois in France. Ticks Tick. Borne. Dis. 3, 387–388. https://doi.org/10.1016/j.ttbdis.2012.10.009

Degeilh, B., 2007. Scientific basis for prevention. Med. Mal. Infect. 37, 360–367. https://doi.org/10.1016/j.medmal.2006.01.031

Degeilh, B., Guiguen, C., Gilot, B., 1999. A study of the distribution of *Ixodes ricinus*/*Borrelia burgdorferi* in Brittany: infectious risk for humans. Ann. La Soc. Entomol. Fr. 35, 534–539.

Degeilh, B., Pichot, J., Gilot, B., Allen, S., Doche, B., Guiguen, C., 1996. Dynamics of human primary-infestation with Lyme borreliosis (Erythema Chronicum Migrans) and dynamics of vector tick populations (*Ixodes ricinus* Linnaeus, 1758): attempt at correlation. Acarologia 37, 7–17.

Degeilh, B., 1996. Anthropozoonoses and hematophagous arthropods: ecoepidemiology of Lyme borreliosis; contribution to the study of its main vector, *Ixodes ricinus*, in the Atlantic area of influence, in mainland France. PhD thesis. France.

Degeilh, P., Guiguen, C., Gilot, B., Doche, B., Pichot, J., Beaucournu, J.-C., 1994. Distribution of *Ixodes ricinus* (Linne, 1758) (Acarina: Ixodidae) in the forest regions of the Massif Armoricain. Acarologia 35, 327–334.

Deprez, V., 2009. Les tiques dures autres qu’*Ixodes ricinus* dans le marais Breton-Vendéen : étude de leur distribution spatio-temporelle et des facteurs influant sur leur présence. DVM thesis. [S.l.] : [s.n.], 2009., France.

Deram, G., 2012. Exposition des bovins aux maladies transmises par les tiques et pratiques de pâturage : enquête dans 15 élevages d’Ille-et-Vilaine. DVM thesis. Université de Nantes, France.

Diarra, O., 1992. Contribution à l’étude de l’infestation des bovins par *Ixodes ricinus*. DVM thesis. Université de Nantes, France.

Dietrich, F., Schmidgen, T., Maggi, R.G., Richter, D., Matuschka, F.R., Vonthein, R., Breitschwerdt, E.B., Kempf, V.A.J., 2010. Prevalence of *Bartonella henselae* and *Borrelia burgdorferi* sensu lato DNA in *Ixodes ricinus* ticks in Europe. Appl. Environ. Microbiol. 76, 1395–1398. https://doi.org/10.1128/AEM.02788-09

Doby, J.M., Lemblé, C., Bigaignon, G., Kremer, M., Rolland, C., Lambert, M.C., 1990. Borrelia burgdorferi, agent of tick spirochaetoses (Lyme disease and other clinical forms) in *Ixodes ricinus* (Acari: Ixodidae) in Alsace. A study of 2223 ticks. Bull. la Société Française Parasitol. 8, 339–350.

Doby, J.M., Anderson, J.F., Couatarmanac’h, A., 1985. Observation of spirochetes in ticks from Britanny (preliminary note). Med. Mal. Infect. 15, 556–557. https://doi.org/10.1016/S0399-077X(85)80221-3

Doby, J.M., Bigaignon, G., 1992. Survey of the infestation level of the *Ixodes ricinus* tick by Borrelia burgdorferi. Complementary report. Bull. la Société Pathol. Exot. 85, 322–323.

Doby, J.M., Bigaignon, G., Degeilh, B., Guiguen, C., 1994. Ectoparasites of large wild mammals (deer and wild boars) and Lyme borreliosis. Search for *Borrelia burgdorferi* in more than 1400 ticks, lice, Pupipara Diptera and fleas. Rev. Med. Vet. (Toulouse). 145, 743–748.

Doby, J.M., Bigaignon, G., Launay, H., Costil, C., Lorvellec, O., 1990. Presence of *Borrelia burgdorferi*, agent of tick spirochaetosis, in *Ixodes* (*Exopalpiger*) *trianguliceps* Birula, 1895 and *Ixodes* (*Ixodes*) *acuminatus* Neumann, 1901 (Acari: Ixodidae) and in *Ctenophthalmus baeticus arvernus* Jordan, 1931 and *Megabothris turbidus* (Rothschild, 1909)(Insecta: Siphonaptera). Bull. la Société Française Parasitol. 8, 311–322. Bull. la Société Française Parasitol. 8, 311–322.

Doby, J.M., Bigaignon, G., Lorvelec, O., Imbert, G., 1991. Survey during 4 years of the infestation level of the tick *Ixodes ricinus* (Acari Ixodidae) by *Borrelia burgdorferi*, the agent of Lyme borreliosis, in 2 forests in Brittany. Bull. la Société Pathol. Exot. 84, 398–402.

Doby, J.M., Chevrier, S., Couatarmanac’h, A., Imbert-Hameurt, C., 1989. Infection of *Ixodes ricinus* (Acarina: Ixodidae) by *Borrelia burgdorferi*, agent of tick-spirochetosis (Lyme disease and other clinical forms) in the West of France. II - Detailed results and comments. Bull. la Soc. Fr. Parasitol. 7, 277–289.

Doby, J.M., Imbert-Hameurt, C., Jeanne, B., Chevrier, S., 1989. Infection of *Ixodes ricinus* (Acarina: Ixodidae) by *Borrelia burgdorferi*, agent of tick-spirochetosis (Lyme disease and other clinical forms) in the West of France. I: Overall results of the examination of 2320 ticks. Bull. la Soc. Fr. Parasitol. 7, 111–125.

Doby, J.M., van Laere, G., 1993. *Hunterellus hookeri* howard, 1907, hymenoptera chalcididae parasitizing *Ixodes ricinus* tick in the western and middle parts of France. Bull. la Société Française Parasitol. 11, 265–270.

Doby, J.M., Degeilh, B., 1996. Lyme borreliosis and urban and periurban public parks. Bull. la Société des Sci. Nat. l’Ouest la Fr. 18, 34–40.

Doby, J.-M., Bigaignon, G., Aubert, M., Imbert, G., 1991. Ectoparasites of the fox and Lyme borreliosis. Research on *Borrelia burgdorferi* in ixodid ticks and fleas (Siphonaptera). Bull. la Soc. Fr. Parasitol. 9, 279–288.

Doby, J.M., Bigaignon, G., Degeilh, B., 1992. Compared potential importance of field-mouse (*Apodemus sylvaticus*) and bank vole (*Clethrionomys glareolus*) in Lyme borreliosis epidemiology in forest in the western part of France, through relations between rodents and Ixodes ricinus tick. Bull. la Société Française Parasitol. 10, 271–293.

Doby, J.-M., Bigaignon, G., Doby-Dubois, M., Brumpt, M., Charmot, M., 1995. Risk of contamination by *Borrelia burgdorferi* sensu lato in a forest environment. Survey during 13 months of abundance of the tick *Ixodes ricinus* and its level of infestation by the Lyme borreliosis agent in Brittany. Bull. la Société Pathol. Exot. 88, 61–65.

Doby, J.M., Bigaignon, G., Godfroid, E., Bollen, A., Murga-Gutierrez, N., 1993. Frequency of *Borrelia burgdorferi*, agent of Lyme Borreliosis, in nymphs of *Ixodes ricinus* (Acari: Ixodidae) caught in some forests in the central part of France. Bull. la Soc. Fr. Parasitol. 11, 163–168.

Doche, B., Gilot, B., Degeilh, B., Pichot, J., Guiguen, C., 1993. Utilisation de l’indicateur végétal pour la cartographie d’une tique exophile à l’échelle de la France : exemple d’*Ixodes ricinus* (Linné, 1758), vecteur de la Borréliose de Lyme. Ann. Parasitol. Hum. Comparée 68, 188–195. https://doi.org/10.1051/parasite/1993684188

Doitrand, B., 1991. Etude hématologique, biochimique et parasitaire d’une population de faons de chevreuil (*Capreolus capreolus*). DVM thesis. Ecole nationale vétérinaire de Lyon, France.

Dorchies, B., 2001. Densité d’*Ixodes ricinus*, Linné, 1758, en vie libre : étude dans quatre exploitations de la Sarthe. DVM thesis. Université de Nantes, France.

Dutraive, J., 2019. Study of the correlation between the density of the *Ixodes ricinus* tick and the density of deer in the Toulouse region. DVM thesis. 2019., France.

Ehrmann, S., Liira, J., Gärtner, S., Hansen, K., Brunet, J., Cousins, S.A.O.O., Deconchat, M., Decocq, G., Frenne, P. de, Smedt, P. de, Diekmann, M., Gallet-Moron, E., Kolb, A., Lenoir, J., Lindgren, J., Naaf, T., Paal, T., Valdés, A., Verheyen, K., Wulf, M., Scherer-Lorenzen, M., 2017. Environmental drivers of *Ixodes ricinus* abundance in forest fragments of rural European landscapes. BMC Ecol. 17, 1–14. https://doi.org/10.1186/s12898-017-0141-0

Ehrmann, S., Ruyts, S.C., Scherer-Lorenzen, M., Bauhus, J., Brunet, J., Cousins, S.A.O., Deconchat, M., Decocq, G., De Frenne, P., De Smedt, P., Diekmann, M., Gallet-Moron, E., Gartner, S., Hansen, K., Kolb, A., Lenoir, J., Lindgren, J., Naaf, T., Paal, T., Panning, M., Prinz, M., Valdes, A., Verheyen, K., Wulf, M., Liira, J., 2018. Habitat properties are key drivers of *Borrelia burgdorferi* (s.l.) prevalence in *Ixodes ricinus* populations of deciduous forest fragments. Parasites and Vectors 11, 23. https://doi.org/10.1186/s13071-017-2590-x

Ferquel, E., Garnier, M., Marie, J., Bernède-Bauduin, C., Baranton, G., Pérez-Eid, C., Postic, D., 2006. Prevalence of *Borrelia burgdorferi* sensu lato and Anaplasmataceae members in *Ixodes ricinus* ticks in Alsace, a focus of Lyme borreliosis endemicity in France. Appl. Environ. Microbiol. 72, 3074–3078. https://doi.org/10.1128/AEM.72.4.3074-3078.2006

Ferté, H., Postic, D., Baranton, G., Ulmer, P., Chippaux, C., Léger, N., 1994. First isolation of *Borrelia afzelii* in France (Marne) from *Ixodes ricinus*. Bull. la Société Pathol. Exot. 87, 226–227.

François, J.-B., 2008. Les tiques chez les bovins en France.

Frederic, E., 2005. Babésiose bovine à *Babesia divergens*, étude d’un cas d’émergence en Corrèze. DVM thesis. Université de Nantes, France.

Gasquet, C., 2014. Etude d’un foyer d’anaplasmose bovine dans le département de la Loire. DVM thesis. VetAgro Sup, France.

George, J.C., Chastel, C., 2002. Maladies vectorielles à tiques et modifications de l’écosystème en Lorraine. Bull. la Société Pathol. Exot. 95, 95–99.

Geurden, T., Becskei, C., Six, R.H., Maeder, S., Latrofa, M.S., Otranto, D., Farkas, R., 2018. Detection of tick-borne pathogens in ticks from dogs and cats in different European countries. Ticks Tick. Borne. Dis. 9, 1431–1436. https://doi.org/10.1016/j.ttbdis.2018.06.013

Gilbert, L., Aungier, J., Tomkins, J.L., 2014. Climate of origin affects tick (*Ixodes ricinus*) host-seeking behavior in response to temperature: Implications for resilience to climate change? Ecol. Evol. 4, 1186–1198. https://doi.org/10.1002/ece3.1014

Gilot, B., Bonnefille, M., Degeilh, B., Beaucournu, J.-C.C., Pichot, J., Guiguen, C., 1994. The development of *Ixodes ricinus* (Linné, 1758) in French forests: The roe-deer, *Capreolus capreolus* (L., 1758) used as a biological marker. Parasite 1, 81–85. https://doi.org/10.1051/parasite/1994011081

Gilot, B., Degeilh, B., Pichot, J., Doche, B., Guiguen, C., 1996. Prevalence of *Borrelia burgdorferi* (sensu lato) in *Ixodes ricinus* (L.) populations in France, according to a phytoecological zoning of the territory. Eur. J. Epidemiol. 12, 395–401. https://doi.org/10.1007/bf00145304

Gilot, B., Doche, B., B, Degeilh, B., Guigen, C., Pichot, J., 1995. Bases acarologiques pour l’étude épidémiologique de la borreliose de Lyme: les populations d’*Ixodes ricinus* Linné, 1758 du sud-ouest français. Acarologia 36, 117–132.

Gilot, B., Guiguen, C., Degeilh, B., Doche, B., Pichot, J., Beaucournu, J.C., 1994. Phytoecological mapping of *Ixodes ricinus* as an approach to the distribution of Lyme borreliosis in France, in: Lyme Borreliosis. New York : Plenum, 1984-, France, pp. 105–112. https://doi.org/10.1007/978-1-4615-2415-1_17

Gilot, B., Pautou, G., Moncada, E., 1975. Analysis of vegetation applied to the detection of populations of free-living ticks in south-eastern France: *Ixodes ricinus* (Linne 1758) (Acarina, Ixodoidea) as an example. Acta Trop. 32, 340–347.

Gilot, B., Pautou, G., Moncada, E., Ain, G., 1975. Ecological study of *Ixodes ricinus* (Linné, 1758) (Acarina, Ixodoides) in southeastern France. Acta Trop. 32, 232–258.

Gilot, B., Perez-Eid, C., 1998. Bioecology of ticks causing the most important pathology in France. Médecine Mal. Infect. 28, 325–334. https://doi.org/10.1016/s0399-077x(98)70217-3

Gilot, B., Pichot, J., Doche, B., 1989. The ticks of the Massif Central (France). 1. - The Ixodides (Acari, Ixodoidea) infesting domestic carnivores and ungulates in the eastern border of the Massif. Acarologia 30, 191–207.

Gilot, B., Rogers, P., Lachet, B., 1985. Biological and ecological data on the ticks of lagomorphs (and especially those of the wild rabbit, *Oryctolagus cuniculus* L.) in the French Alps and their surrounding countryside. Acarologia 26, 335–354.

Gilot, B., Marjolet, M., 1982. Contribution to the study of human parasitism by ticks (Ixodidae and Argasidae), particularly in south-eastern France. Med. Mal. Infect. 12, 340–351. https://doi.org/10.1016/S0399-077X(82)80014-0

Gilot, B., Pautou, G., 1982. The evolution of populations of ticks (Ixodidae and Argasidae) in relation to the artificial alteration of the environment in the French Alps. Epidemiologic effects. Acta Trop. 39, 337–354.

Goldstein, V., 2017. Vectorial epidemiology of Lyme borreliosis in France. PhD thesis. Université de Strasbourg, France.

Goldstein, V., Boulanger, N., Schwartz, D., George, J.-C., Ertlen, D., Zilliox, L., Schaeffer, M., Jaulhac, B., 2018. Factors responsible for *Ixodes ricinus* nymph abundance: Are soil features indicators of tick abundance in a French region where Lyme borreliosis is endemic? Ticks Tick. Borne. Dis. 9, 938–944. https://doi.org/10.1016/j.ttbdis.2018.03.013

Gondard, M., Michelet, L., Nisavanh, A., Devillers, E., Delannoy, S., Fach, P., Aspan, A., Ullman, K., Chirico, J., Hoffmann, B., Van Der Wal, F.J., De Koeijer, A., Van Solt-Smits, C., Jahfari, S., Sprong, H., Mansfield, K.L., Fooks, A.R., Klitgaard, K., Bødker, R., Moutailler, S., 2018. Prevalence of tick-borne viruses in *Ixodes ricinus* assessed by high-throughput real-time PCR. Pathog. Dis. 76, fty083 (13 p). https://doi.org/10.1093/femspd/fty083

Grech-Angelini, S., Stachurski, F., Lancelot, R., Boissier, J., Allienne, J.-F., Marco, S., Maestrini, O., Uilenberg, G., 2016. Ticks (Acari: Ixodidae) infesting cattle and some other domestic and wild hosts on the French Mediterranean island of Corsica. Parasites and Vectors 9, 582. https://doi.org/10.1186/s13071-016-1876-8

Grech-Angelini, S., Stachurski, F., Vayssier-Taussat, M., Devillers, E., Casabianca, F., Lancelot, R., Uilenberg, G., Moutailler, S., 2020. Tick-borne pathogens in ticks (Acari: Ixodidae) collected from various domestic and wild hosts in Corsica (France), a Mediterranean island environment. Transbound. Emerg. Dis. 67, 745–757. https://doi.org/10.1111/tbed.13393

Grégoire, A., Faivre, B., Heeb, P., Cezilly, F., 2002. A comparison of infestation patterns by Ixodes ticks in urban and rural populations of the Common Blackbird *Turdus merula*. Ibis (Lond. 1859). 144, 640–645. https://doi.org/https://doi.org/10.1046/j.1474-919X.2002.00102.x

Guétard, M., 2001. Ixodes ricinus : morphologie, biologie, élevage, données bibliographiques. DVM thesis. Université Paul sabatier (Toulouse), France.

[Guiguen, C., Degeilh, B., 2001. Ticks of medical interest: vector role and laboratory diagnosis. Rev. Française des Lab. 2001, 49–57. https://doi.org/https://doi.org/10.1016/S0338-9898(01)80351-6](https://doi.org/https://doi.org/10.1016/S0338-9898(01)80351-6)

Guiguen, C., Belaz, S., Degeilh, B., 2019. Bio-ecology and pathogenic role of ticks in metropolitan France. Rev. Francoph. des Lab. 513, 24–33. https://doi.org/10.1016/s1773-035x(19)30286-2

Halos, L., Chauvin, A., 2009. Hard ticks of ruminants in metropolitan France. Bull. la Société Vétérinaire Prat. Fr. 93, 23–24.

Halos, L., Bord, S., Cotté, V., Gasqui, P., Abrial, D., Barnouin, J., Boulouis, H.-J., Vayssier-Taussat, M., Vourc’h, G., 2010. Ecological factors characterizing the prevalence of bacterial tick-borne pathogens in *Ixodes ricinus* ticks in pastures and woodlands. Appl. Environ. Microbiol. 76, 4413–4420. https://doi.org/10.1128/AEM.00610-10

Halos, L., Jamal, T., Maillard, R., Beugnet, F., Le Menach, A., Boulouis, H.-J., Vayssier-Taussat, M., 2005. Evidence of *Bartonella* sp. in questing adult and nymphal *Ixodes ricinus* ticks from France and co-infection with *Borrelia burgdorferi* sensu lato and *Babesia* sp. Vet. Res. 36, 79–87. https://doi.org/10.1051/vetres:2004052

Halos, L., Jamal, T., Vial, L. V, Maillard, R., Suau, A., Le Menache, A., Bulouis, H.-J., Vayssier-Taussat, M., 2004. Determination of an efficient and reliable method for DNA extraction from ticks. Vet. Res. 35, 709–713. https://doi.org/10.1051/vetres:2004038

Halos, L., Mavris, M., Vourc’h, G., Maillard, R., Barnouin, J., Boulouis, H.J., Vayssier-Taussat, M., 2006. Broad-range PCR-TTGE for the first-line detection of bacterial pathogen DNA in ticks. Vet. Res. 37, 245–253. https://doi.org/10.1051/vetres:2005055

Halos, L., Vourc’h, G., Cotte, V., Gasqui, P., Barnouin, J., Boulous, H.-J., Vayssier-Taussat, M., 2006. Prevalence of *Anaplasma phagocytophilum*, *Rickettsia* sp. and *Borrelia burgdorferi* sensu lato DNA in questing *Ixodes ricinus* ticks from France. Ann. N. Y. Acad. Sci. 1078, 316–319. https://doi.org/10.1196/annals.1374.059

Jacquot, M., 2014. Genomic diversity of pathogenic bacteria in the *Borrelia burgdorferi* species complex: evolution and molecular epidemiology. PhD thesis. Université Blaise Pascal de Clermont-Ferrand, France.

Jacquot, M., Abrial, D., Gasqui, P., Bord, S., Marsot, M., Masseglia, S., Pion, A., Poux, V., Zilliox, L., Chapuis, J., Vourc, G., Bailly, X., 2016. Multiple independent transmission cycles of a tick-borne pathogen within a local host community. Sci. Rep. 6, 1–12. https://doi.org/10.1038/srep31273

Jacquot, M., Gonnet, M., Ferquel, E., Abrial, D., Claude, A., Gasqui, P., Choumet, V., Charras-Garrido, M., Garnier, M., Faure, B., Sertour, N., Dorr, N., De Goër, J., Vourc’H, G., Bailly, X., 2014. Comparative population genomics of the *Borrelia burgdorferi* species complex reveals high degree of genetic isolation among species and underscores benefits and constraints to studying intra-specific epidemiological processes. PLoS One 9. https://doi.org/10.1371/journal.pone.0094384

Jolivet, F., 1996. Activité et répartition d’*Ixodes ricinus* dans vingt élevages de Sarthe. DVM thesis. Université de Nantes, France.

Jouglin, M., Perez, G., Butet, A., Malandrin, L., Bastian, S., 2017. Low prevalence of zoonotic *Babesia* in small mammals and *Ixodes ricinus* in Brittany, France. Vet. Parasitol. 238, 58–60. https://doi.org/10.1016/j.vetpar.2017.03.020

Kautsmann, L., 2018. Estimation bayésienne du taux d’échantillonnage et de la population à l’affût lors de collecte au drap d’*Ixodes ricinus*. DVM thesis. VetAgro Sup, France.

Kempf, F., De Meeus, T., Arnathau, C., Degeilh, B., McCoy, K.D., 2009. Assortative pairing in *Ixodes ricinus* (Acari: Ixodidae), the European vector of Lyme borreliosis. J. Med. Entomol. 46, 471–474. https://doi.org/10.1603/033.046.0309

Kempf, F., De Meeûs, T., Vaumourin, E., Noel, V., Taragel’ová, V., Plantard, O., Heylen, D.J.A., Eraud, C., Chevillon, C., McCoy, K.D., 2011. Host races in *Ixodes ricinus*, the European vector of Lyme borreliosis. Infect. Genet. Evol. 11, 2043–2048. https://doi.org/10.1016/j.meegid.2011.09.016

Kraemer, D., 2018. Variation intra-journalière de l’activité des tiques *Ixodes ricinus* en fonction des données météorologiques. DVM thesis. VetAgro Sup, France.

Lamontellerie, M., 1965. Les tiques du Sud-Ouest de la France. Ann. Parasitol. 40, 87–100.

Le Coeur, C., Robert, A., Pisanu, B., Chapuis, J.-L., 2015. Seasonal variation in infestations by ixodids on Siberian chipmunks: effects of host age, sex, and birth season. Parasitol. Res. 114, 2069–2078. https://doi.org/10.1007/s00436-015-4391-5

Lebert, I., Agoulon, A., Bastian, S., Butet, A., Cargnelutti, B., Cèbe, N., Chastagner, A., Léger, E., Lourtet, B., Masseglia, S., McCoy, K.D., Merlet, J., Noël, V., Perez, G., Picot, D., Pion, A., Poux, V., Rames, J.-L., Rantier, Y., Verheyden, H., Vourc’h, G., Plantard, O., 2020. Distribution of ticks and tick-borne pathogenic agents and associated local environmental conditions, small mammal hosts and livestock in two French agricultural sites: the OSCAR database. Biodivers. Data J. 8, e50123 (1-34).

Leger, E., 2013. Host community structure and the evolution of host specialisation in the tick *Ixodes ricinus*. PhD thesis. Université de Montpellier I, France.

Lejal, E., Lejal, E., Lejal, E., 2022. La dynamique du pathobiome des tiques, l’exemple d’*Ixodes ricinus*.

Lejal, E., Marsot, M., Chalvet-Monfray, K., Cosson, J.-F., Moutailler, S., Vayssier-Taussat, M., Pollet, T., 2019. A three-years assessment of *Ixodes ricinus*-borne pathogens in a French peri-urban forest. Parasites and Vectors 12, 551 (14 p). https://doi.org/10.1186/s13071-019-3799-7

Lejal, E., Moutailler, S., Šimo, L., Vayssier-Taussat, M., Pollet, T., 2019. Tick-borne pathogen detection in midgut and salivary glands of adult *Ixodes ricinus*. Parasites and Vectors 12, 152 (8 p). https://doi.org/10.1186/s13071-019-3418-7

Lepigeon, J., 2007. Etudes épidémiologique de la babesiose bovine à *Babesia divergens* dans 20 élevages laitiers de l’Ouest de la France : prévalence ches les tiques libres (*Ixodes ricinus*) et densités de tiques sur les patures. DVM thesis.

L’Hostis, M., 1999. *Babesia divergens* in France: Descriptive and analytical epidemiology. Parassitologia 41, 59–62.

L’Hostis, M., Seegers, H., 2002. Tick-borne parasitic diseases in cattle: Current knowledge and prospective risk analysis related to the ongoing evolution in French cattle farming systems. Vet. Res. 33, 599–611.

L’Hostis, M., Bureaud, A., Gorenflot, A., 1996. Female *Ixodes ricinus* (Acari, Ixodidae) in cattle of western France: infestation level and seasonality. Vet. Res. 27, 589–597.

L’Hostis, M., Dumon, H., Dorchies, B., Boisdron, F., Gorenflot, A., 1995. Seasonal incidence and ecology of the tick *Ixodes ricinus* (Acari: Ixodidae) on grazing pastures in Western France. Exp. Appl. Acarol. 19, 211–220. https://doi.org/10.1007/bf00130824

L’Hostis, M., Diarra, O., Seegers, H., 1994. Sites of attachment and density assessment of female *Ixodes ricinus* (Acari: Ixodidae) on dairy cows. Exp. Appl. Acarol. 18, 681–689. https://doi.org/10.1007/BF00051535

L’Hostis, M., Dumon, H., Fusade, A., Lazareff, S., Gorenflot, A., 1996. Seasonal incidence of *Ixodes ricinus* ticks (Acari: Ixodidae) on rodents in western France. Exp. Appl. Acarol. 20, 359–368. https://doi.org/10.1007/BF00130548

Marchant, A., Le Coupanec, A., Joly, C., Perthame, E., Sertour, N., Garnier, M., Godard, V., Ferquel, E., Choumet, V., 2017. Infection of *Ixodes ricinus* by *Borrelia burgdorferi* sensu lato in peri-urban forests of France. PLoS One 12, e0183543. https://doi.org/10.1371/journal.pone.0183543

Marié, J.-L., Davoust, B., Socolovschi, C., Mediannikov, O., Roqueplo, C., Beaucournu, J.-C., Raoult, D., Parola, P., 2012. Rickettsiae in arthropods collected from red foxes (*Vulpes vulpes*) in France. Comp. Immunol. Microbiol. Infect. Dis. 35, 59–62. https://doi.org/https://doi.org/10.1016/j.cimid.2011.10.001

Marié, J.-L., Davoust, B., Socolovschi, C., Raoult, D., Parola, P., 2012. Molecular detection of rickettsial agents in ticks and fleas collected from a European hedgehog (*Erinaceus europaeus*) in Marseilles, France. Comp. Immunol. Microbiol. Infect. Dis. 35, 77–79. https://doi.org/https://doi.org/10.1016/j.cimid.2011.11.005

Marsot, M., Henry, P.-Y., Vourc’h, G., Gasqui, P., Ferquel, E., Laignel, J., Grysan, M., Chapuis, J.-L., 2012. Which forest bird species are the main hosts of the tick, *Ixodes ricinus*, the vector of *Borrelia burgdorferi* sensu lato, during the breeding season? Int. J. Parasitol. 42, 781–788. https://doi.org/10.1016/j.ijpara.2012.05.010

Marsot, M., 2011. Modification of a multi-host disease risk through the introduction of a reservoir species: the case of Lyme disease and of the Siberian chipmunk in French suburban forests. PhD thesis. Université Blaise-Pascale, Clermont-Ferrand II, France.

Marsot, M., Chapuis, J.L., Gasqui, P., Dozières, A., Masséglia, S., Pisanu, B., Ferquel, E., Vourc’h, G., 2013. Introduced Siberian chipmunks (*Tamias sibiricus barberi*) contribute more to Lyme borreliosis risk than native reservoir rodents. PLoS One 8, 1–8. https://doi.org/10.1371/journal.pone.0055377

Martinod, S., Brossard, M., Moreau, Y., 1985. Immunity of dogs against *Babesia canis*, its vector tick *Dermacentor reticulatus*, and *Ixodes ricinus* in endemic area. J. Parasitol. 71, 269–273. https://doi.org/10.2307/3282004

Martinod, S., Gilot, B., Girel, J., Lachet, B., Laurent, N., 1984. La babésiose canine à *Babesia canis* dans les Alpes Françaises du Nord et le Jura Méridional: Cartographie épidémiologique. Doc. Cartogr. Ecol. 27, 3–20.

Marumoto, K., Joncour, G., Lamanda, P., Inokuma, H., Brouqui, P., 2007. Detection of *Anaplasma phagocytophilum* and *Ehrlichia* sp. HF strains in *Ixodes ricinus* ticks in Brittany, France. Clin. Microbiol. Infect. 13, 338–341. https://doi.org/10.1111/j.1469-0691.2006.01630.x

Meha, C., 2013. Forest and public health risk: the case of Lyme borreliosis. Application to the peri-urban forest of Sénart. PhD thesis. Université paris-Sorbonne, France.

Mémeteau, S., 1994. Epidémiologie de la babesiose bovine à *Babesia divergens* : étude dans vingt élevages laitiers de Sarthe. DVM thesis. Université de Nantes, France.

Mémeteau, S., Seegers, H., Jolivet, F., L’Hostis, M., 1998. Assessment of the risk of infestation of pastures by *Ixodes ricinus* due to their phyto-ecological characteristics. Vet. Res. 29, 487–496. https://doi.org/10.1016/S0928-4249(98)80008-4

Michelet, L., Delannoy, S., Devillers, E., Gérald Umhang, Aspan, A., Juremalm, M., Chirico, J., van der Wal, F.J., Sprong, H., Boye Pihl, T.P., Klitgaard, K., Bødker, R., Fach, P., Moutailler, S., 2014. High-throughput screening of tick-borne pathogens in Europe. Front. Cell. Infect. Microbiol. 4, 1–13. https://doi.org/10.3389/fcimb.2014.00103

Michelet, L., Joncour, G., Devillers, E., Torina, A., Vayssier-Taussat, M., Bonnet, S.I., Moutailler, S., 2016. Tick species, tick-borne pathogens and symbionts in an insular environment off the coast of Western France. Ticks Tick. Borne. Dis. 7, 1109–1115. https://doi.org/https://doi.org/10.1016/j.ttbdis.2016.08.014

Moutailler, S., Popovici, I., Devillers, E., Vayssier-Taussat, M., Eloit, M., 2016. Diversity of viruses in *Ixodes ricinus*, and characterization of a neurotropic strain of Eyach virus. New Microbes New Infect. 11, 71–81. https://doi.org/10.1016/j.nmni.2016.02.012

Moutailler, S., Valiente Moro, C., Vaumourin, E., Michelet, L., Tran, F.H., Devillers, E., Cosson, J.F., Gasqui, P., Van, V.T., Mavingui, P., Vourc’h, G., Vayssier-Taussat, M., 2016. Co-infection of ticks: the rule rather than the exception. PLoS Negl. Trop. Dis. 10, e0004539 (17 p). https://doi.org/10.1371/journal.pntd.0004539

Nebbak, A., Dahmana, H., Almeras, L., Raoult, D., Boulanger, N., Jaulhac, B., Mediannikov, O., Parola, P., 2019. Co-infection of bacteria and protozoan parasites in *Ixodes ricinus* nymphs collected in the Alsace region, France. Ticks Tick. Borne. Dis. 10, 101241. https://doi.org/https://doi.org/10.1016/j.ttbdis.2019.06.001

Noël, V., Leger, E., Gómez-Díaz, E., Risterucci, A.M., McCoy, K.D., 2012. Isolation and characterization of new polymorphic microsatellite markers for the tick *Ixodes ricinus* (Acari: Ixodidae). Acarologia 52, 123–128. https://doi.org/10.1051/acarologia/20122041

Noureddine, R., Chauvin, A., Plantard, O., 2011. Lack of genetic structure among Eurasian populations of the tick *Ixodes ricinus* contrasts with marked divergence from north-African populations. Int. J. Parasitol. 41, 183–192. https://doi.org/10.1016/j.ijpara.2010.08.010

Panas, E., Leger, N., Kretz, J.L., Dumesnil, C., 1976. Ixodidae of the Champagne-Ardennes region. Preliminary study. Acarologia 18, 51–55.

Parola, P., Beati, L., Cambon, M., Raoult, D., 1998. First isolation of Rickettsia helvetica from *Ixodes ricinus* ticks in France. Eur. J. Clin. Microbiol. Infect. Dis. 17, 95–100. https://doi.org/10.1007/s100960050024

Parola, P., Raoult, D., 2001. Molecular tools in the epidemiology of tick-borne bacterial diseases. Ann. Biol. Clin. (Paris). 59, 177–182.

Parola, P., Beati, L., Cambon, M., Brouqui, P., Raoult, D., 1998. Ehrlichial DNA amplified from *Ixodes ricinus* (Acari: Ixodidae) in France. J. Med. Entomol. 35, 180–183. https://doi.org/10.1093/jmedent/35.2.180

Paul, R.E.L., Cote, M., Le Naour, E., Bonnet, S.I., 2016. Environmental factors influencing tick densities over seven years in a French suburban forest. Parasites and Vectors 9, 1–10. https://doi.org/10.1186/s13071-016-1591-5

Pérez, C., 1987. La faune des tiques dans le foyer alsacien d’encéphalite à tiques. Acarologia 28, 43–47.

Perez, G., 2016. Influence of the landscape on the small mammal communities as hosts of tick-borne infectious agents. PhD thesis. Rennes 1, France.

Perez, G., Bastian, S., Agoulon, A., Bouju, A., Durand, A., Faille, F., Lebert, I., Rantier, Y., Plantard, O., Butet, A., 2016. Effect of landscape features on the relationship between *Ixodes ricinus* ticks and their small mammal hosts. Parasites and Vectors 9, 1–18. https://doi.org/10.1186/s13071-016-1296-9

Perez, G., Bastian, S., Chastagner, A., Agoulon, A., Plantard, O., Vourc’h, G., Butet, A., 2017. Ecological factors influencing small mammal infection by *Anaplasma phagocytophilum* and *Borrelia burgdorferi* s.l. in agricultural and forest landscapes. Environ. Microbiol. 19, 4205–4219. https://doi.org/10.1111/1462-2920.13885

Perez, G., Bastian, S., Chastagner, A., Agoulon, A., Rantier, Y., Vourc’h, G., Plantard, O., Butet, A., 2020. Relationships between landscape structure and the prevalence of two tick-borne infectious agents, *Anaplasma phagocytophilum* and *Borrelia burgdorferi* sensu lato, in small mammal communities. Landsc. Ecol. 35, 435–451. https://doi.org/10.1007/s10980-019-00957-x

Pérez-Eid, C., 1990. Les relations tiques-petits mammifères dans le foyer alsacien d’encéphalite à tiques. Acarologia 31, 131–141.

Pérez-Eid, C., 1989. Dynamique saisonnière des nymphes et adultes d’*Ixodes ricinus* en phase libre sur la végétation, dans le foyer alsacien d’encéphalite à tiques. Acarologia 30, 355–360.

Pérez-Eid, C., Hannoun, C., Rodhain, F., 1992. The Alsatian tick-borne encephalitis focus: presence of the virus among ticks and small mammals. Eur. J. Epidemiol. 8, 178–186. https://doi.org/10.1007/bf00144797

Perrain, C., 2013. Étude préliminaire pour la mise en place d’un observatoire de tiques dans la région Rhône-Alpes. DVM thesis. VetAgro Sup, France.

Pichon, B., Mousson, L., Figureau, C., Rodhain, F., Pérez-Eid, C., 1999. Density of deer in relation to the prevalence of *Borrelia burgdorferi* s.l. in *Ixodes ricinus* nymphs in Rambouillet forest, France. Exp. Appl. Acarol. 23, 267–75. https://doi.org/10.1023/a:1006023115617

Pichon, B., 1997. Eco-epidemiological study of Lyme borreliosis in the Rambouillet forest (Yvelines, France): circulation of *Borrelia* of the *burgdorferi* group in the vector *Ixodes ricinus* and its main mammalian host species. PhD thesis. France.

Pichot, J., Gilot, B., Almire, N., Polette, K., Degeilh, B., 1997. *Ixodes* populations (*Ixodes ricinus* Linné, 1758; *Ixodes hexagonus* leach, 1815) in the city of Lyon (France) and its outskirts: Preliminary results. Parasite 4, 167–171. https://doi.org/10.1051/parasite/1997042167

Pichot, J., Gilot, B., Soulier, V., Rey-Coquais, A., Degeilh, B., Doche, B., 1994. Ecoepidemiological study of Lyme borreliosis in Rhône-Alpes area. Parasite 1, 335–342. https://doi.org/10.1051/parasite/1994014335

Pisanu, B., Marsot, M., Marmet, J., Chapuis, J.-L., Réale, D., Vourc’h, G., 2010. Introduced Siberian chipmunks are more heavily infested by ixodid ticks than are native bank voles in a suburban forest in France. Int. J. Parasitol. 40, 1277–1283. https://doi.org/10.1016/j.ijpara.2010.03.012

Pisanu, B., Chapuis, J.L., Dozières, A., Basset, F., Poux, V., Vourc’h, G., 2014. High prevalence of *Borrelia burgdorferi* s.l. in the European red squirrel *Sciurus vulgaris* in France. Ticks Tick. Borne. Dis. 5, 1–6. https://doi.org/10.1016/j.ttbdis.2013.07.007

Quessada, T., Martial-Convert, F., Arnaud, S., Leudet De La Vallee, H., Gilot, B., Pichot, J., 2003. Prevalence of *Borrelia burgdorferi* species and identification of *Borrelia valaisiana* in questing *Ixodes ricinus* in the Lyon region of France as determined by polymerase chain reaction-restriction fragment length polymorphism. Eur. J. Clin. Microbiol. Infect. Dis. 22, 165–173. https://doi.org/10.1007/s10096-002-0866-2

Randolph, S.E., Green, R.M., Peacey, M.F., Rogers, D.J., 2000. Seasonal synchrony: the key to tick-borne encephalitis foci identified by satellite data. Parasitology 121, 15–23. https://doi.org/10.1017/s0031182099006083

Raoux, O., 2011. Distribution spatio-temporelle de la tique *Ixodes ricinus* sur la végétation d’une exploitation bovine du bocage nantais : partie 2 : Analyse à l’échelle des pâtures et de l’exploitation, analyse de la fiabilité de la méthode de collecte au drapeau. DVM thesis. [S.l.] : [s.n.], 2011., France.

Reis, C., Cote, M., Paul, R.E.L., Bonnet, S., 2011. Questing ticks in suburban forest are infected by at least six tick-borne pathogens. Vector-borne Zoonotic Dis. 11, 907–916. https://doi.org/10.1089/vbz.2010.0103

Richter, D., Matuschka, F.R., 2012. “*Candidatus* Neoehrlichia mikurensis”, *Anaplasma phagocytophilum*, and Lyme disease spirochetes in questing European vector ticks and in feeding ticks removed from people. J. Clin. Microbiol. 50, 943–947. https://doi.org/10.1128/JCM.05802-11

Richter, D., Schlee, D.B., Matuschka, F.R., 2003. Relapsing fever-like spirochetes infecting European vector tick of Lyme disease agent. Emerg. Infect. Dis. 9, 697–701. https://doi.org/10.3201/eid0906.020459

Romeo, C., Pisanu, B., Ferrari, N., Basset, F., Tillon, L., Wauters, L.A., Martinoli, A., Saino, N., Chapuis, J.-L., 2013. Macroparasite community of the Eurasian red squirrel (*Sciurus vulgaris*): poor species richness and diversity. Parasitol. Res. 112, 3527–3536. https://doi.org/10.1007/s00436-013-3535-8

Serra, V., Cafiso, A., Formenti, N., Verheyden, H., Plantard, O., Bazzocchi, C., Sassera, D., 2018. Molecular and serological evidence of the presence of *Midichloria mitochondrii* in roe deer (*Capreolus capreolus*) in France. J. Wildl. Dis. 54, 597–600. https://doi.org/10.7589/2017-09-241

Stachurski, F., Vial, L., 2018. Installation de la tique Hyalomma marginatum, vectrice du virus de la fièvre hémorragique de Crimée-Congo, en France continentale. Bull. Epidémiologique Santé Anim. - Aliment. 84, 1–5. https://doi.org/10.1080/01647950808683704

Tomkins, J.L., Aungier, J., Hazel, W., Gilbert, L., 2014. Towards an evolutionary understanding of questing behaviour in the tick *Ixodes ricinus*. PLoS One 9, e110028. https://doi.org/10.1371/journal.pone.0110028

Ulmer, P., Ferté, H., Mercier, A., Richard, S., 1999. Lyme Borreliosis: epidemiological survey carried out in a kennel in north-eastern France. Rev. Française des Lab. 1999, 49–56. https://doi.org/10.1016/S0338-9898(99)80375-8

Vah, B., 2011. Distribution spatio-temporelle de la tique *Ixodes ricinus* sur la végétation d’une exploitation bovine du bocage nantais. Partie 1 : analyse à l’échelle des transects. DVM thesis. [S.l.] : [s.n.], 2011., France.

Vassallo, M., Paul, R.E.L., Pérez-Eid, C., 2000. Temporal distribution of the annual nymphal stock of *Ixodes ricinus* ticks. Exp. Appl. Acarol. 24, 941–949. https://doi.org/10.1023/a:1010669003887

Vassallo, M., Pérez-Eid, C., 2002. Comparative behavior of different life-cycle stages of *Ixodes ricinus* (Acari: Ixodidae) to human-produced stimuli. J. Med. Entomol. 39, 234–236. https://doi.org/10.1603/0022-2585-39.1.234

Vassallo, M., Pichon, B., Cabaret, J., Figureau, C.U., Pérez-Eid, C., 2000. Methodology for sampling questing nymphs of *Ixodes ricinus* (Acari: Ixodidae), the principal vector of Lyme disease in Europe. J. Med. Entomol. 37, 335–339. https://doi.org/https://doi.org/10.1093/jmedent/37.3.335

Vassallo, M.-P., 2000. Lyme borreliosis: study of the densities of the main vector, the group of species Ixodes ricinus and its relationships to the environment. Assessment of areas and periods at risk for humans of contracting the disease. PhD thesis. Université de Paris 6, France.

Vayssier-Taussat, M., Moutailler, S., Michelet, L., Devillers, E., Bonnet, S., Eloit, M., Cheval, J., He, C., Hébert, C., Eloit, M., 2013. Next generation sequencing uncovers unexpected bacterial pathogens in ticks in Western Europe. PLoS One 8, e81439 (7 p). https://doi.org/10.1371/journal.pone.0081439

Vénisse, M., 2007. Epidemiological study of bovine babesiosis with *Babesia divergens* in 20 dairy farms in western France: prevalence in fixed ticks (*Ixodes ricinus*), cattle infestation and serological monitoring. DVM thesis. [S.l.] : [s.n.], 2007., France.

Vourc’h, G., Abrial, D., Bord, S., Jacquot, M., Masseglia, S., Poux, V., Pisanu, B., Bailly, X., Chapuis, J.-L., 2016. Mapping human risk of infection with *Borrelia burgdorferi* sensu lato, the agent of Lyme borreliosis, in a periurban forest in France. Ticks Tick. Borne. Dis. 7, 644–652. https://doi.org/10.1016/j.ttbdis.2016.02.008

Vourc’h, G., Marmet, J., Chassagne, M., Bord, S., Chapuis, J.-L., 2007. *Borrelia burgdorferi* sensu lato in Siberian chipmunks (*Tamias sibiricus*) introduced in suburban forests in France. Vector-borne Zoonotic Dis. 7, 637–641. https://doi.org/10.1089/vbz.2007.0111

Zhioua, E., Postic, D., Rodhain, F., Perez-Eid, C., 1996. Infection of *Ixodes ricinus* (Acari: Ixodidae) by *Borrelia burgdorferi* in Ile de France. J. Med. Entomol. 33, 694–697. https://doi.org/10.1093/jmedent/33.4.694

This is a supplementary material for “Perez, G., Bournez, L., Boulanger, N., Fite, J., Livoreil, B., McCoy, K. D., Quillery, E., René-Martellet, M., and Bonnet, S. I. The distribution, phenology, host range and pathogen prevalence of *Ixodes ricinus* in France: a systematic map and a narrative review”, a preprint recommended in PCI Infections <https://doi.org/10.24072/pci.infections.100076>.
