## Supplementary material for "The distribution, phenology, host range and pathogen prevalence of *Ixodes ricinus* in France: a systematic map and narrative review": SI 6

**Supplementary information 6: Temporal trend of publications**

Temporal trend of identified publications on *Ixodes ricinus* ecology in European France by year for each publication type: PhD theses (n = 15), DPharm thesis (n = 1), book section (n = 1), DVM thesis (n = 21), and research and review articles (n = 153).

Number of publications

|  | Reasearch article & reviews | Book section | DVM theses | DPharm thesis | PhD theses |
| --- | --- | --- | --- | --- | --- |
| 1965 | 1 |  |  |  |  |
| 1975 | 3 |  |  |  |  |
| 1976 | 1 |  |  |  |  |
| 1978 | 1 |  |  |  |  |
| 1979 | 1 |  |  |  |  |
| 1982 | 2 |  |  |  |  |
| 1983 |  |  |  |  | 1 |
| 1984 | 2 |  |  |  |  |
| 1985 | 3 |  |  |  |  |
| 1986 | 1 |  |  |  |  |
| 1987 | 1 |  |  |  |  |
| 1989 | 4 |  |  |  |  |
| 1990 | 3 |  |  |  |  |
| 1991 | 2 |  | 1 |  |  |
| 1992 | 3 |  | 1 |  |  |
| 1993 | 3 |  |  |  |  |
| 1994 | 6 | 1 | 1 |  |  |
| 1995 | 3 |  |  |  |  |
| 1996 | 6 |  | 1 |  | 1 |
| 1997 | 1 |  |  |  | 1 |
| 1998 | 5 |  |  |  |  |
| 1999 | 4 |  |  |  |  |
| 2000 | 3 |  |  |  | 1 |
| 2001 | 2 |  | 2 |  |  |
| 2002 | 4 |  |  |  |  |
| 2003 | 2 |  |  |  |  |
| 2004 | 1 |  |  |  |  |
| 2005 | 1 |  | 1 |  |  |
| 2006 | 3 |  |  |  |  |
| 2007 | 5 |  | 3 |  | 1 |
| 2008 | 1 |  |  | 1 |  |
| 2009 | 2 |  | 2 |  |  |
| 2010 | 5 |  |  |  |  |
| 2011 | 4 |  | 2 |  | 1 |
| 2012 | 7 |  | 1 |  |  |
| 2013 | 5 |  | 1 |  | 2 |
| 2014 | 8 |  | 1 |  | 3 |
| 2015 | 2 |  |  |  |  |
| 2016 | 8 |  |  |  | 1 |
| 2017 | 9 |  |  |  | 2 |
| 2018 | 9 |  | 2 |  |  |
| 2019 | 8 |  | 2 |  |  |
| 2020 | 4 |  |  |  | 1 |

This is a supplementary material for “Perez, G., Bournez, L., Boulanger, N., Fite, J., Livoreil, B., McCoy, K. D., Quillery, E., René-Martellet, M., and Bonnet, S. I. The distribution, phenology, host range and pathogen prevalence of *Ixodes ricinus* in France: a systematic map and a narrative review”, a preprint recommended in PCI Infections <https://doi.org/10.24072/pci.infections.100076>.
