## Supplementary material for "The distribution, phenology, host range and pathogen prevalence of *Ixodes ricinus* in France: a systematic map and narrative review": SI 7

### Supplementary information 7:
### Map showing the 96 French European departments

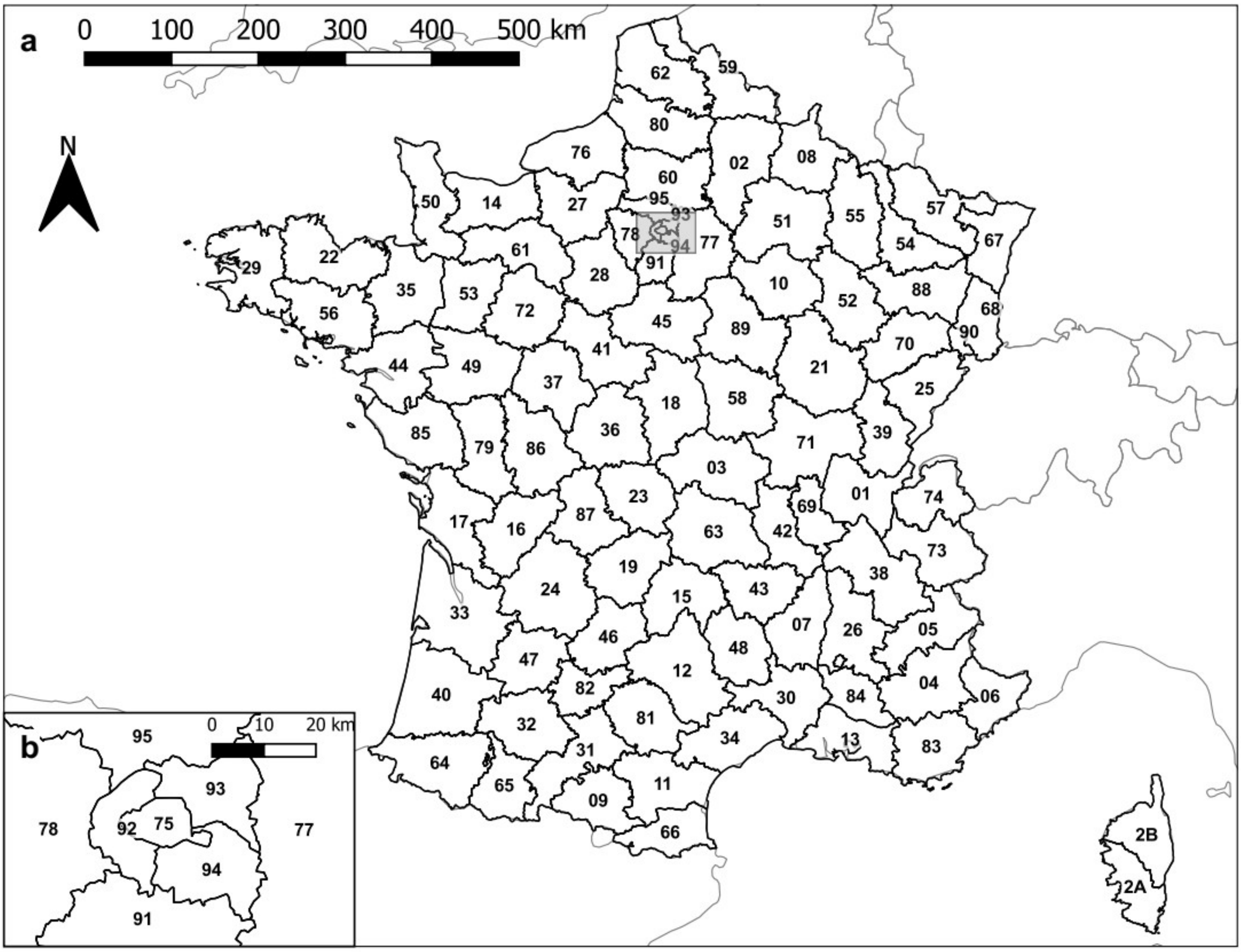


#### Departments

- 01: Ain
- 02: Aisne
- 03: Allier
- 04: Apes-de-Haute-Provence
- 05: Hautes-Alpes
- 06: Alpes-Maritimes
- 07: Ardèche
- 08: Ardennes
- 09: Ariège
- 10: Aube
- 11: Aude
- 12: Aveyron
- 13: Bouches-du-Rhônes
- 14: Calvados
- 15: Cantal
- 16: Charente
- 17: Charente-Maritime
- 18: Cher
- 19: Corrèze
- 2A: Corse-du-Sud
- 2B: Haute-Corse
- 21: Côte-d'Or
- 22: Côtes-d'Armor
- 23: Creuse
- 24: Dordogne
- 25: Doubs
- 26: Drôme
- 27: Eure
- 28: Eure-et-Loir
- 29: Finistère
- 30: Gard
- 31: Haute-Garonne
- 32: Gers
- 33: Gironde
- 34: Hérault
- 35: Ille-et-Vilaine
- 36: Indre
- 37: Indre-et-Loire
- 38: Isère
- 39: Jura
- 40: Landes
- 41: Loir-et-Cher
- 42: Loire
- 43: Haute-Loire
- 44: Loire-Atlantique
- 45: Loiret
- 46: Lot
- 47: Lot-et-Garonne
- 48: Lozère
- 49: Maine-et-Loire
- 50: Manche
- 51: Marne
- 52: Haute-Marne
- 53: Mayenne
- 54: Meurthe-et-Moselle
- 55: Meuse
- 56: Morbihan
- 57: Moselle
- 58: Nièvre
- 59: Nord
- 60: Oise
- 61: Orne
- 62: Pas-de-Calais
- 63: Puy-de-Dôme
- 64: Purénées-Atlantiques
- 65: Hautes-Pyrénées
- 66: Pyrénées-Orientales
- 67: Bas-Rhin
- 68: Haut-Rhin
- 69: Rhône
- 70: Haute-Saône
- 71: Saône-et-Loire
- 72: Sarthe
- 73: Savoie
- 74: Haute-Savoie
- 75: Paris
- 76: Seine-Maritime
- 77: Seine-et-Marne
- 78: Yvelines
- 79: Deux-Sèvres
- 80: Somme
- 81: Tarn
- 82: Tarn-et-Garonne
- 83: Var
- 84: Vaucluse
- 85: Vendée
- 86: Vienne
- 87: Haute-Vienne
- 88: Vosges
- 89: Yonne
- 90: Territoire-de-Belfort
- 91: Essonne
- 92: Hauts-de-Seine
- 93: Seine-Saint-Denis
- 94: Val-de-Marne
- 95: Val-d'Oise

This is a supplementary material for “Perez, G., Bournez, L., Boulanger, N., Fite, J., Livoreil, B., McCoy, K. D., Quillery, E., René-Martellet, M., and Bonnet, S. I. The distribution, phenology, host range and pathogen prevalence of *Ixodes ricinus* in France: a systematic map and a narrative review”, a preprint recommended in PCI Infections https://doi.org/10.24072/pci.infections.100076.
