## Supplementary material for "The distribution, phenology, host range and pathogen prevalence of *Ixodes ricinus* in France: a systematic map and narrative review": SI 8

**Supplementary information 8: vertebrate hosts of *Ixodes ricinus* in European France**

Vertebrate species studied as potential hosts for *Ixodes ricinus* in European France and infestation level classes by tick stage.

| **Scientific name** | **Common name** | ***I*. *ricinus* stages (number of inspected hosts)** | **References (original or related reference)** |
| --- | --- | --- | --- |
| **Aves** | **Birds** |  |  |
| **Passeriformes** | **Passerines** |  |  |
| *Anthus pratensis* | Meadow pipit | Not detailed (?) | Kempf et al. 2011 |
| *Anthus trivialis* | Tree pipit | Not detailed (?) | Kempf et al. 2011 |
| *Certhia brachydactyla* | Short-toed treecreeper | 0 (7) | Marsot et al. 2012 |
| *Chloris chloris* | European greenfinch | F (?) | Grech-Angelini et al. 2016 |
| *Cyanistes caeruleus* | Eurasian blue tit | L, N, L<N (13) | Marsot et al. 2012 |
| *Erithacus rubecula* | Eurasian robin | Not detailed (?) | Kempf et al. 2011 |
|  |  | L++, N (99) | Marsot et al. 2012 |
| *Fringilla coelebs* | Common chaffinch | L, N (14) | Marsot et al. 2012 |
| *Garrulus glandarius* | Eurasian jay | L, N (2) | Marsot et al. 2012 |
| *Luscinia megarhynchos* | Common nightingale | Not detailed (?) | Kempf et al. 2011 |
| *Motacilla flava* | Western yellow wagtail | Not detailed (?) | Grech-Angelini et al. 2016 |
| *Parus major* | Great tit | Not detailed (?) | Kempf et al. 2011 |
|  |  | L+, N (98) | Marsot et al. 2012 |
| *Phoenicurus phoenicurus* | Common redstart | L, N (3) | Marsot et al. 2012 |
| *Phylloscopus collybita* | Common chiffchaff | Not detailed (?) | Kempf et al. 2011 |
|  |  | L, N (28) | Marsot et al. 2012 |
| *Poecile palustris* | Marsh tit | 0 (14) | Marsot et al. 2012 |
| *Prunella modularis* | Dunnock | L+, N+, L < N (8) | Marsot et al. 2012 |
| *Sitta europaea* | Eurasian nuthatch | N (7) | Marsot et al. 2012 |
| *Sturnus vulgaris* | Common starling | L, N, L < N (4) | Marsot et al. 2012 |
| *Sylvia atricapilla* | Eurasian blackcap | Not detailed (?) | Kempf et al. 2011 |
|  |  | L, N (41) | Marsot et al. 2012 |
| *Sylvia borin* | Garden warbler | 0 (3) | Marsot et al. 2012 |
| *Sylvia communis* | Common whitethroat | Not detailed (?) | Kempf et al. 2011 |
| *Troglodytes troglodytes* | Eurasian wren | Not detailed (?) | Kempf et al. 2011 |
|  |  | L, N (32) | Marsot et al. 2012 |
| *Turdus merula* | Common blackbird | L&N++ (106); L&N (224)^c^ | Grégoire et al. 2002 |
|  |  | Not detailed (?) | Kempf et al. 2011 |
|  |  | L++, N++, L<N (55) | Marsot et al. 2012 |
|  |  | Not detailed (?) | Grech-Angelini et al. 2016 |
| *Turdus philomelos* | Song thrush | Not detailed (?) | Kempf et al. 2011 |
|  |  | L++, N+ (13) | Marsot et al. 2012 |
| *Turdus viscivorus* | Mistle thrush | L, N (4) | Marsot et al. 2012 |
| **Piciformes** | **Piciformes** |  |  |
| *Dendrocopos major* | Great spotted woodpecker | 0 (11) | Marsot et al. 2012 |
| *Picus viridis* | European green woodpecker | N (2) | Marsot et al. 2012 |
| **Mammalia** | **Mammals** |  |  |
| **Carnivora** | **Carnivores** |  |  |
| *Canis lupus familiaris* | Dog | F, M (?) | Panas et al. 1976 |
|  |  | A++ (249); A (190)^c^ | Martinod et al. 1985 |
|  |  | F, M (?) | Gilot et al. 1989 |
|  |  | A (?) | Ulmer et al. 1999 |
|  |  | 0 (31) | Grech-Angelini et al. 2016 |
|  |  | F‘+’, N<F (?) | Geurden et al. 2018 |
| *Vulpes vulpes* | Red fox | L, N, F, M, N<F (203)^d^ | Aubert et al. 1976 |
|  |  | A+ (3) | Doby et al. 1994 |
|  |  | 0 (4) | Marié et al. 2012 |
| *Felis catus* | Domestic cat | F, M (?) | Panas et al. 1976 |
|  |  | L, N, F (?) | Pichot et al. 1997 |
|  |  | 1, not detailed (4) | Grech-Angelini et al. 2016 |
|  |  | F‘+’ N<F (?) | Geurden et al. 2018 |
| *Mustela furo* | Ferret | 0 (?) | Panas et al. 1976 |
| **Cetartiodactyla** | **“Even-toed ungulates”** |  |  |
| *Bos taurus* | Cattle | F, M (?) | Gilot et al. 1989 |
|  |  | F++ (110)^a^ | L’Hostis et al. 1994 |
|  |  | F (173-207)^b^ | L’Hostis et al. 1996a |
|  |  | Not detailed (418) | Grech-Angelini et al. 2016 |
|  |  | 0 (42) | Cicculli et al. 2019b |
| *Capra hircus* | Domestic goat | 0 (258) | Grech-Angelini et al. 2016 |
| *Ovis aries* | Sheep | F (?) | Gilot et al. 1989  (citing Nevers 1987) |
|  |  | 0 (51) | Grech-Angelini et al. 2016 |
|  |  | 0 (60) | Cicculli et al. 2019b |
| *O*. *aries musimon* | European mouflon | 0 (24) | Grech-Angelini et al. 2016 |
| *Rupicapra pyrenaica* | Pyrenean chamois | N+, F++, M, N<F (7) | Davoust et al. 2012 |
| *Capreolus capreolus* | Roe deer | F (?) | Panas et al. 1976 |
|  |  | L, N+, F+, M+ (39) | Doby et al. 1994 |
|  |  | L+, N+ (105); L, N (32)^c^ | Gilot et al. 1994 |
|  |  | N, A, N<A (?) | Kempf et al. 2011 |
|  |  | Not detailed (7) | Serra et al. 2018 |
|  |  | N (?) | Cafiso et al. 2019 |
| *Cervus elaphus* | Red deer | F (?) | Panas et al. 1976 |
|  |  | L, N, A++ (12) | Doby et al. 1994 |
| *C*. *elaphus corsicanus* | Corsican red deer | Not detailed (1) | Grech-Angelini et al. 2016 |
| *Sus scrofa* | Eurasian wild boar | A (49) | Doby et al. 1994 |
|  |  | Not detailed (56) | Grech-Angelini et al. 2016 |
| **Eulipotyphla** | **“Insectivores”** |  |  |
| *Erinaceus europaeus* | West European hedgehog | 0 (1) | Marié et al. 2012 |
|  |  | 0 (1) | Grech-Angelini et al. 2016 |
| *Crocidura russula* | Greater white-toothed shrew | 0 (5) | L’Hostis et al. 1996b |
|  |  | L (12) | Boyard et al. 2008 |
| *Crocidura suaveolens* | Lesser white-toothed shrew | 0 (9) | L’Hostis et al. 1996b |
| *Crocidura* sp. | White-toothed shrews | 0 (18) | L’Hostis et al. 1996b |
| *Sorex coronatus* | Crowned shrew | 0 (3) | Perez et al. 2017  (Perez et al. 2016;  Lebert et al. 2020) |
| *Sorex* sp. | Long-tailed shrews | L (17) | Boyard et al. 2008 |
| **Lagomorpha** | **Lagomorphs** |  |  |
| *Oryctolagus cuniculus* | European rabbit | M (1) | Chastel et al. 1984 |
|  |  | N, F, M (700) | Gilot et al. 1985 |
| **Perissodactyla** | **“Odd-toed ungulates”** |  |  |
| *Equus caballus* | Horse | F, M (?) | Gilot et al. 1989  (citing Monteil 1987) |
|  |  | 0 (29) | Grech-Angelini et al. 2016 |
| **Rodentia** | **Rodents** |  |  |
| *Apodemus flavicollis* | Yellow-necked mouse | L+, N (5) | Anderson et al. 1986 |
|  |  | L++ (8)^e^ | Richter et al. 2004 |
|  |  | L+ (18) | Boyard et al. 2008 |
|  |  | L, N (227) | Bournez et al. 2020a |
| *A*. *sylvaticus* | Wood mouse | L++, N (60) | Anderson et al. 1986 |
|  |  | L+, N (175) | Doby et al. 1992b |
|  |  | L+, [N]^f^ (572) | L’Hostis et al. 1996b |
|  |  | L+ (11)^e^ | Richter et al. 2004 |
|  |  | L++ (22) | Vourc’h et al. 2007 |
|  |  | L++, N (42) | Boyard et al. 2008 |
|  |  | L++ (34) | Marsot et al. 2013  (Pisanu et al. 2010) |
|  |  | L, N (453) | Perez et al. 2017  (Perez et al. 2016;  Lebert et al. 2020) |
| *Apodemus* sp. | “Wild mice” | L++, N (1174) | Pérez-Eid 1990  (Pérez 1987) |
| *Eliomys quericinus* | Garden dormouse | L++ (35)^e^ | Richter et al. 2004 |
| *Microtus agrestis* | Short-tailed field-vole | L, [N] ^f^ (62) | L’Hostis et al. 1996b |
|  |  | L (4) | Perez et al. 2017  (Perez et al. 2016;  Lebert et al. 2020) |
| *M*. *arvalis* | Common vole | L, [N] ^f^ (19) | L’Hostis et al. 1996b |
|  |  | L+, N (26) | Boyard et al. 2008 |
| *M*. *subterraneus* | European pine vole | 0 (2) | Perez et al. 2017  (Perez et al. 2016;  Lebert et al. 2020) |
| *Muscardinus avellanarius* | Hazel dormouse | L++ (4)^e^ | Richter et al. 2004 |
| *Myodes glareolus* | Bank vole | L+ (8) | Anderson et al. 1986 |
|  |  | L+, N (1,529) | Pérez-Eid 1990  (Pérez 1987) |
|  |  | L, N (136) | Doby et al. 1992b |
|  |  | L, [N] ^f^ (113) | L’Hostis et al. 1996b |
|  |  | L ++ (2)^e^ | Richter et al. 2004 |
|  |  | L+, N (22) | Vourc’h et al. 2007 |
|  |  | L, N (79) | Boyard et al. 2008 |
|  |  | L+ (547)^g^ | Marsot et al. 2013  (Pisanu et al. 2010) |
|  |  | L (147) | Perez et al. 2017  (Perez et al. 2016;  Lebert et al. 2020) |
|  |  | L, N (333) | Bournez et al. 2020a |
| *Rattus rattus* | Black rat | F+, N<F (9) | Cicculli et al. 2019b |
| *Sciurus vulgaris* | Eurasian red squirrel | Not detailed (311) | Romeo et al. 2013 |
|  |  | L+, N+ (237) | Pisanu et al. 2014 |
| *Tamias sibiricus* (*barberi*) | Siberian chipmunk | L++, N++ (29) | Vourc’h et al. 2007 |
|  |  | L++, N++ (800)^g^ | Marsot et al. 2013  (Pisanu et al. 2010;  Boyer et al. 2010  Le Coeur et al. 2011) |

Notes: 0: no tick; L: larvae; N: nymphs; F: adult females adults; M: adult males ad; A: adults of unspecified sex; L&N: larvae and/or nymphs; Not detailed: tick stages were not detailed; +: 1-5 ticks per host on average; ++: more than five ticks per host on average; L<N: more nymphs than larvae; N<F: more adult females than nymphs; ‘+’: average based on infested hosts only and thus overestimated (burden intensity and not mean burden); ?: unknown number of hosts.

^a^ Larvae and nymphs not studies.

^b^ Study on a variable host number over two years, only host axilla were inspected.

^c^ Two study areas.

^d^ Only host feet were inspected.

^e^ Nymphs and adults not mentioned.

^f^ Possible presence of nymphs, but results insufficiently detailed.

^g^ Extrapolated data from inspection of the head, adults not mentioned, but rare, other tick species reported (*I*. *acuminatus*) (Vourc’h et al. 2007).

This is a supplementary material for “Perez, G., Bournez, L., Boulanger, N., Fite, J., Livoreil, B., McCoy, K. D., Quillery, E., René-Martellet, M., and Bonnet, S. I. The distribution, phenology, host range and pathogen prevalence of *Ixodes ricinus* in France: a systematic map and a narrative review”, a preprint recommended in PCI Infections <https://doi.org/10.24072/pci.infections.100076>.
