## Supplementary material for "The distribution, phenology, host range and pathogen prevalence of *Ixodes ricinus* in France: a systematic map and narrative review": SI 9

**Supplementary information 9: *Anaplasma* prevalence in *Ixodes ricinus* ticks**

Prevalence of *Anaplasma* spp. in questing *Ixodes ricinus* ticks sampled in European France by tick stage.

| Region, department | Reference | Sampling | Method | *Anaplasma* species^a^ | Positive/tested |
| --- | --- | --- | --- | --- | --- |
| Alsace | Richter & Matuschka 2012 | Questing | PCRc  + seq | *A*. *phagocytophilum* | A: 0/60 |
| Alsace, Bas-Rhin | Michelet et al. 2014 | Questing | PCRqm | *A. phagocytophilum* | **N (g. n = 25): 12/47** |
|  |  |  |  | *A. centrale*, *A. marginale*, *A. ovis* and *A. platys* | N (g. n = 25): 0/47 |
|  | Vayssier-Taussat et al. 2013 | Questing | Hseq | *Anaplasma* spp. (*A. phagocytophilum*) | **N (g. n = 1,450): 1/1** |
| Alsace, Haut-Rhin | Ferquel et al. 2006 | Questing | PCRc  + seq | Anaplasmataceae  (*A. phagocytophilum*) | **N: 4/1,065**  **A: 2/171** |
|  | Michelet et al. 2014 | Questing | PCRqm | *A. phagocytophilum* | **N (g. n = 25): 8/47** |
|  |  |  |  | *A. centrale*, *A. marginale*, *A. ovis* and *A. platys* | N (g. n = 25): 0/47 |
|  | Nebbak et al. 2019 | Questing | PCRq  + seq | Anaplasmataceae  (*Wolbachia* spp.: 4/27) | **N: 27/50** |
| Auvergne, Puy-de-Dôme | Beytout et al. 2007 | Questing | PCRn/c + seq | *A*. *phagocytophilum* | **N (525), A (526): 2/1,051** |
|  | Halos et al. 2006a (Halos et al.2004) | Questing | PCRc  + PCRT  + seq | *A. phagocytophilum* | **N (g. n = 10): 11/55** |
|  | Halos et al. 2010  Halos et al. 2006b | Questing | PCRc  + seq | *A*. *phagocytophilum* | **N (g. n = 5): 6/56**  **N (g. n = 10): 46/220**  **N (g. n = 50): 12/20**  **F: 7/102**  **M: 8/123** |
|  | Parola et al. 1998b | Questing | PCRc  + seq | *A*. *phagocytophilum* | **A: 1/80** |
| Burgundy, Côte-d’Or | Bonnet et al. 2013 | Questing | PCRq | *A*. *marginale* and *A. phagocytophilum* | F: 0/1 |
| Burgundy, Saône-et-Loire | Bonnet et al. 2013 | Questing | PCRq | *A. marginale* and *A. phagocytophilum* | F: 0/3 |
| Burgundy, Yonne | Bonnet et al. 2013 | Questing | PCRq | *A. marginale* and *A. phagocytophilum* | F: 0/3  M: 0/1 |
| Brittany | Marumoto et al. 2007 | Questing | PCRc  + seq | *A. phagocytophilum* | **? (g. n = ?): 18/113 (1,706 ticks)** |
| Brittany, Côte-d’Armor | Bonnet et al. 2017 | Questing | PCRc  + seq | *Anaplasma/Midichloria/ Wolbachia* spp. | **N (g. n = 5): 45/86**  **F: 10/60**  **M: 4/85** |
|  |  |  |  | - *A. phagocytophilum* | **N (g. n = 5): 1/86**  F: 0/60  M: 0/85 |
| Brittany, Ille-et-Vilaine | Lebert et al. 2020 | Questing | PCRq  + seq | *A. phagocytophilum* | **N: 35/2,627** |
| Champagne-Ardenne, Ardennes | Moutailler et al. 2016b | Questing | PCRqm | *A*. *phagocytophilum* | **F: 6/267** |
| Corsica | Grech-Angelini et al. 2020a | Attached (cattle, red deer, wild boar, cat) | PCRqm | *A*. *marginale* | **A: 2/115** |
|  |  |  |  | *A*. *phagocytophilum* | **A: 72/115** |
|  |  |  |  | *A. bovis*, *A*. *centrale*, *A*. *ovis* and *A*. *platys* | A: 0/115 |
| Corsica, Haute-Corse | Cicculli et al. 2019b | Attached (black rats) | PCRq/c | Anaplasmaceae | F: 0/13 |
| Île-de-France, Essonne | Lejal et al. 2019a | Questing | PCRqm | *A*. *phagocytophilum* | **F: 7 (2-2-3)^b^/30**  **M: 8 (1-1-6)^b^/30** |
|  |  |  |  | *A. bovis*, *A. centrale*, *A. marginale*, *A*. *ovis* and *A*. *platys* | F: 0^b^/30  M: 0^b^/30 |
|  | Lejal et al. 2019b | Questing | PCRqm | *A*. *phagocytophilum* | **N: 53/998** |
|  |  |  |  | *A. bovis*, *A. centrale*, *A. marginale*, *A*. *ovis* and *A*. *platys* | N: 0/998 |
|  |  |  |  | *Anaplasma* sp. | **N: 1/998** |
|  | Paul et al. 2016 | Questing | PCRc  + seq | *Anaplasma* spp.: | **F: 28/259** |
|  |  |  |  | - *A*. *phagocytophilum* | **F: 17/259** |
|  | Reis et al. 2011 | Questing | PCRc  + seq | *Anaplasma* spp. | **N (g. n = 10): 33/36**  **F: 63/69**  **M: 36/122** |
| Lorraine, Meuse | Beytout et al. 2007 | Questing | PCRn  + seq | *A*. *phagocytophilum* | **N (434), A (164): 7/598** |
| Midi-Pyrénées, Haute-Garonne | Lebert et al. 2020  (Chastagner et al. 2017) | Questing | PCRq  + seq | *A. phagocytophilum* | **N: 57/1,891** |
| Midi-Pyrénées, Hautes-Pyrénées | Akl et al. 2019 | Questing | PCRc | Anaplasmataceae | **N: 53/398**  **F: 50/77**  **M: 4/66** |
|  |  |  | PCRn | - *A*. *phagocytophilum* | **N: 7/398 + 6/133^d^**  **F: 1/77 + 1/9^d^**  **M: 1/66 + 0/13^d^** |
| Pays-de-la-Loire, Loire-Atlantique | Cotté et al. 2010 | Questing | PCRc  + seq | *A*. *phagocytophilum* | N: 0/267  **F: 1/203**  **M: 1/102** |
| Poitou-Charentes, Deux-Sèvres | Bonnet et al. 2013 | Questing | PCRq | *A*. *marginale* | F: 0/18  M: 0/12 |
|  |  |  | PCRq | *A*. *phagocytophilum* | **F: 4/18**  **M: 5/12** |
| Rhône-Alpes, Isère | Bonnet et al. 2013 | Questing | PCRq | *A*. *marginale* and  *A. phagocytophilum* | F: 0/4  M: 0/2 |

Notes: “Region, department”: French region and department of sampling; “Reference”: source of the data; “Sampling”: sampling of the ticks: Attached: tick sampled on a host with host species in brackets (shaded), Questing: tick sampled while questing for a host; “Method”: detection method used: PCR: Polymerisation Chain Reaction, PCRc: classical PCR followed by gel electrophoresis, PCRn: nested PCR, PCRq: quantitative PCR, PCRqm: microfluidic quantitative PCR, PCRT: Temporal Temperature Gradient gel Electrophoresis PCR, seq: amplicon sequencing, Hseq: high-throughput sequencing; “*Anaplasma* species”: *Anaplasma* species searched for; “Positive/tested”: number of positive ticks or pool of ticks on the number of tested ones: L: larvae, N: nymphs, F: adult females, M: adult males, A: adults of unspecified sex, ?: tick stage not specified, “g. n = x”: analyses of pooled DNA of several (x) ticks, /?: unknown number of tested ticks or group of ticks. Positive results are in bold.

^a^ Results of sequence analyses when different from expected by PCR are in brackets.

^b^ Identified as *Ehrlichia phagocytophila* and *E*. *equii*, former denominations.

^c^ Results respectively for salivary glands alone, gut alone, and both.

^d^ PCR performed to search directly *A*. *phagocytophilum*.

This is a supplementary material for “Perez, G., Bournez, L., Boulanger, N., Fite, J., Livoreil, B., McCoy, K. D., Quillery, E., René-Martellet, M., and Bonnet, S. I. The distribution, phenology, host range and pathogen prevalence of *Ixodes ricinus* in France: a systematic map and a narrative review”, a preprint recommended in PCI Infections <https://doi.org/10.24072/pci.infections.100076>.
