## Supplementary material for "The distribution, phenology, host range and pathogen prevalence of *Ixodes ricinus* in France: a systematic map and narrative review": SI 10

**Supplementary information 10: *Bartonella* prevalence in *Ixodes ricinus* ticks**

Prevalence of *Bartonella* spp. in questing *Ixodes ricinus* ticks sampled in European France by tick stage.

| Region, department | Reference | Sampling | Method | *Bartonella* spp. ^a^ | Positive/tested |
| --- | --- | --- | --- | --- | --- |
| Alsace, Bas-Rhin | Michelet et al. 2014 | Questing | PCRqm | *B.* *henselae* | **N (g. n = 25): 1/47** |
|  |  |  |  | *B.* *quintana* | N (g. n = 25): 0/47 |
|  | Dietrich et al. 2010 |  | PCRn  + seq | *B. henselae* | **N: 39/102**  **A: 6/50** |
|  | Vayssier-Taussat et al. 2013 | Questing | Hseq | *Bartonella* spp. (*B. henselae* and *B. grahamii*) | **N (g. n = 1,450): 1/1** |
| Alsace, Haut-Rhin | Michelet et al. 2014 | Questing | PCRqm | *B.* *henselae* | N (g. n = 25): 0/47 |
|  |  |  |  | *B.* *quintana* | N (g. n = 25): 0/47 |
|  | Nebbak et al. 2019 | Questing | PCRq  + seq | *Bartonella* spp. (probably a new species: 1/5) | **N: 5/50** |
|  |  |  |  | *B.* *henselae* | N: 0/50 |
| Brittany, Côte-d’Armor | Bonnet et al. 2017 | Questing | PCRc  + seq | *Bartonella* spp.  (no sequence obtained) | **N (g. n = 5): 2/86**  F: 0/60  M: 0/85 |
| Burgundy, Côte-d’Or | Bonnet et al. 2013 | Questing | PCRq  + seq | *Bartonella* spp. | F: 0/1 |
| Burgundy, Saône-et-Loire | Bonnet et al. 2013 | Questing | PCRq  + seq | *Bartonella* spp. | F: 0/3 |
| Burgundy, Yonne | Bonnet et al. 2013 | Questing | PCRq  + seq | *Bartonella* spp.  (doubtful species) | **F: 1/3**  M: 0/1 |
| Champagne-Ardennes, Ardennes | Moutailler et al. 2016b | Questing | PCRqm + seq | *B.* *henselae* | **F: 23/267** |
|  |  |  |  | *B.* *quintana* | F: 0/267 |
| Corsica | Grech-Angelini et al. 2020a | Attached (cattle, red deer, wild boar, cat) | PCRqm | *B.* *henselae* | **A: 3/115** |
|  |  |  |  | *B.* *quintana* | A: 0/115 |
| Île-de-France, Essonne | Lejal et al. 2019a | Questing | PCRqm | *Bartonella* spp. | F: 0^b^ /30  M: 0^b^/30 |
|  |  |  |  | *B.* *henselae* | F: 0^b^/30  M: 0^b^/30 |
|  | Lejal et al. 2019b | Questing | PCRqm | *Bartonella* spp. | N: 0/998 |
|  |  |  |  | *B.* *henselae* | N: 0/998 |
|  | Paul et al. 2016 | Questing | PCRc  + seq | *Bartonella* spp.  (no sequence obtained) | **F: 4/259** |
|  | Reis et al. 2011 | Questing | PCRc  + seq | *Bartonella* spp.  (*B.* *birtlesii)* | **N (g. n = 10): 1/36**  F: 0/69  M: 0/122 |
| Midi-Pyrénées, Hautes-Pyrénées | Davoust et al. 2012 | Attached (Pyrenean Chamois) | PCRq | *Bartonella* spp. | 9 N, 60 F, 2 M: 0/71 |
| Nord-Pas-De-Calais, Nord | Halos et al. 2005 | Questing | PCRc  + seq | *Bartonella* spp.  (sequence close to *B.* *schoenbuschensiis*: 1/1) | **N: 3/74**  **F: 3/9**  **M: 3/9** |
| Pays-de-la-Loire, Loire-Atlantique | Cotté et al. 2010 | Questing | PCRc  + seq | *Bartonella* spp.  (related to *B.* *henselae* or *B.* *rochalimae*) | N: 0/267  **F: 1/203**  M: 0/102 |
| Poitou-Charente, Deux-Sèvres | Bonnet et al. 2013 | Questing | PCRq  + seq | *Bartonella* spp.  (doubtful species) | **F: 2/18**  **M: 1/12** |
| Rhône-Alpes, Isère | Bonnet et al. 2013 | Questing | PCRq  + seq | *Bartonella* spp. | F: 0/4  M: 0/2 |

Notes: “Region, department”: French region and department of sampling; “Reference”: source of the data; “Sampling”: sampling of the ticks: Attached: tick sampled on a host with host species in brackets (shaded), Questing: tick sampled while questing for a host; “Method”: detection method used: PCRc: classical PCR followed by gel electrophoresis, PCRq: quantitative PCR, PCRqm: microfluidic quantitative PCR, seq: amplicon sequencing, Hseq: high-throughput sequencing. “*Bartonella* species”: *Bartonella* species searched for; “Positive/tested”: number of positive ticks or pool of ticks on the number of tested ones: L: larvae, N: nymphs, F: adult females, M: adult males, A: adults of unspecified sex, ?: tick stage not specified, “g. n = x”: analyses of pooled DNA of several (x) ticks, /?: unknown number of tested ticks or group of ticks. Positive results are in bold.

^a^ Results of sequence analyses when different from expected by PCR are in brackets.

^b^ Results respectively for salivary glands, gut, and both.

This is a supplementary material for “Perez, G., Bournez, L., Boulanger, N., Fite, J., Livoreil, B., McCoy, K. D., Quillery, E., René-Martellet, M., and Bonnet, S. I. The distribution, phenology, host range and pathogen prevalence of *Ixodes ricinus* in France: a systematic map and a narrative review”, a preprint recommended in PCI Infections <https://doi.org/10.24072/pci.infections.100076>.
