## Supplementary material for "The distribution, phenology, host range and pathogen prevalence of *Ixodes ricinus* in France: a systematic map and narrative review": SI 11

**Supplementary information 11: *Borrelia burgdorferi* s.l. prevalence in *Ixodes ricinus* ticks**

Prevalence of *Borrelia burgdorferi* s.l. in questing *Ixodes ricinus* ticks sampled in European France by tick stage.

| Region, department | Reference | Sampling | Method | *Borrelia* species ^a^ | Positive/tested |
| --- | --- | --- | --- | --- | --- |
| Pooled over several departments: Allier, Ariège, Cantal, Gironde, Landes, and Vaucluse | Geurden et al. 2018 | Attached (cats and dogs) | PCRq  + seq | *B.* *burgdorferi* s.l.: | **A (g. n ≤ 10): 8/52** |
|  |  |  |  | - *B.* *afzelii* | **A (g. n ≤ 10): 2/52** |
|  |  |  |  | - *B.* *burgdorferi* s.s. | **A (g. n ≤ 10): 2/52** |
|  |  |  |  | - *B.* *garinii* | **A (g. n ≤ 10): 3/52** |
|  |  |  |  | - *B.* *valaisiana* | **A (g. n ≤ 10): 1/52** |
| “Eastern” | Gilot et al. 1996 | Questing | IFA | *B.* *burgdorferi* s.l. | **N: 76/1,014** |
| “North-Eastern” | Doby et al. 1991a | Attached (foxes) | IFA | *B.* *burgdorferi* s.l. | N: 0/3  A: 0/2 |
| “North-Western” | Gilot et al. 1996 | Questing | IFA | *B.* *burgdorferi* s.l. | **N: 64/1,422** |
|  | Doby et al. 1989a | Questing (+ few attached) | IFA | *B.* *burgdorferi* s.l. | L: 0/351  **F: 26/180**  **M: 16/186** |
| “South-Western” | Gilot et al. 1996 | Questing | IFA | *B.* *burgdorferi* s.l. | **N: 21/811** |
| Alsace | Richter & Matuschka 2012 | Questing | PCRn  + seq | *B. burgdorferi* s.l.: | **A: 16/60^b^** |
|  |  |  |  | - *B.* *afzelii* | **A: 4/60** |
|  |  |  |  | - *B.* *burgdorferi* s.s. | A: 0/60 |
|  |  |  |  | - *B.* *garinii* | **A: 8/60** |
|  |  |  |  | - *B. lusitaniae* | A: 0/60 |
|  |  |  |  | - *B.* *spielmanii* | A: 0/60 |
|  |  |  |  | - *B.* *valaisiana* | **A: 4/60** |
| Alsace, Bas-Rhin | Boulanger et al. 2018 | Questing | PCRq  + FRET | *B. burgdorferi* s.l. | **N: 5.2-26.7% (/?)** |
|  | Dietrich et al. 2010 | Questing | PCRc  + seq | *B. burgdorferi* s.l. | **N: 2/102**  A: 0/50 |
|  | Michelet et al. 2014 | Questing | PCRqm | *B.* *afzelii* | **N (g. n = 25): 17/47** |
|  |  |  |  | *B.* *bissettii* | N (g. n = 25): 0/47 |
|  |  |  |  | *B.* *burgdorferi* s.s. | **N (g. n = 25): 1/47** |
|  |  |  |  | *B.* *garinii* | **N (g. n = 25): 8/47** |
|  |  |  |  | *B. lusitaniae* | N (g. n = 25): 0/47 |
|  |  |  |  | *B.* *spielmanii* | **N (g. n = 25): 1/47** |
|  |  |  |  | *B.* *valaisiana* | **N (g. n = 25): 1/47** |
|  | Richter et al. 2003 | Questing | PCRn  + seq | *B. burgdorferi* s.l.: | **N: 8/45^b^**  **A: 81/203^b^** |
|  |  |  |  | - *B.* *afzelii* | **N: 5/45^b^**  **A: 30/203^b^** |
|  |  |  |  | - *B. burgdorferi* s.s. | N: 0/45  **A: 1/203^b^** |
|  |  |  |  | - *B.* *garinii* | **N: 1/45^b^**  **A: 21/203^b^** |
|  |  |  |  | - *B.* *lusitaniae* | **N: 2/45^b^**  **A: 18/203^b^** |
|  |  |  |  | - *B.* *valaisiana* | **N: 2/45^b^**  **A: 6/203^b^** |
|  | Vayssier-Taussat et al. 2013 | Questing | Hseq | *Borrelia* spp.: *B.* *afzelii*, *B.* *garinii* and *B.* *burgdorferi* s.s. | **N (g. n = 1,450): 1/1** |
| Alsace, Haut-Rhin | Boulanger et al. 2018 | Questing | PCRq  + FRET | *B. burgdorferi* s.l. | **N: 0.7-23.5% (/?)** |
|  | Boyer et al. 2017 | Questing | PCRq | *B. burgdorferi* s.l.: | **N: 12/81** |
|  |  |  |  | - *B.* *afzelii* | **N: 6/81^c^** |
|  |  |  |  | - *B. burgdorferi* s.s. | **N: 4/81^c^** |
|  |  |  |  | - *B.* *garinii* | **N: 3/81^c^** |
|  | Ferquel et al. 2006 (Goldstein 2017) | Questing | PCRc  + RFLP | *B. burgdorferi* s.l.: | **N: 406/2,296**  **A: 361/1,459** |
|  |  |  |  | - *B.* *afzelii* | **N: 212/2,296^b^**  **A: 63/1,459^b^** |
|  |  |  |  | - *B. burgdorferi* s.s. | **N: 22/2,296^b^**  **A: 39/1,459^b^** |
|  |  |  |  | - *B.* *garinii* | **N: 112/2,296^b^**  **A: 164/1,459^b^** |
|  |  |  |  | - *B.* *lusitaniae* | N: 0/2,296  A: 0/1,459 |
|  |  |  |  | - *B.* *spielmanii* | **N: 1/2,296**  A: 0/1,459 |
|  |  |  |  | - *B.* *valaisiana* | **N: 54/2,296**  **A: 71/1,459** |
|  | Michelet et al. 2014 | Questing | PCRqm | *B.* *afzelii* | **N (g. n = 25): 13/47** |
|  |  |  |  | *B.* *bisettii* | N (g. n = 25): 0/47 |
|  |  |  |  | *B.* *burgdorferi* s.s. | N (g. n = 25): 0/47 |
|  |  |  |  | *B.* *garinii* | **N (g. n = 25): 5/47** |
|  |  |  |  | *B. lusitaniae* | N (g. n = 25): 0/47 |
|  |  |  |  | *B.* *spielmanii* | **N (g. n = 25): 1/47** |
|  |  |  |  | *B.* *valaisiana* | **N (g. n = 25): 1/47** |
|  | Nebbak et al. 2019 | Questing | PCRq  + seq | *Borrelia* spp.: | **N: 10/50** |
|  |  |  |  | - *B.* *afzelii* | **N: 1/50** |
|  |  |  |  | - *B.* *burgdorferi* s.s. | **N: 2/50** |
|  |  |  |  | - *B.* *garinii* | **N: 1/50** |
|  |  |  |  | *- B.* *valaisiana* | **N: 2/50** |
|  | Richter et al. 2003 | Questing | PCRn  + seq | *B. burgdorferi* s.l.: | **N: 28/107^b^**  **A: 18/34^b^** |
|  |  |  |  | - *B.* *afzelii* | **N: 8/107^b^**  **A: 6/34^b^** |
|  |  |  |  | - *B. burgdorferi* s.s. | **N: 2/107^b^**  **A: 1/34^b^** |
|  |  |  |  | - *B.* *garinii* | **N: 8/107^b^**  **A: 4/34^b^** |
|  |  |  |  | - *B.* *valaisiana* | **N: 9/107^b^**  **A: 5/34^b^** |
| Auvergne, Allier | Doby et al. 1993a | Questing | IFA | *B. burgdorferi* s.l. | **N: 18/144** |
|  |  |  | PCRc | *B. burgdorferi* s.l. | **N: 15/81** |
| Auvergne,  Puy-de-Dôme | Beytout et al. 2007 | Questing | PCRc  + RFLP | *B. burgdorferi* s.l.: | **N: 92/525**  **A: 152/526** |
|  |  |  |  | - *B.* *afzelii* | **N: 60/525^b^**  **A: 49/526^b^** |
|  |  |  |  | - *B.* *burgdorferi* s.s. | **N: 9/525^b^**  **A: 15/526^b^** |
|  |  |  |  | - *B.* *garinii* | **N: 9/525^b^**  **A: 56/526^b^** |
|  |  |  |  | - *B.* *lusitaniae* | **N: 2/525^b^**  **A: 2/526^b^** |
|  |  |  |  | - *B.* *spielmanii* | N: 0/525  **A: 3/526^b^** |
|  |  |  |  | - *B.* *valaisiana* | **N: 11/525^b^**  **A: 30/526^b^** |
|  | Halos et al. 2006a (Halos et al. 2004) | Questing | PCRc | *B. burgdorferi* s.l. | **N (g. n = 10): 20/55** |
|  |  |  | PCR-T | *B. burgdorferi* s.l. | **N (g. n = 10): 15/55** |
|  | Halos et al. 2010  (Halos et al. 2006b) | Questing | PCRc  + seq | *B. burgdorferi* s.l.  (*B.* *garinii* ou *B.* *afzelii* : 17/17) | **N (g. n = 5): 11/56**  **N (g. n = 10): 64/220**  **N (g. n = 50): 7/20**  **F: 11/102**  **M: 2/123** |
| Brittany,  Côtes d'Armor | Bonnet et al. 2017 | Questing | PCRc  + seq | *B.* *burgdorferi* s.l.  (no sequence obtained) | **N (g. n = 5): 2/86**  F: 0/60  **M: 2/85** |
|  | Doby et al. 1989b | Questing (+ few attached) | IFA | *B. burgdorferi* s.l. | **N: 8/233** |
| Brittany,  Ille-et-Vilaine | Doby et al. 1989b | Questing (+ few attached) | IFA | *B. burgdorferi* s.l. | **N: 54/616** |
|  | Doby et al. 1991b | Questing | IFA | *B. burgdorferi* s.l. | **N: 50/357** |
|  | Doby et al. 1995 | Questing | IFA | *B. burgdorferi* s.l. | **N: 144/1,506** |
|  | Lebert et al. 2020 | Questing | PCRq  + seq | *B. burgdorferi* s.l.: | **N: 78/2,627** |
|  |  |  |  | - *B.* *afzelii* | **N: 16/2,627** |
|  |  |  |  | - *B.* *burgdorferi* s.s. | **N: 13/2,627** |
|  |  |  |  | - *B.* *garinii* | **N: 20/2,627** |
|  |  |  |  | - *B. lusitaniae?* | **N: 1/2,627** |
|  |  |  |  | - *B.* *spielmanii* | **N: 1/2,627** |
|  |  |  |  | - *B.* *valaisiana* | **N: 14/2,627** |
|  |  |  |  | - *B.* sp. | **N: 7/2,627** |
|  |  |  |  | - *B.* spp. coinfection | **N: 6/2,627** |
| Brittany, Morbihan | Doby et al. 1989b | Questing (+ few attached) | IFA | *B. burgdorferi* s.l. | **N: 5/123** |
| Burgundy,  Côte-d’Or | Bonnet et al. 2013 | Questing | PCRc  + seq | *B. burgdorferi* s.l. | F: 0/1 |
| Burgundy,  Saône-et-Loire | Bonnet et al. 2013 | Questing | PCRc  + seq | *B. burgdorferi* s.l. | **F: 1/3** |
| Burgundy, Yonne | Bonnet et al. 2013 | Questing | PCRc  + seq | *B. burgdorferi* s.l. | **F: 1/3**  M: 0/1 |
| Centre,  Loir-et-Cher | Doby et al. 1993a | Questing | IFA | *B. burgdorferi* s.l. | **N: 9/82** |
|  |  |  | PCRc | *B. burgdorferi* s.l. | **N: 2/15** |
| Champagne-Ardenne, Ardennes | Moutailler et al. 2016b | Questing | PCRqm | *B.* *afzelii* | **F: 21/267** |
|  |  |  |  | *B.* *bisettii* | F: 0/267 |
|  |  |  |  | *B.* *burgdorferi* s.s. | **F: 13/267** |
|  |  |  |  | *B.* *garinii* | **F: 25/267** |
|  |  |  |  | *B. lusitaniae* | F: 0/267 |
|  |  |  |  | *B.* *spielmanii* | **F: 5/267** |
|  |  |  |  | *B.* *valaisiana* | **F: 15/267** |
| Champagne-Ardenne, Marne | Ferté et al. 1994  (Ulmer et al. 1999) | Questing | Culture + PCRc + RFLP | *B.* *burgdorferi* s.l.  (*B.* *afzelii* for 1 group of 2 positive ticks) | **N: 3/54**  F: 0/58  **M: 1/24** |
| Corsica | Cicculli et al. 2019b | Attached (black rats) | PCRq | *B.* *burgdorferi* s.l. | **F: 2/13** |
|  | Grech-Angelini et al. 2020a | Attached (cattle, red deer, wild boar, cat) | PCRqm | *B.* *afzelii* | **A: 4/115** |
|  |  |  |  | *B.* *bissetti* | A: 0/115 |
|  |  |  |  | *B.* *burgdorferi* s.s. | A: 0/115 |
|  |  |  |  | *B.* *garinii* | A: 0/115 |
|  |  |  |  | *B. lusitaniae* | A: 0/115 |
|  |  |  |  | *B.* *spielmanii* | A: 0/115 |
|  |  |  |  | *B.* *valaisiana* | A: 0/115 |
| Île-de-France, Essonne | Lejal et al. 2019a | Questing | PCRqm | *B.* *afzelii* | **F: 2 (0-2-0)^d^/30**  M: 0^d^ /30 |
|  |  |  |  | *B.* *bisettii* | F: 0^d^ /30  M: 0^d^ /30 |
|  |  |  |  | *B.* *burgdorferi* s.s. | F: 0^d^ /30  M: 0^d^ /30 |
|  |  |  |  | *B.* *garinii* | **F: 1 (0-0-1)^d^/30**  M: 0^d^ /30 |
|  |  |  |  | *B. lusitaniae* | **F: 2 (0-0-2)^d^/30**  M: 0^d^ /30 |
|  |  |  |  | *B.* *spielmanii* | **F: 2 (0-1-1)^d^/30**  M: 0^d^ /30 |
|  |  |  |  | *B.* *valaisiana* | F: 0^d^ /30  M: 0^d^ /30 |
|  | Lejal et al. 2019b | Questing | PCRqm | *B.* *afzelii* | **N: 11/998** |
|  |  |  |  | *B.* *bisettii* | N: 0/998 |
|  |  |  |  | *B.* *burgdorferi* s.s. | **N: 15/998** |
|  |  |  |  | *B.* *garinii* | **N: 11/998** |
|  |  |  |  | *B. lusitaniae* | N: 0/998 |
|  |  |  |  | *B.* *spielmanii* | **N: 4/998** |
|  |  |  |  | *B.* *valaisiana* | **N: 6/998** |
|  | Marchant et al. 2017 | Questing | PCRn/c + RFLP | *B.* *burgdorferi* s.l.: | **N: 519/4,973**  **A: 123/1,226** |
|  |  |  |  | *- B.* *afzelii* | **N: 181/4,973**  **A: 25/1,226** |
|  |  |  |  | *- B.* *burgdorferi* s.s. | **N: 150/4,973**  **A: 34/1,226** |
|  |  |  |  | *- B. garinii* | **N: 87/4,973**  **A: 32/1,226** |
|  |  |  |  | *- B. lusitaniae* | **N: 21/4,973**  A: 0/1,226 |
|  |  |  |  | *- B.* *spielmanii* | **N: 9/4,973**  **A: 9/1,226** |
|  |  |  |  | *- B.* *valaisiana* | **N: 48/4,973**  **A: 11/1,226** |
|  |  |  |  | *- B.* spp. coinfection | **N: 23/4,973**  **A: 6/1,226** |
|  | Paul et al. 2016 | Questing | PCRc  + seq | *B.* *burgdorferi* s.l. | **F: 20/259** |
|  | Reis et al. 2011b | Questing | PCRc  + seq | *B.* *burgdorferi* s.l. | **N (g. n = 10): 24/36**  **F: 19/69 ^d^**  **M: 24/122** |
|  | Vourc’h et al. 2016 | Questing | PCRq | *B.* *burgdorferi* s.l. | **N: 394/3,903** |
| Île-de-France, Seine-et-Marne | Zhioua et al. 1996 | Questing | IFA | *B.* *burgdorferi* s.l. | **N: 22/195**  **F: 1/12**  **M: 1/10** |
| Île-de-France, Val-de-Marne | Marchant et al. 2017 | Questing | PCRn/c + RFLP | *B.* *burgdorferi* s.l. | **N: 32/326**  **A: 4/46** |
| Île-de-France, Yvelines | Marchant et al. 2017 | Questing | PCRn/c + RFLP | *B.* *burgdorferi* s.l. | **N: 109/1,234**  **A: 42/239** |
|  | Pichon et al. 1999 | Questing | PCRc  + RFLP | *B.* *burgdorferi* s.l. | **N: 38/461** |
|  |  |  |  | *B.* *afzelii* | **N: 20/461^c^** |
|  |  |  |  | *B.* *burgdorferi* s.s. | **N: 4/461^c^** |
|  |  |  |  | *B.* *garinii* | **N: 11/461^c^** |
|  | Zhioua et al. 1996 | Questing | IFA | *B.* *burgdorferi* s.l. | **N: 17/119**  F: 0/22  M: 0/25 |
| Limousin, Corrèze | Doby et al. 1993a | Questing | PCRc | *B. burgdorferi* s.l. | **N: 2/15** |
| Lorraine, Meuse | Beytout et al. 2007 | Questing | PCRc  + RFLP | *B.* *burgdorferi* s.l.: | **N: 33/434**  **A: 31/164** |
|  |  |  |  | - *B.* *afzelii* | **N: 20/434^b^**  **F: 2/164^b^** |
|  |  |  |  | - *B.* *burgdorferi* s.s. | **N: 1/434^b^**  **F: 1/164^b^** |
|  |  |  |  | - *B.* *garinii* | **N: 9/434^b^**  **F: 8/164^b^** |
|  |  |  |  | - *B.* *lusitaniae* | N: 0/434  F: 0/164 |
|  |  |  |  | - *B.* *spielmanii* | N: 0/434  F: 0/164 |
|  |  |  |  | - *B.* *valaisiana* | **N: 3/434^b^**  **F: 13/164^b^** |
| Midi-Pyrénées, Haute-Garonne | Ehrmann et al. 2018 | Questing | PCRq | *B.* *burgdorferi* s.l | **N (g. n ≤ 10): 8 % (/?)**  **A (g. n ≤ 10): 3 % (/?)** |
|  | Lebert et al. 2020 | Questing | PCRq  + seq | *B. burgdorferi* s.l.: | **N: 47/1,891** |
|  |  |  |  | - *B.* *afzelii* | **N: 8/1,891** |
|  |  |  |  | - *B.* *burgdorferi* s.s. | **N: 15/1,891** |
|  |  |  |  | - *B.* *garinii* | **N: 6/1,891** |
|  |  |  |  | - *B.* *valaisiana* | **N: 10/1,891** |
|  |  |  |  | - *B.* *burgdorferi* sp. | **N: 4/1,891** |
|  |  |  |  | - *B.* spp. coinfection | **N: 4/1,891** |
| Midi-Pyrénées, Hautes-Pyrénées | Akl et al. 2019 | Questing | PCRc  + seq | *B. burgdorferi* s.l.:  (*B.* *afzelii* : 5/15; *B.* *burgdorferi* s.s.: ?/15 and *B.* *garinii*: ?/15) | **N: 52/531**  **F: 3/86**  **M: 4/79** |
| Nord-Pas-De-Calais, Nord | Halos et al. 2005 | Questing | PCRc | *B.* *burgdorferi* s.l. | **N: 1/74**  **F: 1/9**  **M: 1/9** |
| Lower Normandy, Manche | Doby et al. 1989b | Questing (+ few attached) | IFA | *B. burgdorferi* s.l. | **N: 2/6** |
| Lower Normandy, Orne | Doby et al. 1989b | Questing (+ few attached) | IFA | *B. burgdorferi* s.l. | **N: 14/138** |
| Picardy | Ehrmann et al. 2018 | Questing | PCRq | *B.* *burgdorferi* s.l. | **N (g. n ≤ 10): 13% (/?)**  **A (g. n ≤ 10): 24% (/?)** |
| Pays-de-la-Loire, Loire-Atlantique | Cotté et al. 2010 | Questing | PCRc  + seq | *B.* *burgdorferi* s.l.  (*B.* *afzelii* or *B.* *garinii*; *B.* *carolinensis*?: 1) | **N: 5/267**  **F: 27/203**  **M: 3/102** |
|  | Doby et al. 1991b | Questing | IFA | *B. burgdorferi* s.l. | **N: 32/320** |
|  | Doby et al. 1989b | Questing (+ few attached) | IFA | *B. burgdorferi* s.l. | **N: 18/210** |
| Pays-de-la-Loire, Maine-et-Loire | Doby et al. 1989b | Questing (+ few attached) | IFA | *B. burgdorferi* s.l. | **N: 3/78** |
| Pays-de-la-Loire, Mayenne | Doby et al. 1989b | Questing (+ few attached) | IFA | *B. burgdorferi* s.l. | **N: 3/38** |
| Pays-de-la-Loire, Sarthe | Doby et al. 1989b | Questing (+ few attached) | IFA | *B. burgdorferi* s.l. | **N: 12/158** |
| Poitou-Charentes, Deux-Sèvres | Bonnet et al. 2013 | Questing | PCRc  + seq | *B.* *burgdorferi* s.l. | **F: 0/18**  **M: 0/12** |
|  | Doby et al. 1993 | Questing | IFA | *B.* *burgdorferi* s.l. | **N: 11/72** |
| Rhône-Alpes, Ain | Quessada et al. 2003 | Questing | PCRn  + RFLP | *B.* *burgdorferi* s.l.: | L: 0/50  **N: 9/50**  **F: 7/25**  **M: 2/25** |
|  |  |  |  | - *B.* *afzelii* | **N: 6/50**  **F: 4/25**  M: 0/25 |
|  |  |  |  | - *B.* *burgdorferi* s.s. | **N: 2/50**  **F: 1/25**  M: 0/25 |
|  |  |  |  | - *B.* *garinii* | **N: 1/50**  **F: 2/25**  **M: 1/25** |
|  |  |  |  | - *B. lusitaniae* | N: 0/50  F: 0/25  M: 0/25 |
|  |  |  |  | - *B.* *valaisiana* | N: 0/50  F: 0/25  **M: 1/25** |
| Rhône-Alpes, Isère | Bonnet et al. 2013 | Questing | PCRc  + seq | *B.* *burgdorferi* s.l. | F: 0/4  M: 0/2 |
| Rhône-Alpes, Loire | Quessada et al. 2003 | Questing | PCRn  + RFLP | *B.* *burgdorferi* s.l.: | **N: 4/50** |
|  |  |  |  | - *B.* *afzelii* | N: 0/50 |
|  |  |  |  | - *B.* *burgdorferi* s.s. | N: 0/50 |
|  |  |  |  | - *B.* *garinii* | **N: 2/50^c^** |
|  |  |  |  | - *B. lusitaniae* | N: 0/50 |
|  |  |  |  | - *B.* *valaisiana* | **N: 3/50^c^** |
| Rhône-Alpes, Rhône | Quessada et al. 2003 | Questing | PCR  + RFLP | *B.* *burgdorferi* s.l.: | **N: 35/207**  **F: 6/49**  **M: 4/54** |
|  |  |  |  | - *B.* *afzelii* | **N: 19/207^c^**  **F: 1/49**  **M: 1/54** |
|  |  |  |  | - *B.* *burgdorferi* s.s. | **N: 2/207^c^**  F: 0/49  M: 0/54 |
|  |  |  |  | - *B.* *garinii* | **N: 8/207**  **F: 3/49**  **M: 2/54** |
|  |  |  |  | - *B. lusitaniae* | N: 0/207  F: 0/49  M: 0/54 |
|  |  |  |  | - *B.* *valaisiana* | **N: 5/207**  **F: 2/49**  **M: 1/54** |
| Rhône-Alpes, Rhône (Lyon city) | Quessada et al. 2003 | Questing | PCRn  + RFLP | *B.* *burgdorferi* s.l.: | **N: 11/112**  **F: 5/28**  **M: 8/38** |
|  |  |  |  | - *B.* *afzelii* | **N: 2/112^c^**  **F: 3/28^c^**  M: 0/38 |
|  |  |  |  | - *B.* *burgdorferi* s.s. | N: 0/112  F: 0/28  M: 0/38 |
|  |  |  |  | - *B.* *garinii* | **N: 4/112^c^**  **F: 1/28^c^**  **M: 4/38^c^** |
|  |  |  |  | - *B. lusitaniae* | N: 0/112  F: 0/28  M: 0/38 |
|  |  |  |  | - *B.* *valaisiana* | **N: 8/112^c^**  **F: 2/28^c^**  **M: 5/38^c^** |

Notes: “Region, department”: French region and department of sampling; “Reference”: source of the data; “Sampling”: sampling of the ticks: Attached: tick sampled on a host with host species in brackets (shaded), Questing: tick sampled while questing for a host; “Method”: detection method used: IFA: immunofluorescence assay, PCR: polymerisation chain reaction, PCRc: classical PCR followed by gel electrophoresis, PCRn: nested PCR, PCRq: quantitative PCR, PCRqm: microfluidic quantitative PCR, PCR-T: Temporal Temperature Gradient gel Electrophoresis PCR, PCR-FRET: PCR using fluorescence resonance energy transfer hybridization probe method, RFLP: amplicon analyses with a restriction fragment length polymorphism method, seq: amplicon sequencing, Hseq: high-throughput sequencing; “*Borrelia* species”: *Borrelia burgdorferi* s.l. species searched for; “Positive/tested”: number of positive ticks or pool of ticks on the number of tested ones: L: larvae, N: nymphs, F: adult females, M: adult males, A: adults of unspecified sex, ?: tick stage not specified, “g. n = x”: analyses of pooled DNA of several (x) ticks, /?: unknown number of tested ticks or group of ticks. Positive results are in bold.

^a^ Results of sequence analyses when different from expected by PCR are in brackets.

^b^ Estimate from percentages.

^c^ Including co-infections, explaining a total number of genospecies exceeding number of total positive ticks.

^d^ Results respectively for salivary glands alone, gut alone, and both.

^e^ Adult females infected with *B.* *miyamotoi* have been excluded from presented results.

This is a supplementary material for “Perez, G., Bournez, L., Boulanger, N., Fite, J., Livoreil, B., McCoy, K. D., Quillery, E., René-Martellet, M., and Bonnet, S. I. The distribution, phenology, host range and pathogen prevalence of *Ixodes ricinus* in France: a systematic map and a narrative review”, a preprint recommended in PCI Infections <https://doi.org/10.24072/pci.infections.100076>.
