## Supplementary material for "The distribution, phenology, host range and pathogen prevalence of *Ixodes ricinus* in France: a systematic map and narrative review": SI 12

**Supplementary information 12: *Borrelia miyamotoi* prevalence in *Ixodes ricinus* ticks**

Prevalence of *Borrelia miyamotoi* in questing *Ixodes ricinus* ticks sampled in European France by tick stage.

| Region, department | Reference | Sampling | Method | *Borrelia* RF species | Positive/tested |
| --- | --- | --- | --- | --- | --- |
| Alsace, Bas-Rhin | Boulanger et al. 2018 | Questing | PCRq  + seq | *B. burgdorferi* s.l. | **N: 0.75-4.2% (/?)** |
|  | Richter et al. 2003 | Questing | PCRn  + seq | *Borrelia* RF | N: 0/45^a^  **A: 7/203^a^** |
|  | Vayssier-Taussat et al. 2013 | Questing | Hseq | *Borrelia miyamotoi* | **N (g. n = 1,450): 1/1** |
| Alsace, Haut-Rhin | Boyer et al. 2020 | Questing | PCRq | *Borrelia miyamotoi* | **N: 94/4,354** |
|  | Richter et al. 2003 | Questing | PCRn  + seq | *Borrelia* RF | **N: 3/107^a^**  **A: 3/34^a^** |
|  | Nebbak et al. 2019 | Questing | PCRq  + seq | *Borrelia miyamotoi* | **N: 1/50** |
| Champagne-Ardenne, Ardennes | Cosson et al. 2014 | Questing | PCRq | *Borrelia miyamotoi* | **F^b^: 8/267** |
|  | Moutailler et al. 2016b | Questing | PCRqm | *Borrelia miyamotoi* | **F^ab^: 8/267** |
| Corsica | Cicculli et al. 2019b | Attached (black rats) | PCRq | *Borrelia miyamotoi* | F: 0/13 |
|  | Grech-Angelini et al. 2020a | Attached (cattle, red deer, wild boar, cat) | PCRqm | *Borrelia miyamotoi* | **A: 2/115** |
| Île-de-France, Essonne | Lejal et al. 2019a | Questing | PCRqm | *Borrelia miyamotoi* | F: 0^c^ /30  **M: 13 (10-0-3)^c^/30** |
|  | Lejal et al. 2019a | Questing | PCRqm | *Borrelia miyamotoi* | **N: 12/998** |
|  | Paul et al. 2016 | Questing | PCRc  + seq | *Borrelia miyamotoi* | **F: 2/259** |

Notes: “Region, department”: French region and department of sampling; “Reference”: source of the data; “Sampling”: sampling of the ticks: Attached: tick sampled on a host with host species in brackets (shaded), Questing: tick sampled while questing for a host; “Method”: PCR: polymerisation chain reaction, PCRc: classical PCR followed by gel electrophoresis, PCRn: nested PCR, PCRq: quantitative PCR, PCRqm: microfluidic quantitative PCR, seq: amplicon sequencing, Hseq: high-throughput sequencing; “*Borrelia* RF species”: species of the *Borrelia* relapsing fever (RF) group searched for, it is either *Borrelia miyamotoi* or the clade of *Borrelia* RF group without species identification; “Positive/tested”: number of positive ticks or pool of ticks on the number of tested ones: L: larvae, N: nymphs, F: adult females, M: adult males, A: adults of unspecified sex, ?: tick stage not specified, “g. n = x”: analyses of pooled DNA of several (x) ticks, /?: unknown number of tested ticks or group of ticks. Positive results are in bold.

^a^ Estimate from percentages.

^b^ Same ticks analysed with different methods.

^c^ Results respectively for salivary glands alone, gut alone, and both.

This is a supplementary material for “Perez, G., Bournez, L., Boulanger, N., Fite, J., Livoreil, B., McCoy, K. D., Quillery, E., René-Martellet, M., and Bonnet, S. I. The distribution, phenology, host range and pathogen prevalence of *Ixodes ricinus* in France: a systematic map and a narrative review”, a preprint recommended in PCI Infections <https://doi.org/10.24072/pci.infections.100076>.
