## Supplementary material for "The distribution, phenology, host range and pathogen prevalence of *Ixodes ricinus* in France: a systematic map and narrative review": SI 13

**Supplementary information 13: *Coxiella burnetii* prevalence in *Ixodes ricinus* ticks**

Prevalence of *Coxiella burnetii* in questing *Ixodes ricinus* ticks sampled in European France by tick stage.

| Region, department | Reference | Sampling | Method | *Coxiella* species ^a^ | Positive/tested |
| --- | --- | --- | --- | --- | --- |
| Alsace, Bas-Rhin | Michelet et al. 2014 | Questing | PCRqm | *C*. *burnetii* | N (g. n = 25): 0/47 |
|  | Vayssier-Taussat et al. 2013 | Questing | Hseq | *Coxiella* spp. | **N (g. n = 1,450): 1/1** |
|  |  |  | Hseq  + PCRq | *C*. *burnetii* | N (g. n = 25): 0/58  A (g. n = 2): 0/31 |
| Alsace, Haut-Rhin | Michelet et al. 2014 | Questing | PCRqm | *C*. *burnetii* | N (g. n = 25): 0/47 |
|  | Nebbak et al. 2019 | Questing | PCRq  + seq | *C*. *burnetii* | N: 0/50 |
| Burgundy,  Côte-d’Or | Bonnet et al. 2013 | Questing | PCRc | *C*. *burnetii* | F: 0/1 |
| Burgundy,  Saône-et-Loire | Bonnet et al. 2013 | Questing | PCRc | *C*. *burnetii* | **F: 1/3** |
| Burgundy, Yonne | Bonnet et al. 2013 | Questing | PCRc | *C*. *burnetii* | **F: 1/3**  **M: 1/1** |
| Corsica | Grech-Angelini et al. 2020a | Attached (cattle, red deer, wild boar, cat) | PCRqm | *Coxiella* spp. | A: 0/115 |
|  |  |  |  | *C*. *burnetii* | A: 0/115 |
| Île-de-France, Essonne | Lejal et al. 2019a | Questing | PCRqm | *C*. *burnetii* | F: 0^b^ /30  M: 0^b^/30 |
|  | Lejal et al. 2019b | Questing | PCRqm | *C*. *burnetii* | N: 0/988 |
| Poitou-Charente,  Deux-Sèvres | Bonnet et al. 2013 | Questing | PCRc | *C*. *burnetii* | **F: 2/18**  **M: 1/12** |
| Rhône-Alpes, Isère | Bonnet et al. 2013 | Questing | PCRc | *C*. *burnetii* | **F: 1/4**  **M: 1/2** |

Notes: “Region, department”: French region and department of sampling; “Reference”: source of the data; “Sampling”: sampling of the ticks: Attached: tick sampled on a host with host species in brackets (shaded), Questing: tick sampled while questing for a host; “Method”: PCR: Polymerisation Chain Reaction, PCRc: classical PCR followed by gel electrophoresis, PCRqm: microfluidic quantitative PCR, seq: amplicon sequencing, Hseq: high-throughput sequencing; “*Coxiella* species”: *Coxiella* species searched for, it is either *C*. *burnetii* or the genus *Coxiella* without species identification; “Positive/tested”: number of positive ticks or pool of ticks on the number of tested ones: L: larvae, N: nymphs, F: adult females, M: adult males, A: adults of unspecified sex, ?: tick stage not specified, “g. n = x”: analyses of pooled DNA of several (x) ticks, /?: unknown number of tested ticks or group of ticks. Positive results are in bold.

^a^ Results of sequence analyses when different from expected by PCR are in brackets.

^b^ Results for salivary glands, gut and both.

This is a supplementary material for “Perez, G., Bournez, L., Boulanger, N., Fite, J., Livoreil, B., McCoy, K. D., Quillery, E., René-Martellet, M., and Bonnet, S. I. The distribution, phenology, host range and pathogen prevalence of *Ixodes ricinus* in France: a systematic map and a narrative review”, a preprint recommended in PCI Infections <https://doi.org/10.24072/pci.infections.100076>.
