## Supplementary material for "The distribution, phenology, host range and pathogen prevalence of *Ixodes ricinus* in France: a systematic map and narrative review": SI 15

**Supplementary information 15: *Rickettsia* spp. prevalence in *Ixodes ricinus* ticks**

Prevalence of *Rickettsia* spp. in questing *Ixodes* *ricinus* ticks sampled in European France.

| Region, department | Reference | Sampling | Method | *Rickettsia* species ^a^ | Positive/tested |
| --- | --- | --- | --- | --- | --- |
| Alsace, Bas-Rhin | Michelet et al. 2014 | Questing | PCRqm | *Rickettsia* spp. SFG^b^ | **N (g. n = 25): 46/47** |
|  |  |  |  | *R. helvetica* | **N (g. n = 25): 46/47** |
|  |  |  |  | *R. conorii*, *R. massiliae* and *R. slovaca* | N (g. n = 25): 0/47 |
|  | Vayssier-Taussat et al. 2013 | Questing | Hseq | *Rickettsia* spp. SFG^b^ | **N (g. n = 1,450): 1/1** |
|  |  |  | Hseq | *R. helvetica* | **N (g. n = 1,450): 1/1** |
|  |  |  | Hseq | *R. felis* | **N (g. n = 1,450): 1/1** |
|  |  |  | PCRq | *R. felis* | **N (g. n = 25): 1/58**  A (g. n = 2): 0/31 |
| Alsace, Haut-Rhin | Michelet et al. 2014 | Questing | PCRqm | *Rickettsia* spp. SFG^b^ | **N (g. n = 25): 46/47** |
|  |  |  |  | *R. helvetica* | **N (g. n = 25): 46/47** |
|  |  |  |  | *R. conorii*, *R. massiliae* and *R. slovaca* | N (g. n = 25): 0/47 |
|  | Nebbak et al. 2019 | Questing | PCRq  + seq | *Rickettsia* spp. | **N: 3/50** |
|  |  |  |  | - *R.* *helvetica* | **N: 3/50** |
| Auvergne,  Puy-de-Dôme | Halos et al. 2006a | Questing | PCRc | *Rickettsia* spp. SFG^b^ | **N (g. n = 10): 17/55** |
|  |  |  | PCR-T | *Rickettsia* spp. SFG^b^ | **N (g. n = 10): 16/55** |
|  | Halos et al. 2010  (Halos et al. 2006b) | Questing | PCRc  + seq | *Rickettsia* spp.  (*R.* *helvetica*: 12/12) | **N (g. n = 5): 9/56**  **N (g. n = 10): 49/220**  **N (g. n = 50): 9/20**  **F: 8/102**  **M: 6/123** |
|  | Parola et al. 1998a |  | PCRc  + WB  + culture | *R. helvetica* | **A: 2/80** |
| Brittany,  Côte-d’Armor | Bonnet et al. 2017 | Questing | PCRc  + seq | *Rickettsia* spp. SFG^b^  (*R.* *helvetica*) | **N (g. n = 5): 8/86**  F: 0/60  **M: 9/85** |
| Champagne-Ardenne, Ardennes | Moutailler et al. 2016b | Questing | PCRqm | *R. helvetica* | **F: 43/267** |
|  |  |  |  | *R. conorii*, *R. massiliae* and *R. slovaca* | F: 0/267 |
| Corsica | Grech-Angelini et al. 2020a | Attached (cattle, red deer, wild boar, cat) | PCRqm | *R.* *helvetica* | **A: 7/115** |
|  |  |  |  | *R. aeschlimannii*, *R.* *conorii,* *R.* *massiliae* and *R.* *slovaca* | A: 0/115 |
| Corsica, Haute-Corse | Cicculli et al. 2019b | Attached (black rats) | PCRq/c  + seq | *Rickettsia* spp. | F: 0/13 |
| Île-de-France, Essonne | Lejal et al. 2019a | Questing | PCRqm | *R. felis* | **F: 2 (2-0-0)^c^/30**  **M: 2 (2-0-0)^c^/30** |
|  |  |  |  | *R. helvetica* | **F: 5 (1-1-3)^c^/30**  **M: 10 (2-4-4)^c^/30** |
|  |  |  |  | *R. aeschlimannii*, *R.* *conorii,* *R.* *massiliae* and *R.* *slovaca* | F: 0^c^/30  M: 0^c^/30 |
|  | Lejal et al. 2019b | Questing | PCRqm | *R. felis* | **N: 1/998** |
|  |  |  |  | *R. helvetica* | **N: 45/998** |
|  |  |  |  | *R. aeschlimannii*, *R.* *conorii,* *R. massiliae* and *R.* *slovaca* | N: 0/998 |
|  | Paul et al. 2016 | Questing | PCRc  + seq | *Rickettsia* spp. SFG^b^ (*R.* *helvetica*) | **F: 6/259** |
|  | Reis et al. 2011 | Questing | PCRc  + seq | *Rickettsia* spp. SFG^b^ (*R.* *helvetica*) | **N (g. n = 10): 13/36**  **F: 5/69**  **M: 7/122** |
| Midi-Pyrénées,  Hautes-Pyrénées | Akl et al. 2019 | Questing | PCRc  + seq | *Rickettsia* spp.  (*R.* *helvetica:* 2/2) | **N: 75/531**  **F: 28/86**  **M: 20/79** |
|  |  |  |  | *- Rickettsia* spp. SFG^b^:  (*R.* *monacensis*: 5/10) | **N: 4/75**  **F: 4/28**  **M: 2/20** |
|  | Davoust et al. 2012 | Attached (Pyrenean chamois) | PCRq  + seq | *Rickettsia* spp. (*R. helvetica*) | **60F, 2M, 9N: 4/71** |
| Pays-de-la-Loire, Loire-Atlantique | Cotté et al. 2010 | Questing | PCRc  + seq | *Rickettsia* spp. SFG^b^  (*R.* *helvetica*) | **N: 3/267**  **F: 4/203**  **M: 1/102** |

Notes: “Region, department”: French region and department of sampling; “Reference”: source of the data; “Sampling”: sampling of the ticks: Attached: tick sampled on a host with host species in brackets (shaded), Questing: tick sampled while questing for a host; “Method”: PCR: Polymerisation Chain Reaction, PCRc: classical PCR followed by gel electrophoresis, PCRq: quantitative PCR, PCRqm: microfluidic quantitative PCR, seq: amplicon sequencing, Hseq: high-throughput sequencing, WB: western blot, culture: cell culture; “*Rickettsia* species”: *Rickettsia* species searched for; “Positive/tested”: number of positive ticks or pool of ticks on the number of tested ones: L: larvae, N: nymphs, F: adult females, M: adult males, A: adults of unspecified sex, ?: tick stage not specified, “g. n = x”: analyses of pooled DNA of several (x) ticks, /?: unknown number of tested ticks or group of ticks. Positive results are in bold.

^a^ Results of sequence analyses when different from expected by PCR are in brackets.

^b^ SFG: spotted fever group.

^c^ Results respectively for salivary glands, gut, and both.

This is a supplementary material for “Perez, G., Bournez, L., Boulanger, N., Fite, J., Livoreil, B., McCoy, K. D., Quillery, E., René-Martellet, M., and Bonnet, S. I. The distribution, phenology, host range and pathogen prevalence of *Ixodes ricinus* in France: a systematic map and a narrative review”, a preprint recommended in PCI Infections <https://doi.org/10.24072/pci.infections.100076>.
