## Supplementary material for "The distribution, phenology, host range and pathogen prevalence of *Ixodes ricinus* in France: a systematic map and narrative review": SI 16

**Supplementary information 16: *Babesia* spp. and *Theileria* spp. prevalence in *Ixodes ricinus* ticks**

Prevalence of *Babesia* spp. and *Theileria* spp. in questing *Ixodes ricinus* ticks sampled in European France by tick stage.

| Region, department | Reference | Sampling | Method | *Babesia* spp.-*Theileria* spp. ^a^ | Positive/tested |
| --- | --- | --- | --- | --- | --- |
| Alsace,  Bas-Rhin | Michelet et al. 2014 | Questing | PCRqm | *B.* *venatorum* | **N (g. n = 25): 3/47** |
|  |  |  |  | *B. bigemin*a, *B. bovi*s, *B.* *caballi*, *B.* *canis*, *B.* *divergens*, *B.* *major*, *B.* *microti*, *B.* *ovis*, *B. vogeli*, *Th*. *equi* and *Th*. *annulata* | N (g. n = 25): 0/47 |
| Alsace,  Haut-Rhin | Michelet et al. 2014 | Questing | PCRqm | *B.* *venatorum* | **N (g. n = 25): 2/47** |
|  |  |  |  | *B. bigemin*a, *B. bovi*s, *B. caballi*, *B. canis*, *B. divergens*, *B. major*, *B. microti*, *B. ovis*, *B. vogeli*, *Th*. *equi* and *Th*. *annulata* | N (g. n = 25): 0/47 |
|  | Nebbak et al. 2019 | Questing | PCRq  + seq | *Babesia* spp.:  *- B.* *venatorum* | **N: 1/50** |
| Brittany,  Côte-d’Armor | Bonnet et al. 2017 | Questing | PCRc  + seq | *Babesia* spp.-*Theileria* spp. | N (g. n 5): 0/86  F: 0/60  M: 0/85 |
| Brittany,  Ille-et-Vilaine | Lebert et al. 2020 (Jouglin et al. 2017b) | Questing | PCRn  + seq | *Babesia spp.:*  *- B. capreoli*  *- B. divergens* | **N: 15/2,620**  **N: 2/2,620**  **N: 13/2,620** |
| Burgundy,  Yonne | Bonnet et al. 2013 | Questing | PCRc  + RLBH^b^ | *Babesia* spp. and *Theileria* spp. | F: 0/3  M: 0/1 |
| Burgundy,  Côte-d’Or | Bonnet et al. 2013 | Questing | PCRc  + RLBH^b^ | *Babesia* spp. and *Theileria* spp. | F: 0/1 |
| Burgundy,  Saône-et-Loire | Bonnet et al. 2013 | Questing | PCRc  + RLBH^b^ | *Babesia* spp.:  *- B.* *divergens* | **F: 3/3**  **F: 2/3** |
|  |  |  |  | *Theileria* spp. | F: 0/3 |
| Champagne-Ardenne,  Ardennes | Moutailler et al. 2016 | Questing | PCRqm | *B.* *divergens* | **F: 1/267** |
|  |  |  |  | *B.* *caballi*, *B.* *canis*, *B.* *bigemina*, *B. bovis*, *B.* *major*, *B.* *microti*, *B.* *ovis*, *B.* *venatorum*, *B.* *vogeli*, *Th*. *equi* and *Th*. *annulata* | F: 0/267 |
| Corsica | Grech-Angelini et al. 2020a | Attached (cattle, red deer, wild boar, cat) | PCRqm  + seq | *B.* *caballi*, *B.* *canis*, *B.* *bigemina*, *B.* *bovis*, *B.* *divergens*, *B.* *major*, *B.* *microti*, *B.* *Ovis*, *B.* *venatorum*, *B.* *vogeli*, *Th*. *equi* and *Th*. *annulata* | A: 0/115 |
| Île-de-France, Essonne | Lejal et al. 2019a | Questing | PCRqm | *B. venatorum* | F: 0/30  **M: 6 (4-0-2)^c^/30** |
|  |  |  |  | *B. bovis*, *B.* *caballi*, *B.* *canis*, *B. divergens*, *B. microti*, *B.* *ovis* and *Theileria* spp. | F: 0/30  M: 0/30 |
|  | Lejal et al. 2019b | Questing | PCRqm  + seq | *B.* *venatorum* | **N: 15/998** |
|  |  |  |  | *B.* *divergens* (*B.* *capreoli*^d^) | **N: 1/998** |
|  |  |  |  | *B. bovis*, *B. caballi*, *B.* *canis*, *B. microti*, *B.* *ovis* and *Theileria* spp. | N: 0/998 |
|  | Paul et al. 2016 | Questing | PCRc  + seq | *Babesia* spp.-*Theileria* spp.:  *- B.* *divergens*  - *B.* *venatorum* | **F: 5/259**  **F: 1/259**  **F: 4/259** |
|  | Reis et al. 2011 | Questing | PCRc  + seq | *Babesia* spp.-*Theileria* spp.:  *- B.* *venatorum* | **N (g. n = 10): 3/36**  **F: 1/69**  **M: 2/122** |
| Midi-Pyrénées,  Haute-Garonne | Lebert et al. 2020 | Questing | PCRn  + seq | *Babesia* spp.:  - *B.* *venatorum* | **N: 23/1,891** |
| Midi-Pyrénées,  Hautes-Pyrénées | Akl et al. 2019 | Questing | PCRc  + seq | *Babesia* spp.-*Theileria* spp.:  *- B.* *venatorum* | **N: 3/531**  **F: 0/86**  **M: 0/79** |
| Nord-Pas-De-Calais, Nord | Halos et al. 2005 | Questing | PCRc | *Babesia* spp. | **N: 9/74**  **F: 6/9**  **M: 4/9** |
| Pays-de-la-Loire, Loire-Atlantique | Cotté et al. 2010 | Questing | PCRc | *Babesia* spp. | **N: 4/267**  **F: 21/203**  **M: 10/102** |
| Poitou-Charente, Deux-Sèvres | Bonnet et al. 2013 | Questing | PCRc  + RLBH^b^ | *Babesia* spp. and *Theileria* spp. | F: 0/18  M: 0/12 |
| Rhône-Alpes,  Isère | Bonnet et al. 2013 | Questing | PCRc  + RLBH^b^ | *Babesia* spp. and *Theileria* spp. | F: 0/4  M: 0/2 |

Notes: “Region, department”: French region and department of sampling; “Reference”: source of the data; “Sampling”: sampling of the ticks: Attached: tick sampled on a host with host species in brackets (shaded), Questing: tick sampled while questing for a host; “Method”: PCR: Polymerisation Chain Reaction, PCRc: classical PCR followed by gel electrophoresis, PCRn: nested PCR, PCRq: quantitative PCR, PCRqm: microfluidic quantitative PCR, seq: amplicon sequencing, RLBH: identification by reverse line blot hybridization;

“*Babesia* spp.-*Theileria* spp.”: *Babesia* and/or *Theileria* species searched for; “Positive/tested”: number of positive ticks or pool of ticks on the number of tested ones: L: larvae, N: nymphs, F: adult females, M: adult males, A: adults of unspecified sex, ?: tick stage not specified, “g. n = x”: analyses of pooled DNA of several (x) ticks, /?: unknown number of tested ticks or group of ticks. Positive results are in bold.

^a^ Results of sequence analyses when different from expected by PCR are in brackets.

^b^ The following species have been searched for: *Babesia*/*Theileria* spp., *B.* *bigemina*, *B.* *bovis*, *B.* *divergens*, *B.* *major*, *B.* *motasi*, *B.* *ovis*, *B.* *crassa*, *Th*. *annulata*, *Th*. *velifera*, *Th*. *taurotrago*, *Th*. *mutans*, *Th*. *buffeli*/*orientalis*, *Th*. *ovis*, *Th*. *lestoquardi*, *Th*. spp. sp1 (China), *Th*. spp. sp2 (China), *B.* spp. sp1 (Turkey), and *B.* sp2 (Lintan).

^c^ Results respectively for salivary glands, gut, and both.

^d^ Species not searched for by PCRqm, but results obtained after checking by sequencing.

This is a supplementary material for “Perez, G., Bournez, L., Boulanger, N., Fite, J., Livoreil, B., McCoy, K. D., Quillery, E., René-Martellet, M., and Bonnet, S. I. The distribution, phenology, host range and pathogen prevalence of *Ixodes ricinus* in France: a systematic map and a narrative review”, a preprint recommended in PCI Infections <https://doi.org/10.24072/pci.infections.100076>.
