## Supplementary material for "The distribution, phenology, host range and pathogen prevalence of *Ixodes ricinus* in France: a systematic map and narrative review": SI 17

**Supplementary information 17: Tick-borne encephalitis virus prevalence in *Ixodes ricinus* ticks**

Prevalence of the tick-borne encephalitis virus (TBEv) in questing *Ixodes ricinus* ticks sampled in European France by tick stage.

| Region, department | Reference | Sampling | Method | TBEv sub-type | Positive ticks/Tested ticks |
| --- | --- | --- | --- | --- | --- |
| Alsace,  Bas-Rhin | Bestehorn et al. 2018 | Questing | RT-PCR  + seq | “European” | **N (g. n = 10): 1/? (954 ticks)**  F (g. n = 5): 0/? (378 ticks)  M (g n = 5): 0/? (392 ticks) |
|  | Chatelain & Ardoin 1978 | Questing | Mouse inoculation | “European” | **L&N: 3/11,068**  **F: 11/1,583**  **M: 3/1,396** |
|  | Pérez-Eid et al. 1992 | Questing | Mouse inoculation | “European” | **N (g. = 30-40): 4/251 (8,587 ticks)**  **F (g. n = 10-15): 20/? (2,975 ticks)**  **M (g. n = 15-20): 10/? (2,644 ticks)** |
|  |  | Attached (vertebrates) | Mouse inoculation | “European” | ? (g. = ?): 0/? (10,298 ticks) |
| Alsace,  Haut-Rhin | Bestehorn et al. 2018 | Questing | RT-PCR  + seq | “European” | N (g. n = 10): 0/? (363 ticks)  F (g n = 5): 0/? (34 ticks)  M (g n =  5): 0/? (23 ticks) |
|  | Bournez et al. 2020a | Questing | RT-PCRq  + RT-PCRn  + seq | “European” | **N (g. n = 1-5): 8/570 (7,070 ticks)**  F: 0/219  M: 0/199 |
|  |  | Attached (rodents) | RT-PCRq  + RT-PCRn  + seq | “European” | **L (g. n = 10): 3/? (283 ticks**)  N: 0/9  F: 0/4 |
|  | Gondard et al. 2018 | Questing | RT-PCRqm | “European” | N (g. n = 31): 0/45 (1,395 ticks) |
|  |  |  |  | “Far East” | N (g. n = 31): 0/45 (1,395 ticks) |
|  |  |  |  | “Siberian” | N (g. n = 31): 0/45 (1,395 ticks) |
|  | Moutailler et al. 2014 | Questing | Hseq | All viruses | N (g. n = 1,455): 0/1 |
| Champagne-Ardennes, Ardennes | Moutailler et al. 2014 | Questing | Hseq | All viruses | N (g. n = 285): 0/1  F (g. n = 268): 0/1  M (g. n = 228): 0/1 |
| Ile-de-France,  Essonne | Gondard et al. 2018 | Questing | RT-PCRmq | “European” | N (g. n = 31): 0/45 (1,395 ticks) |
|  |  |  |  | “Far East” | N (g. n = 31): 0/45 (1,395 ticks) |
|  |  |  |  | “Siberian” | N (g. n = 31): 0/45 (1,395 ticks) |
| Picardy,  Somme | Gondard et al. 2018 | Questing | RT-PCRmq | “European” | N (g. n = 31): 0/45 (1,395 ticks) |
|  |  |  |  | “Far East” | N (g. n = 31): 0/45 (1,395 ticks) |
|  |  |  |  | “Siberian” | N (g. n = 31): 0/45 (1,395 ticks) |

Notes: “Region, department”: French region and department of sampling; “Reference”: source of the data; “Sampling”: sampling of the ticks: Attached: tick sampled on a host with host species in brackets (shaded), Questing: tick sampled while questing for a host; “Method”: PCR: polymerisation chain reaction, RT-PCR: reverse transcriptase PCR, RT-PCRn: nested RT-PCR, RT-PCRq: quantitative RT-PCR, PCRqm: microfluidic RT-PCRq, Hseq: high-throughput sequencing, seq: amplicon sequencing; “TBEv sub-type”: sub-type of the virus searched for; “Positive ticks/Tested ticks” L: larvae, N: nymphs, F: adult females, M: adult males, A: adults of unspecified sex, L&N: larvae and nymphs, ?: tick stage not specified, “g. n = x”: analyses of pooled DNA of several (x) ticks, /?: unknown number of tested ticks or group of ticks. Positive results are in bold.

This is a supplementary material for “Perez, G., Bournez, L., Boulanger, N., Fite, J., Livoreil, B., McCoy, K. D., Quillery, E., René-Martellet, M., and Bonnet, S. I. The distribution, phenology, host range and pathogen prevalence of *Ixodes ricinus* in France: a systematic map and a narrative review”, a preprint recommended in PCI Infections <https://doi.org/10.24072/pci.infections.100076>.
