## Supplementary material for "The distribution, phenology, host range and pathogen prevalence of *Ixodes ricinus* in France: a systematic map and narrative review": version 1

**Keywords:** *Anaplasma;* *Babesia*; *Bartonella*; *Borrelia*; *Coxiella*; *Francisella*; *Theileria*; *Rickettsia*; tick-borne encephalitis virus; hard tick; tick-borne diseases.

### 1 Introduction

Among the forty or so tick species present in European France (i.e., excluding the ultra-marine territories, France hereafter), the hard tick *Ixodes ricinus* is the most frequently involved in human tick bites (Gilot & Marjolet 1982; Pérez-Eid 2007). This tick transmits several pathogens responsible for human and animal diseases (e.g. Lyme borreliosis, tick-borne encephalitis, piroplasmosis, anaplasmosis) (Heyman et al. 2010). Both the biology and ecology of this European tick species have been recently reviewed by Gray et al. (2021), and are known to vary greatly among regions. Data collected at national levels can provide insight into the factors responsible for at least some of this variation, but are rarely available to the general scientific community because of a mix of published and unpublished work and its production in different languages. Here, we synthesize the available data on *I. ricinus* and its associated pathogens in France up to 2020.

**Table 1:** Components elements of the literature search.

| Component | Elements |
| --- | --- |
| Population | Any *I*. *ricinus* populations at larval, nymphal or adult stage |
| Context | Temporal variations (daily, seasonal, annual); micro-habitat; habitat; landscape; climate; vertebrate host density |
| Comparator | Temporal comparisons: inter-annual, intra-annual, weather-related and daily; spatial comparisons according to climate (biome), land cover and land use (landscape context and management, habitats, vegetation types, anthropic activities, management); host density and diversity (wildlife management, ex-closure or enclosure, population fluctuation); potential treatments (e.g. acaricide treatments, vaccinations) |
| Outcome variables | Presence/absence; abundance; density; hosts parasitic burden (presence, prevalence or mean infestation); distribution and dispersal (displacements at different scales, gene flows); behaviour (e.g. questing activity, host preference); search for tick-borne pathogens (presence and/or prevalence) |

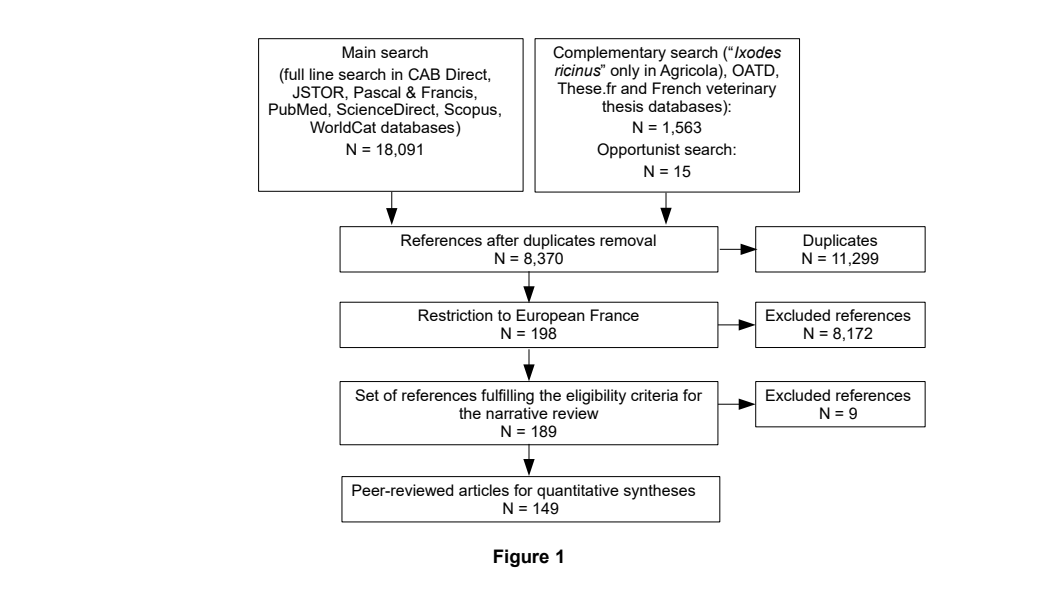

**Figure 1:** PRISMA-style scheme reporting the literature search and selection strategy arriving to the final 189 references including the 149 peer-reviewed articles used for data compilation.

##### 3.1.2 Description of eligible references

Most of the 189 references were peer-reviewed journal articles and included primary research articles (n = 137, 72.5%) and reviews (n = 12, 6.3%), whereas other documents were DVM and Doctor of Pharmacy (PharmD) dissertations (n = 24, 12.2%), PhD theses (n = 15, 8.5%), or book sections (n = 1, 0.5%). Two-thirds of the journal articles were in English (n = 97, 65.1%), while others were in French (n = 52, 34.9%). The oldest reference recorded was published in 1965. References were scarce until the 1990’s, slowed-down again in the 2000’s and then increased again from 2010 onwards (**SI 6**). The 12 selected reviews provided only a broad overview of ticks and tick-borne diseases or, conversely, a synthesis on some precise aspect of *I*. *ricinus* ecology and tick-borne diseases in France; none contained a systematic map providing a summary of existing data for the species.

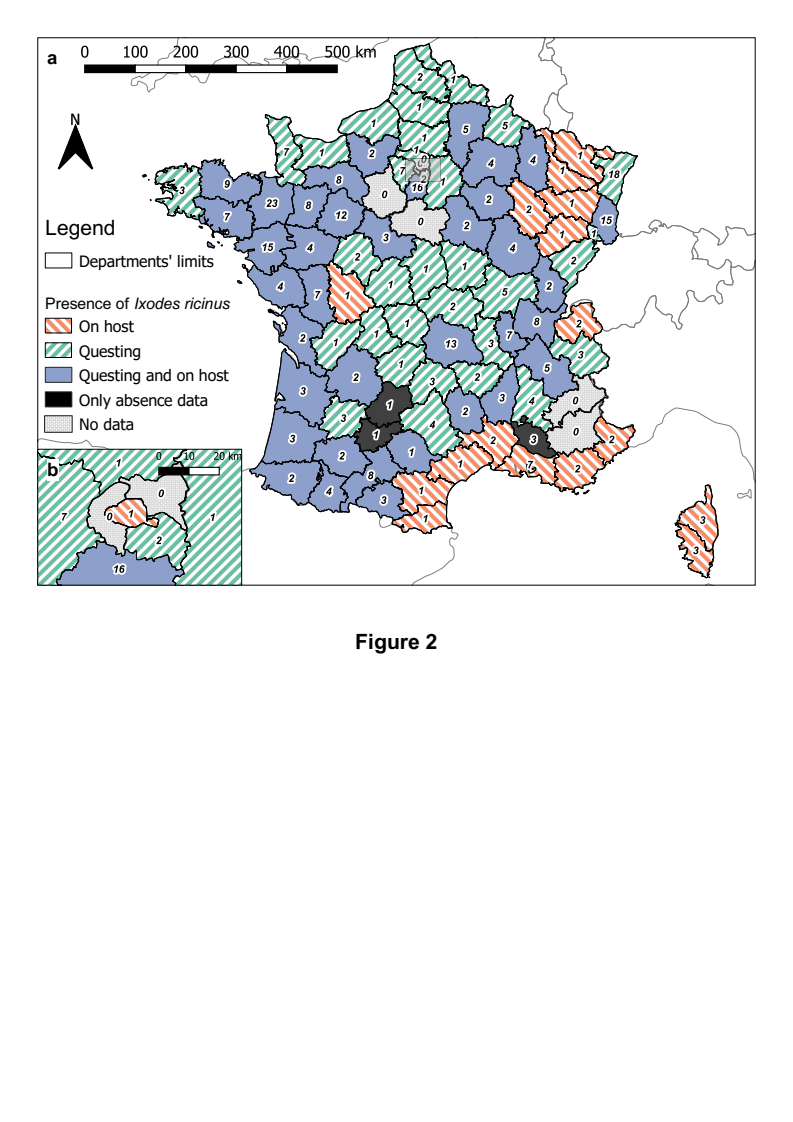

**Figure 2:** Distribution map of *Ixodes ricinus* in European France by departments according to the final 189 references, and whether ticks were questing or attached on host. Refer to **SI 5** for the name of the departments.

In south-western France, in a study that aimed at detecting the presence of *I*. *ricinus* on 100 sites over 15 different vegetation types covering 23 departments, the species was never detected in sites with Mediterranean vegetation (Doche et al. 1993; Gilot et al. 1995). The species was therefore considered absent from the Mediterranean climatic region. However, in Corsica, Grech-Angelini et al. (2016) recently reported the species feeding on cattle in areas above 600 m above sea level (a.s.l.). Thus, *I. ricinus* ticks seem to be present in some Mediterranean departments, but only under specific environmental conditions and above a certain altitude (Stachurski & Vial 2018). An inventory in the Rhône valley (south-eastern France) of a 330×80 km area recorded the presence of *I. ricinus* from 200 to 1,150 m a.s.l. (Gilot et al. 1989). More recently, *I. ricinus* was collected at the top of the Pic de Bazès, in the eastern part of the French Pyrenees, at 1,800 m a.s.l. (Akl et al. 2019; Bourgoin et al. 2012). These data support other observations in Europe that have shown that its presence is increasingly detected at higher altitudes, beyond the previously established 1,500 m (Gern, Morán Cadenas & Burri 2008; Danielova et al. 2006; Gilbert 2010; Martello et al. 2014; Garcia-Vozmediano et al. 2020). *I.* *ricinus* was also considered to be scarce on the Atlantic coast because of coastal environmental conditions, like wind and spray which can be highly desiccative (Degeilh et al. 1994). This tick is nonetheless present on Belle-Île-en-Mer island (Bonnet et al. 2007), indicating that the species can be present on islands, but at low abundance. Indeed, ten years after the first study, Michelet et al. (2016) did not collect any *I*. *ricinus* ticks on this same island.

#### 3.3 Tick-borne pathogens in *Ixodes ricinus* ticks of France

Different tick-borne pathogens that have been examined in *I*. *ricinus* ticks in France and are briefly presented below (see McCoy & Boulanger for more detail). We summarised existing data from across studies and present this data in detail in the Supplementary Materials (**SI 9-17**). Not enough data exist to establish prevalence maps for most pathogens (with the exception of *Borrelia burgdorferi* s.l.), but presence/absence information is illustrated (**Figures 3-10**).

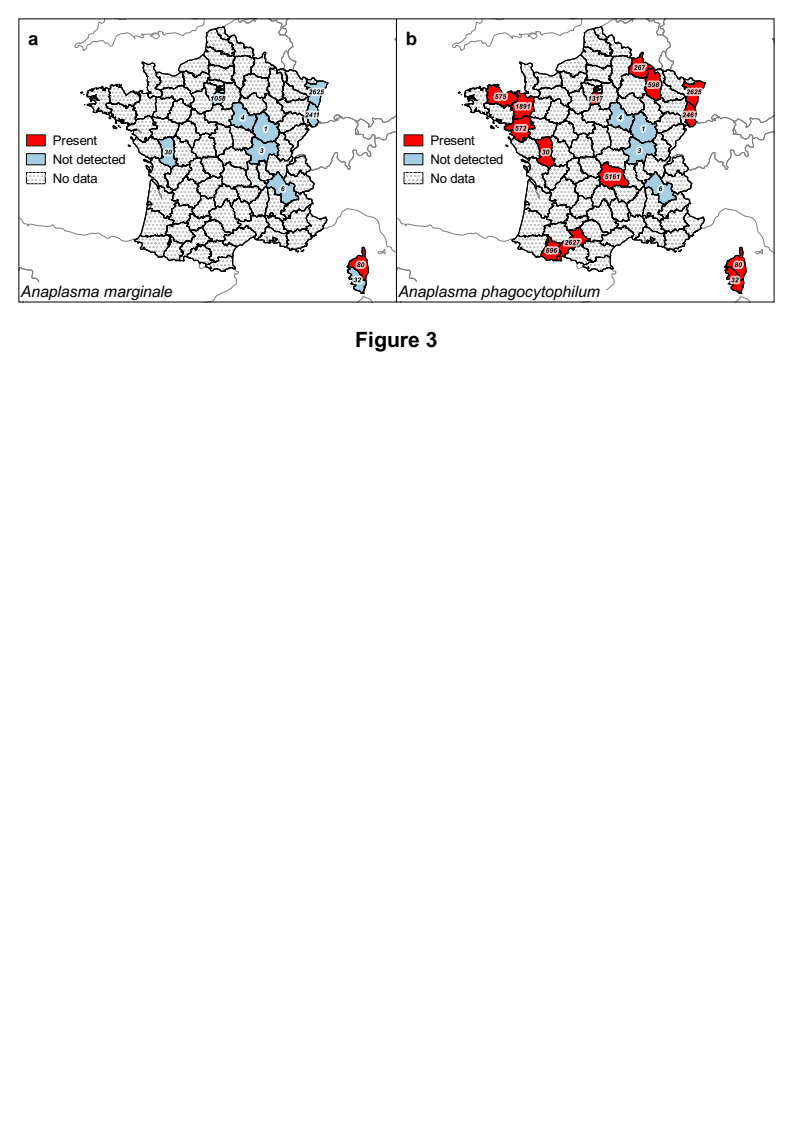

**Figure 3:** Presence maps of *Anaplasma marginale* and *A*. *phagocytophilum* in questing and attached nymph and adult *Ixodes ricinus* ticks in European France by department.

Detected presence by PCR methods and respective sampling effort per department in number of tested ticks (nymphs and adults) for *A*. *marginale* (**a**) and *A*. *phagocytophilum* (**b**).

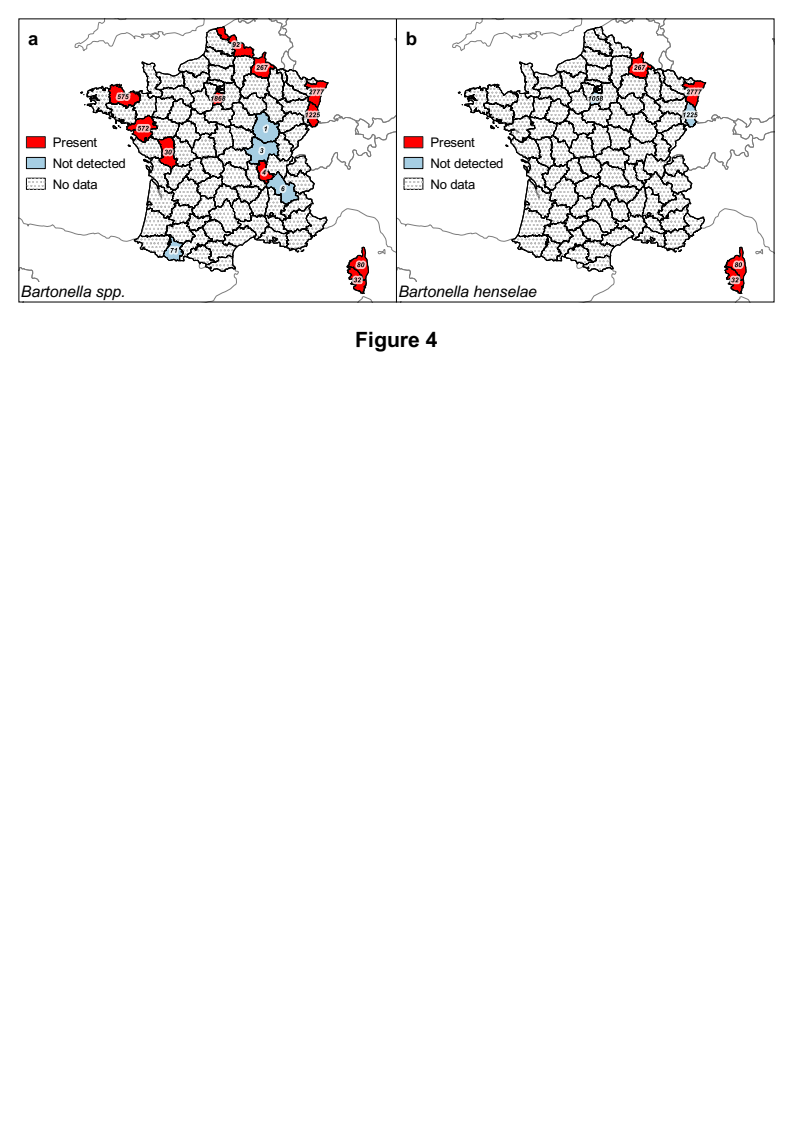

**Figure 4:** Presence maps of *Bartonella* spp. and *B*. *henselae* alone in questing and attached nymph and adult *Ixodes ricinus* ticks in European France by department.

Detected presence by PCR methods and respective sampling effort per department in number of tested ticks (nymphs and adults) for *Bartonella* spp. (**a**) and *B*. *henselae* alone (**b**).

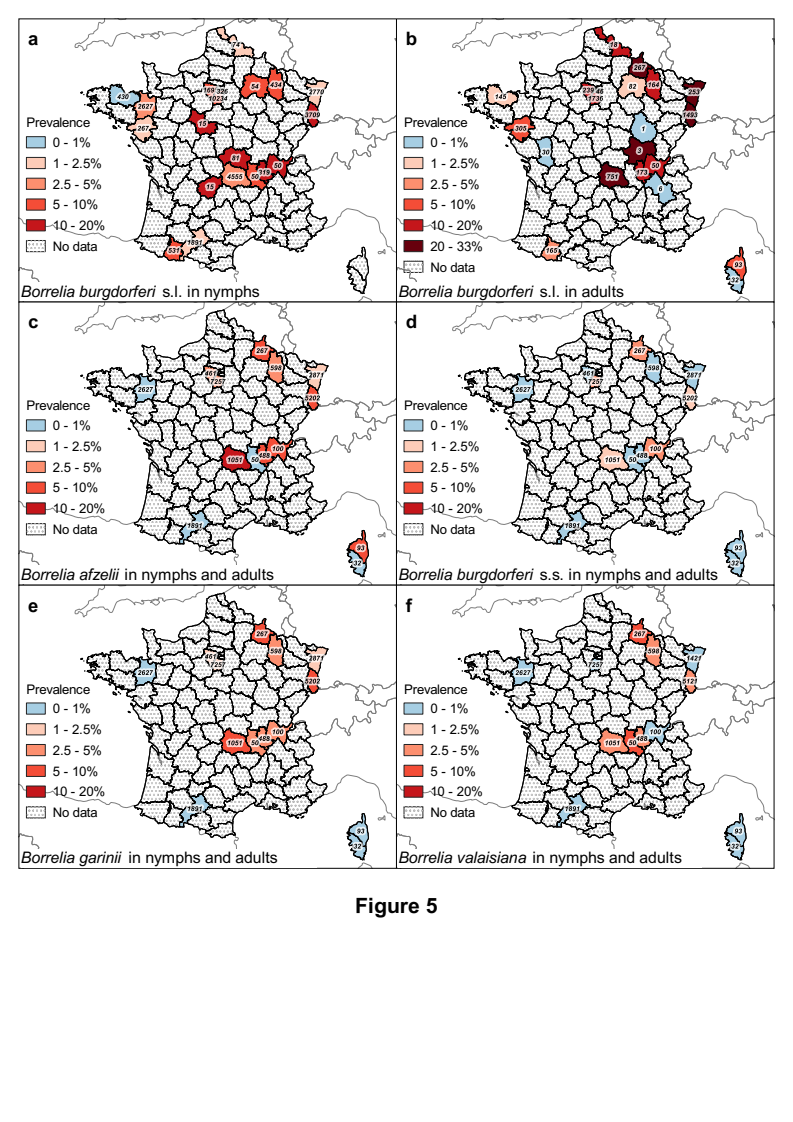

**Figure 5:** Prevalence maps of *Borrelia burgdorferi* sensu lato and *B*. *afzelii*, *B*. *burgdorferi* sensu stricto (s.s.), *B*. *garinii*, and *B*. *valaisiana* in questing and attached nymph and adult *Ixodes ricinus* ticks in European France by department.

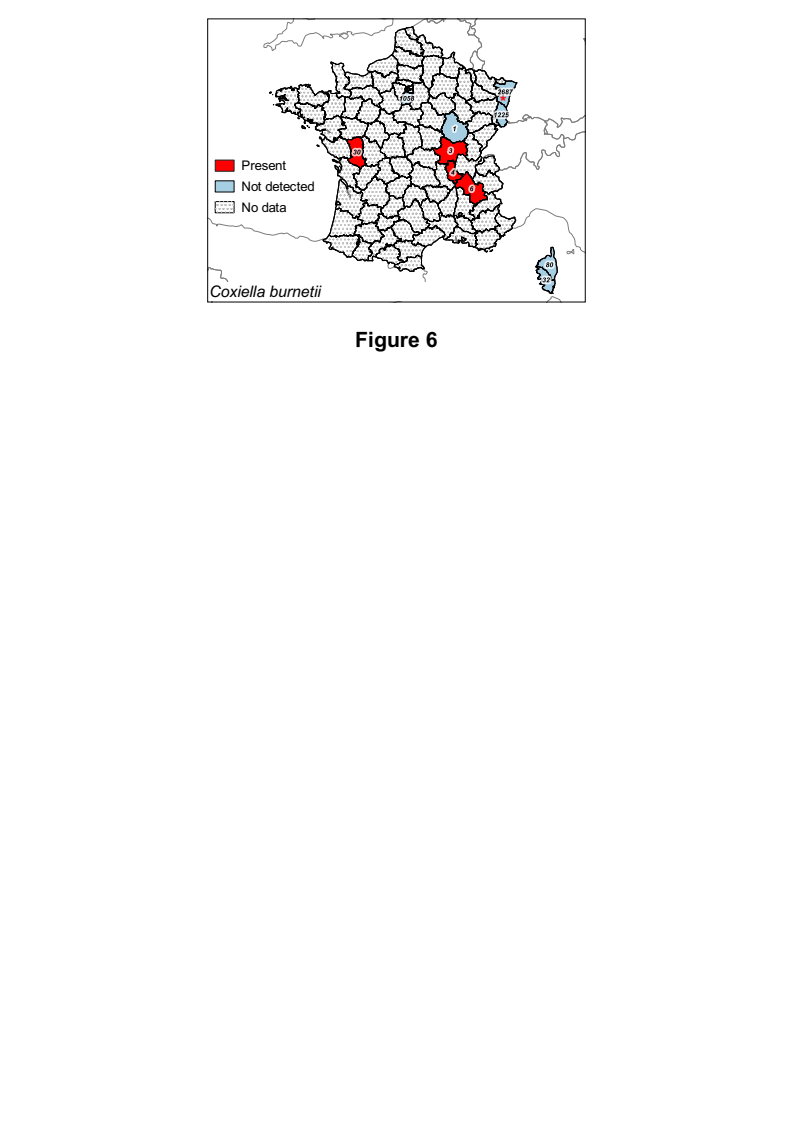

**Figure 6:** Presence map of *Coxiella burnetii* in questing and attached nymph and adult *Ixodes ricinus* ticks in European France by department.

Detected presence by PCR methods and respective sampling effort per department in number of tested ticks (nymphs and adults) for *C*. *burnetii*.

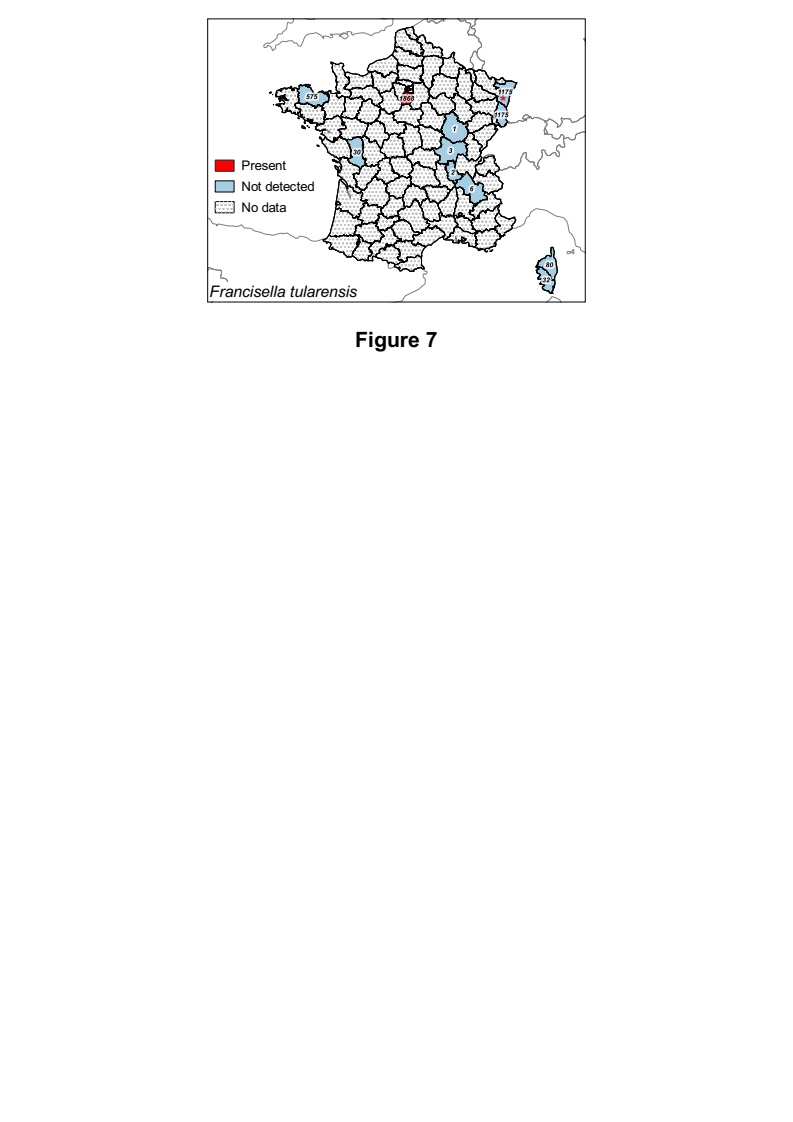

**Figure 7:** Presence map *Francisella tularensis* in questing and attached nymph and adult *Ixodes ricinus* ticks in European France by department.

Detected presence by PCR methods and respective sampling effort per department in number of tested ticks (nymphs and adults) for *F*. *tularensis*. * Detection of unidentified *Fransicella* sp..

##### 3.3.7 The genus *Rickettsia*

The genus *Rickettsia* includes more than thirty intracellular bacterial species. The pathogenic species are separated into two groups: the “typhus” group (TG), with two insect-transmitted species (*R*. *typhi* transmitted by fleas and *R*. *prowazekii* transmitted by lice) and the “spotted fever” group (SFG), with most species transmitted by hard ticks (except for *R.* *felis* transmitted by fleas and *R*. *akari* by mites). Many other *Rickettsiae* are endosymbionts involved in various metabolic functions in their hosts (Abdad et al. 2018; Bonnet & Pollet 2021). For the pathogenic *Rickettsiae,* several vertebrate species could play the role of reservoir. Species of the SFG are more or less pathogenic in humans and responsible for rickettsioses, characterised by an eschar at the bite site and/or fever and headache, and sometimes with complications like cutaneous necrosis, pneumonia, endothelial damages and organ failure (Abdad et al. 2018).

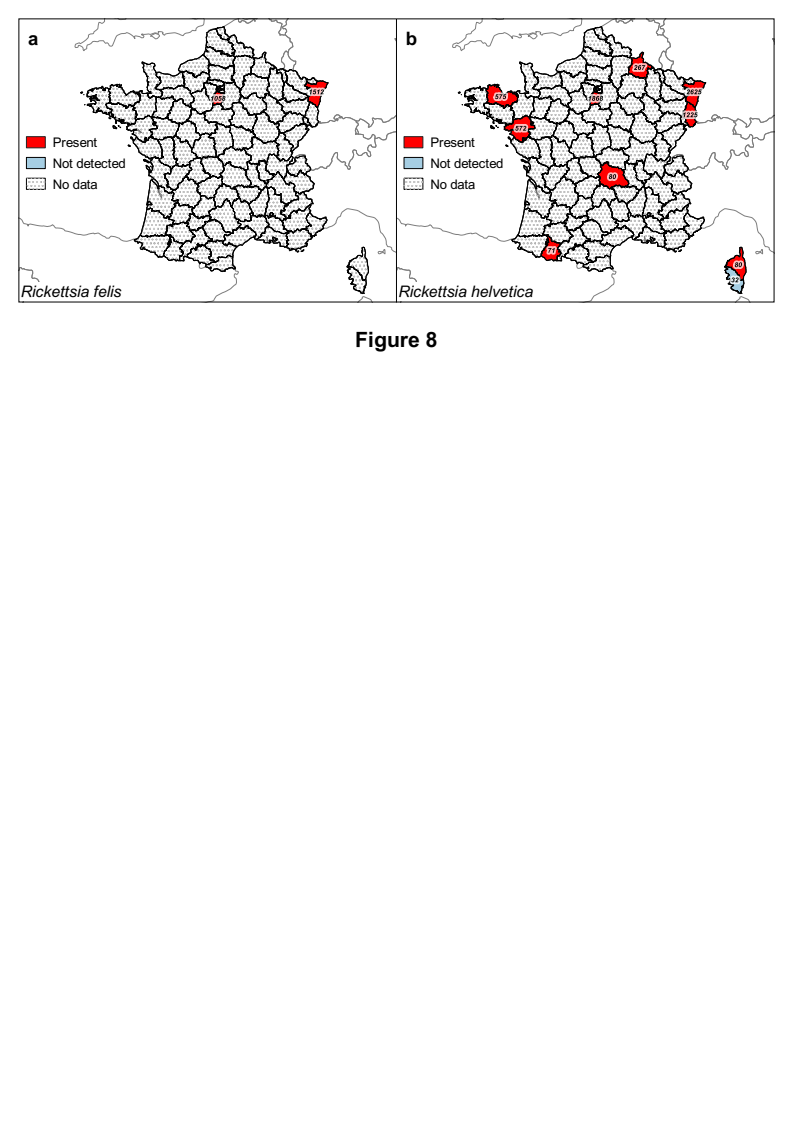

**Figure 8:** Presence maps of *Rickettsia felis* and *R*. *helvetica* in questing and attached nymph and adult *Ixodes ricinus* ticks in European France by department.

Detected presence by PCR methods and respective sampling effort per department in number of tested ticks (nymphs and adults) for *R*. *felis* (**a**) and *R*. *helvetica* (**b**).

In France, the *Rickettsia* species detected in questing *I*. *ricinus* ticks, when identified at the species level, are *R*. *felis*, *R*. *helvetica* and *R*. *monacensis* (**SI 15**). *R*. *helvetica* DNA was found in most regions examined, suggesting a distribution throughout France (**Figure 8**) (Parola et al. 1998b; Davoust et al. 2012; Michelet et al. 2014; Moutailler et al. 2016b; Bonnet et al. 2017). When detected, observed prevalence was estimated between 1.1 and 14.3% in nymphs and between 1.6 and 25.0% in adults (Cotté et al. 2010; Michelet et al. 2014; Nebbak et al. 2019; Lejal et al. 2019a). Data are too scarce to allow valid comparison of the prevalence between geographical regions, but it seems higher in forested departments. DNA of *R*. *felis* was detected in questing ticks from the Paris region (Essonne) and from north-eastern (Alsace) France (**Figure 8a**), whereas *R*. *monacensis* DNA was found in southern France (Vayssier-Taussat et al. 2013; Akl et al. 2019; Lejal et al. 2019a; Lejal et al. 2019b). Tests to detect DNA of *R*. *conorii*, *R*. *slovaka*, and *R*. *massiliae* were carried out on a large number of questing adult and nymphal ticks in northern-eastern France, but none were found positive (Michelet et al. 2014).

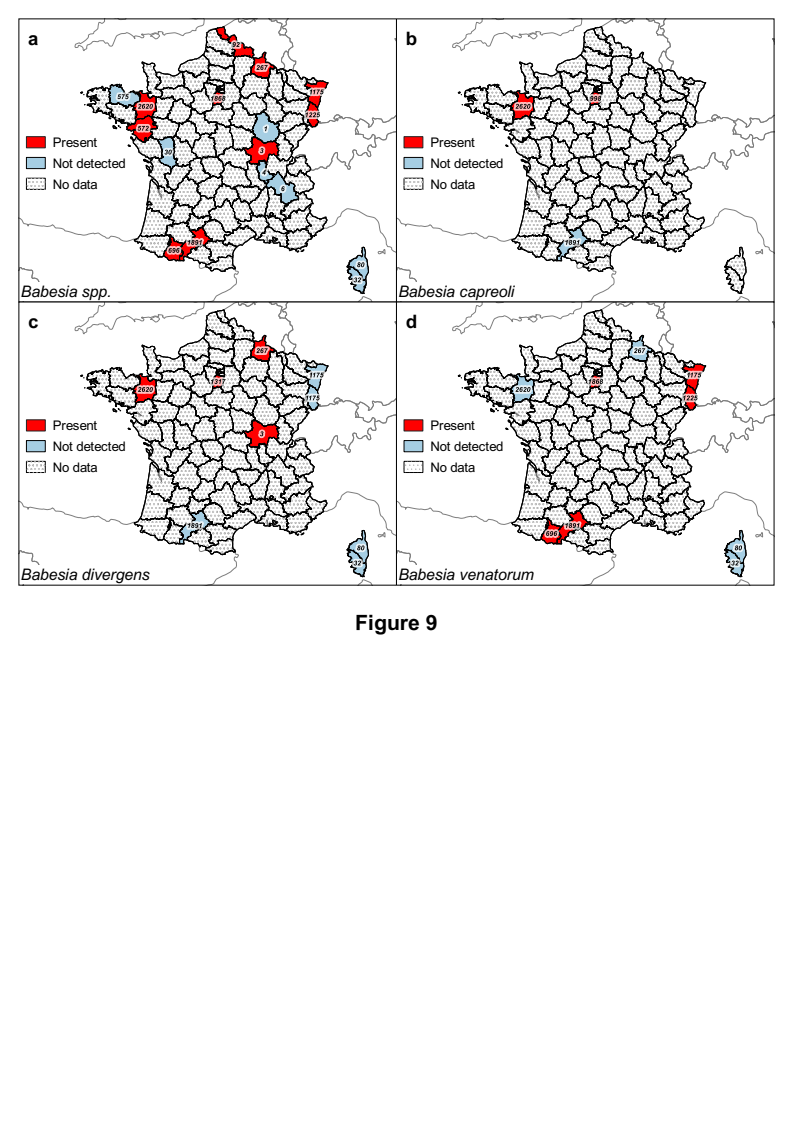

**Figure 9:** Presence maps of all *Babesia* spp., *B*. *capreoli*, *B*. *divergens* and *B*. *venatorum* in questing and attached nymph and adult *Ixodes ricinus* ticks in European France by department.

Detected presence maps and respective sampling effort per department in number of tested ticks (nymphs and adults) for *Babesia* spp. (**a**), *B*. *capreoli* (**b**), *B*. *divergens* (**c**) and *B*. *venatorum* (**d)**.

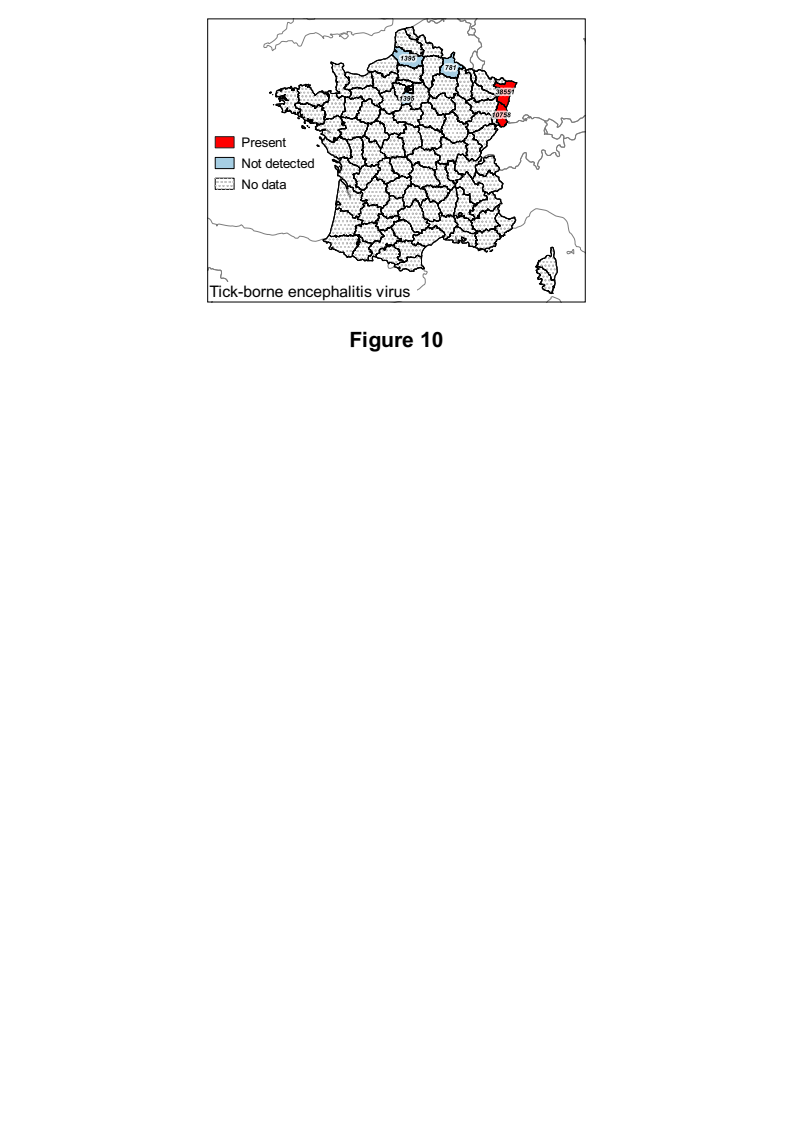

**Figure 10:** Presence map of tick-borne encephalitis virus (TBEV) in questing and attached nymph and adult *Ixodes ricinus* ticks in European France by department.

Detected presence of TBEV by RT-PCR methods and sampling effort per department in number of tested ticks (nymphs and adults).
